# Directed evolution of nanosensors for the detection of mycotoxins

**DOI:** 10.1101/2023.06.13.544576

**Authors:** Benjamin P. Lambert, Afsaneh Taheri, Shang-Jung Wu, Alice J. Gillen, Mahdi Kashaninejad, Ardemis A. Boghossian

## Abstract

In this study, we develop and apply a directed evolution approach to engineer the optical sensing properties of DNA-wrapped single-walled carbon nanotubes (DNA-SWCNTs) towards mycotoxins, a class of molecules critical to detect in the food industry. We successfully demonstrate the creation of sensors for the detection of both the aflatoxin B1 (AFB1) and fumonisin B1 (FB1) mycotoxins based on the specific response of the (9,4) and (7,5) SWCNT chirality fluorescence peaks, respectively. The resulting chirality-specific responsivity was used to demonstrate the multimodal detection of both mycotoxins at different wavelengths of light in the presence of complex food medium. Moreover, we show that directed evolution can be used not only to improve the chiral-dependent selectivity of our sensors to the mycotoxins, but also the sensor sensitivity and fluorescence intensity through multiple rounds of evolution. The approach demonstrated in this study is versatile and could be generalized to other SWCNT sensors as well as other nanosensors comprising a biological element.

---

Nanosensors, sensors with at least one dimension below 100 nm, have garnered significant interest in a variety of fields including health care^1^, environmental monitoring^2^, and the food industry^3^. However, despite their many advantages, the interaction and performance of such sensors in the presence of a desired target analyte can be difficult to predict, and therefore difficult to engineer, because of uncertainties surrounding the sensor’s structure-function relationship. In such cases, the sensors are typically engineered using empirical approaches^4–7^ based on extensive screening of random configurations yielding sensors with suboptimal performances. Analogous challenges in engineering complexes with ill-defined structure-function relationships have already been addressed in the field of protein engineering, where synthetic biologists have relied on directed evolution to engineer proteins in a guided manner^8^. With this approach, a continuous relationship between protein’s structure and function is assumed, limiting the screening to variants of an initial protein with residual, albeit sub-optimal, performance. This strategy therefore biases the screening to variants that are likely to show at least some desired activity to enable the relatively efficient identification of optimized protein mutants that would, otherwise, be almost impossible to find through random screening^9^.

Recently, the applicability of directed evolution has been demonstrated beyond the field of protein engineering for DNA-wrapped single-walled carbon nanotube (DNA-SWCNT) nanosensors^10^. Semiconducting SWCNTs are particularly attractive materials for the creation of optical sensors owing to their sensitive and photostable near-infrared (NIR) fluorescence, as well as their ability to be functionalized with a wide variety of molecules^11^. When non-covalently conjugated with single-stranded DNA, SWCNTs can exhibit various sensing capabilities where the sensitivity and selectivity depend on the DNA sequence^5–7^. So far, how the DNA sequence and SWCNT chirality determines the sensing properties of DNA-SWCNT complexes remains unknown, limiting the performance and applicability of such sensors. In previous work, we demonstrated the application of directed evolution to improve the fluorescence quantum yield of DNA-SWCNT complexes by up to 56% in the absence of information on the complex’s structure. Herein, we use directed evolution to engineer the properties of DNA-SWCNTs beyond fluorescence intensity. Specifically, we demonstrate the first DNA-SWCNT sensors for the detection of mycotoxins and show how directed evolution can be used to engineer their performance.

Mycotoxins are secondary metabolites produced by fungi in food products that are responsible for a range of diseases in humans^12,13^. The Food and Agriculture Organization estimates that at least 25% of the global food crop is contaminated with mycotoxins^14^; aflatoxin B1 (AFB1), fumonisin B1 (FB1), ochratoxin A (OTA), zearalenone (ZEN), and deoxynivalenol (DON) are the most commonly occurring toxins in cereals such as corn^15^. AFB1 is of particular concern owing to its high prevalence and toxicity. The rapid, sensitive, and specific detection of mycotoxins is therefore crucial to increase food security and avoid economic losses associated with contamination. Conventional methods for assaying mycotoxins in food sources include chromatographic and immunoassay-based approaches. However, these methods tend to be time consuming, expensive, non-reversible or require extensive sample preparation^12,13^.

In this study, we propose a label-free approach for the rapid multimodal detection of mycotoxins. As this approach is based on the NIR fluorescence of SWCNTs, it enables a non-destructive, *in situ* detection of mycotoxin, while simultaneously reducing the optical background from the food matrix at visible wavelengths^16^. Although the initial DNA-SWCNT sensors found by random screening exhibited weak responses towards mycotoxins, we were able to engineer sensors towards two mycotoxins, AFB1 and FB1, by monitoring two different SWCNT chiralities. We further demonstrated that we could use directed evolution to enhance the performance of these sensors, increasing the AFB1 response by more than 3-fold. Moreover, we demonstrate the accelerated engineering of mutants with enhanced performances using a DNA shuffling recombination technique. Finally, our studies enabled us to elucidate two distinct modes of interaction for the AFB1 and the FB1 sensors, allowing a better understanding of the sensors’ underlying properties such as sensor reversibility. This work therefore demonstrates the strength of directed evolution for engineering the overall properties of DNA-SWCNT sensors as well as nanosensors in general that comprise a biological component.

## Results and discussion

We studied the response of DNA-SWCNT complexes towards a range of mycotoxins commonly found in corn-related products (**Figure 1a**): AFB1, FB1, OTA, ZEN and DON. Owing to its prevalence in the food industry, we first searched for promising DNA sequences for the detection of AFB1. We initially screened a library of 100 diverse DNA-SWCNT complexes (see **Methods**) against AFB1 and monitored the fluorescence response of the SWCNTs (**Figure S7**). From this screening, we identified an analogue of the (AG)_15_ sequence, Δ(AG), that exhibited a strong selective response towards AFB1 (**Figure 1b**). Specifically, we observed a red-shift of the SWCNT fluorescence emission towards higher wavelengths for the (9,4) chirality (1.65 ± 0.16 nm). Although we observed strong shifting response, the intensity of the (9,4) peak post-addition (**Figure S8**) was not significantly different from the control.

**Figure 1:**
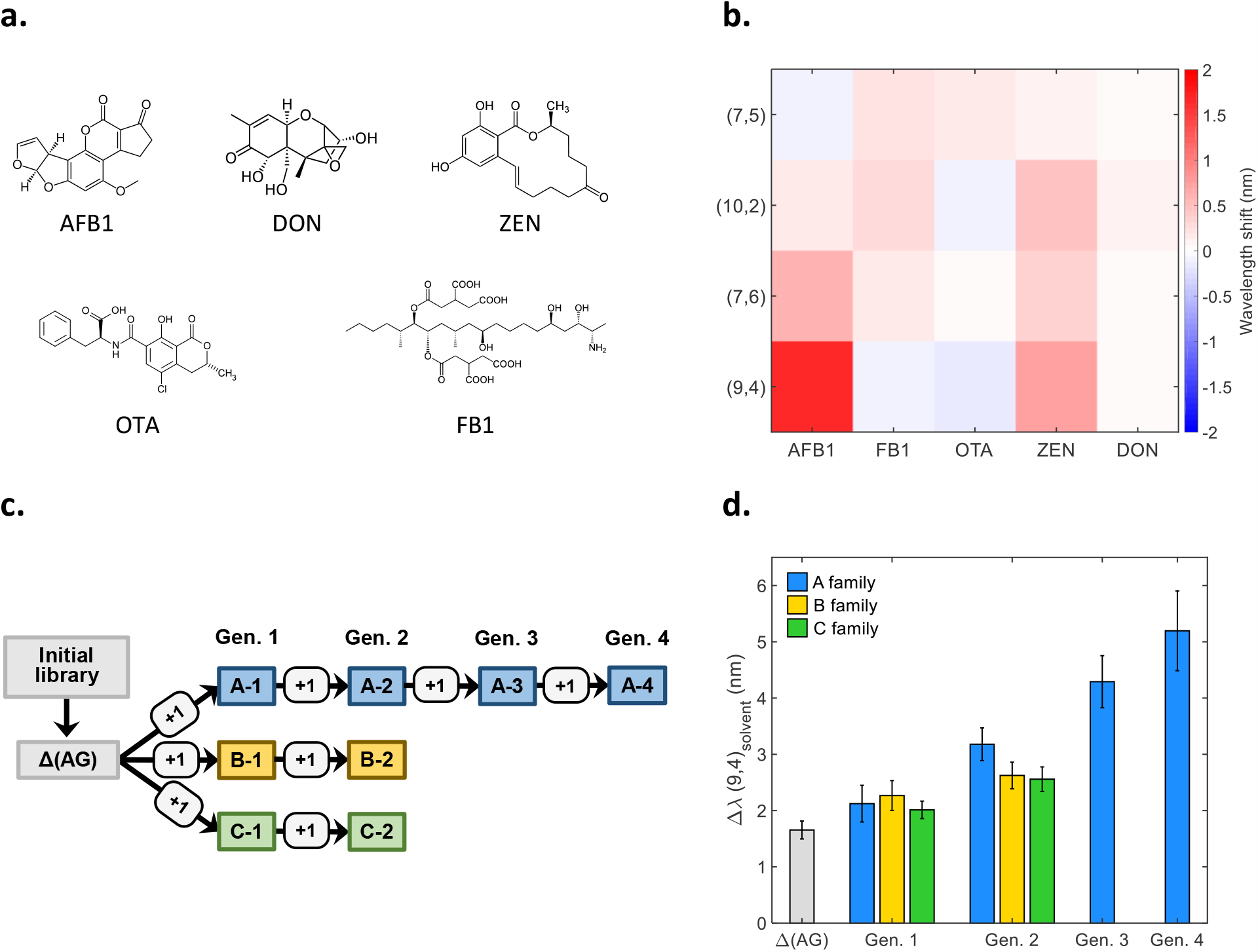
Directed evolution of the AFB1 sensor. (a) Structures of the mycotoxins screened: AFB1, DON, ZEN, OTA, FB1. (b) Fluorescence wavelength shifting response of the Δ(AG) sensor following the addition of different toxin solutions (10 μM). Wavelength shifts were calculated for the (7,5), (10,2), (7,6) and (9,4) SWCNT chiralities as the difference in peak position with and without toxin. (c) Schematic of the evolution of the AFB1 sensor. The sequences were selected based on increases in the shifting response of the (9,4) peak in the presence of AFB1. The Δ(AG) sensor was selected from an initial library (light grey). Three families of mutants were created (blue, yellow and green for the A, B, and C families, respectively) from individual mutants harboring a distinct random substitution in the Δ(AG) sequence. The “+1” denotes the introduction of a single, random base substitution. The numbers following the family letters correspond to the generation of the mutant. (d) Response of the AFB1 sensors across multiple rounds of evolution. The colors correspond to the families of sensors shown in (c). All measurements were taken in the presence of AFB1 (10 μM) following 200 min incubation. Error bars represent 1 σ standard deviation (n = 4).

We then applied the directed evolution approach on the Δ(AG) sequence to increase the shifting response of the (9,4) chirality towards AFB1 (**Figure 1c-d**). We sequentially introduced random base substitutions, one per round of evolution, and selected the mutants that showed greater fluorescence shifts in response to AFB1. The sequences of the main DNA mutants are listed in **Table S2**. From the first library of mutants originating from the Δ(AG) sequence, we selected three sensors, A-1, B-1 and C-1, which showed the greatest increase in response (**Figure 1d, Table S3**). These mutants were subsequently subjected to another round of mutagenesis, resulting in the selection of one additional mutant per sequence family (A-2, B-2 and C-2). As the A family of sensors demonstrated the greatest improvement of all three mutant families, we performed two additional rounds of mutagenesis resulting in the discovery of the A-3 and A-4 mutants (**Figure 1d, Table S3**). Using our directed evolution approach, we managed to successfully improve the shifting response of the sensors by 314% compared to the starting sequence after four rounds of evolution (**Figure 1d, Table S3**). When considering the selective engineering of DNA-SWCNT sensors, the increase demonstrated here by directed evolution is greater than what was previously shown for serotonin sensors through high-throughput selection from an even larger library size^17^. Moreover, this improvement was achieved without compromising the selectivity of the response (**Figure S9**). This systematic improvement of a single sensing property is unlikely in screens of random DNA-SWCNT sensors, which can also result in drastic differences across multiple properties, e.g. analyte sensitivity and selectivity^6,18^.

Although this approach has been shown to yield mutants with improved performances during the initial rounds of evolution, further improvements may eventually stagnate, in agreement with previous observations on the directed evolution of proteins. According to protein engineering theory^9^, the likelihood of identifying a sequence with improved performance is expected to decrease after several evolution rounds, particularly as one approaches a local optimum. To circumvent this limitation, one approach is to screen a larger library size to increase the search space within the vicinity of the local optimum. This approach increases the probability of including increasingly rare mutants with performances that exceed the already improved performance of the previous mutant. Another approach is to create a new library based on the combinatorial shuffling of mutations and fragments from the mutants from the previous rounds of evolution^19^. This approach may benefit from not only synergistic coupling of beneficial mutations that may lead to enhancements beyond those achieved with the original mutants, but also the ability to navigate the sequence space towards a new local optimum with higher performance.

Applying an analogous approach to our engineering strategy, we implemented a DNA shuffling approach to recombine the fragments of mutants from the previous evolution rounds^19^. Specifically, we selected the two best sensors from the second round of mutagenesis, A-2 and B-2, combined them using DNA shuffling and screened the resulting mutants for an improved AFB1 response (**Figure 2a, Figure S11**). From this screening, we identified the S5 sensor, using a fragment size of 5 nucleotides, that exhibited the highest increase in response compared to its parent sequences (up to 90% higher, **Figure 2b**). This shuffled sequence also demonstrated a significantly higher response compared to the A-4 sequence (**Figure 2b**) that was achieved after 4 rounds of evolution. This observation confirmed that the DNA shuffling strategy is an effective means of further engineering DNA-SWCNT sensors.

**Figure 2:**
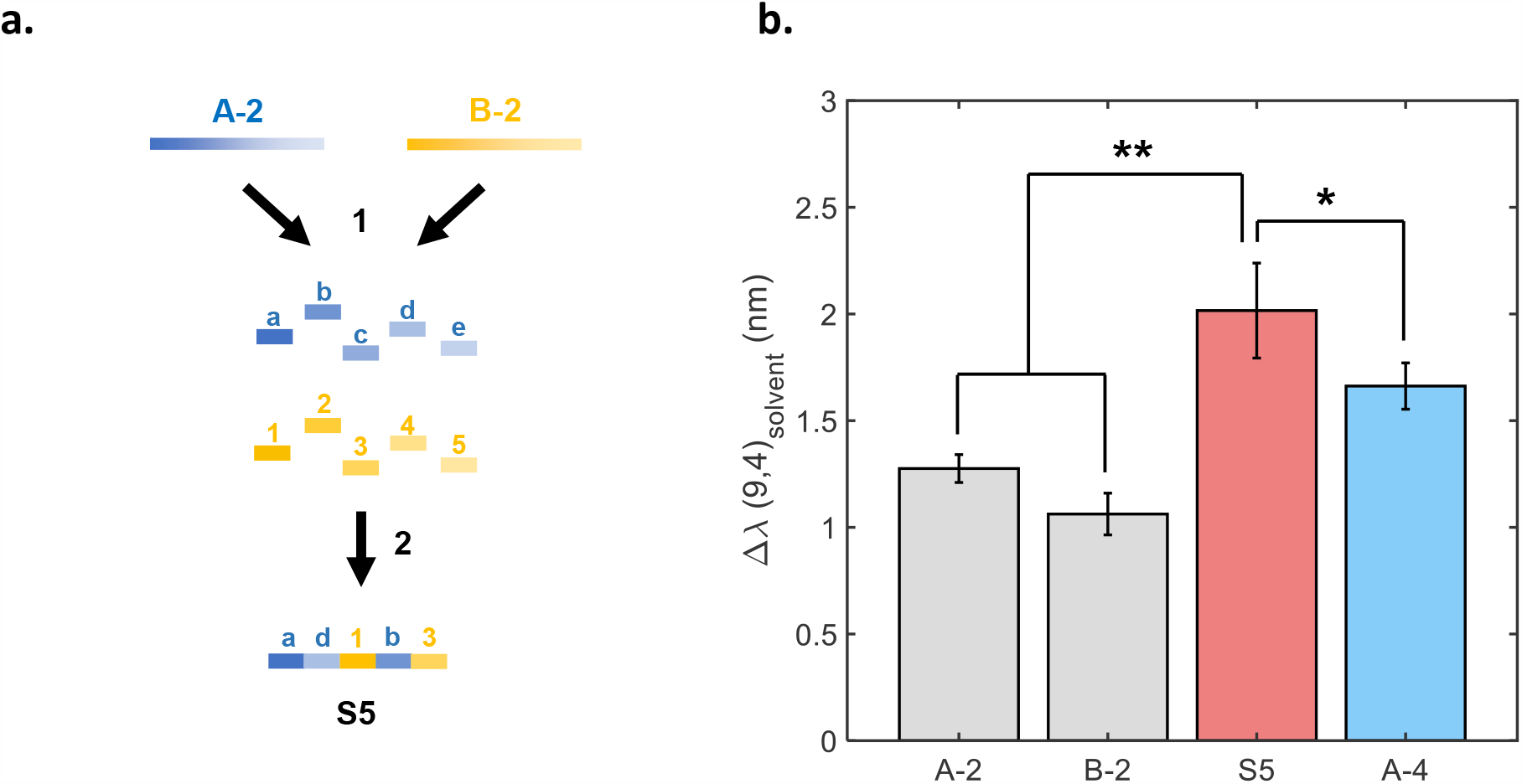
Improving sensor performance using DNA sequence shuffling and recombination. (a) Schematic of the methodology used to create shuffled sequences. The parent sequences, A-2 (blue) and B-2 (yellow), were decomposed into small fragments of fixed length, which were subsequently randomly recombined to form a library of shuffled sequences, including the S5 chimera sequence. (b) AFB1 response for the parent sensors (A-2 and B-2), the resulting shuffled sensor (S5) and the sensor selected after 4 rounds of mutagenesis (A-4). Measurements were taken in the presence of 2.5 μM AFB1 following 200 min incubation. * p < 0.05 and ** p < 0.01 (two-sample t-test, n = 4). Error bars represent 1 σ standard deviation (n = 4).

Despite the improved response of our DNA-SWCNT sensors towards AFB1, we observed a simultaneous decrease in the fluorescence intensity of the sensors with progressing rounds of evolution (**Figure 3a**), analogous to the decrease in stability observed in protein engineering^20^. For example, the intensity of the S5 sensor decreased by almost 50% compared to the Δ(AG) sensor. As the performance of fluorescence sensors is often linked to their brightness, this decrease could be detrimental to the applicability of the sensor. To address this issue, we applied an additional three rounds of mutagenesis on the S5 sensor to improve the fluorescence intensity (**Figure S12**). The final Q-3 sensor showed an increase in the fluorescence intensity of 21% versus the S5 sensor, without significant changes to the AFB1 response (**Figure 3b**). Furthermore, the Q-3 sensor showed a comparable intensity to the A-2 and B-2 sensors, while exhibiting a greater shifting response (+35% and +58%, respectively).

**Figure 3:**
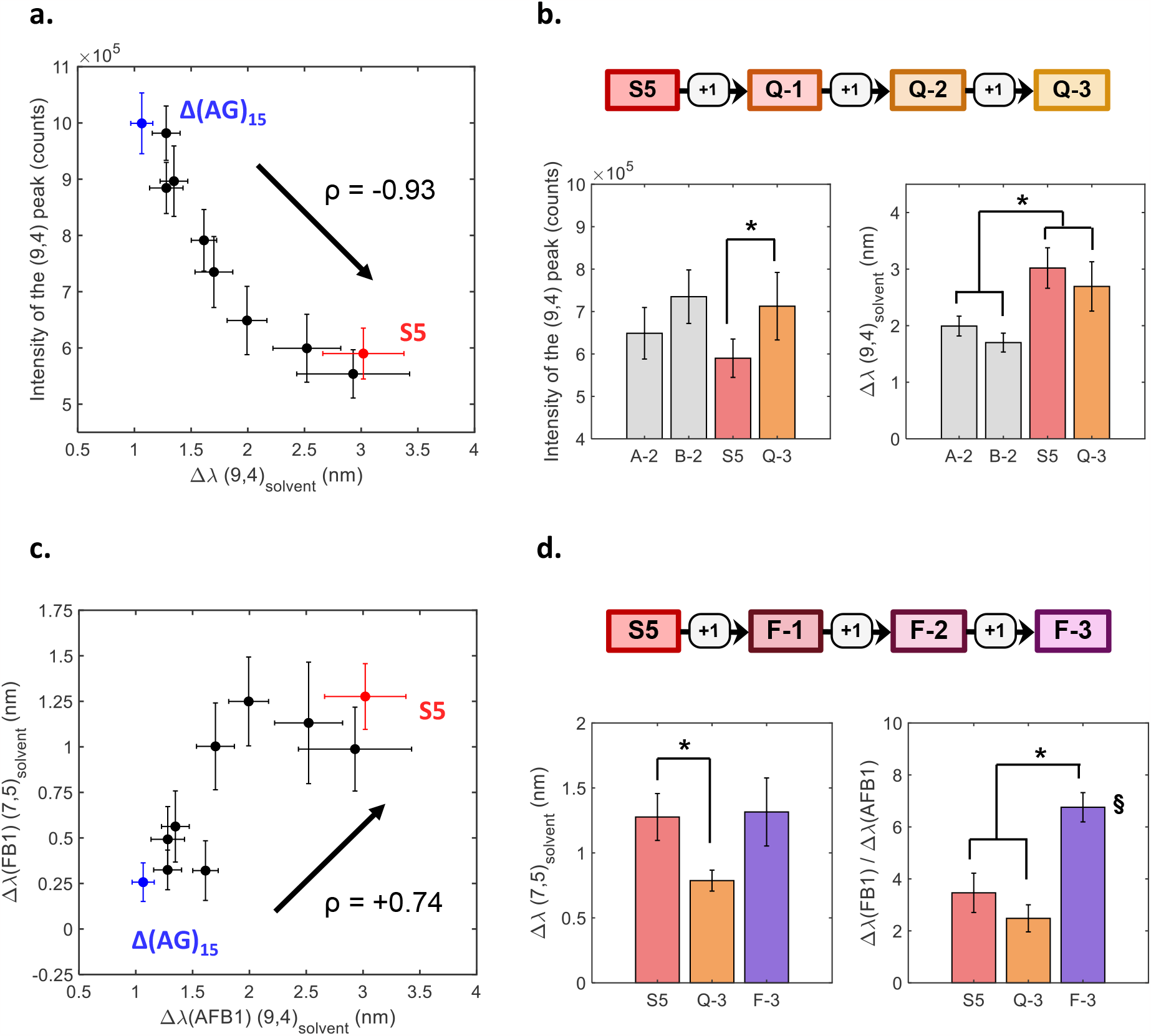
Directed evolution of the sensor’s intensity and selectivity. (a, c) Correlation between (a) the fluorescence intensity and the (9,4) response to AFB1 and between (c) the (7,5) response to FB1 and the (9,4) response to AFB1 for the selected mutants (Fig. 1c), the Δ(AG) and the S5 sensors. The increase in AFB1 response is accompanied by (a) a decrease in peak intensity (Spearman: ρ = - 0.95, p < 0.05) and (c) an increase in FB1 response (Spearman: ρ = 0.81, p < 0.05). (b) Comparison of the fluorescence intensity (left) and AFB1 response (right) between the A-2, B-2, S5 sensors and the Q-3 evolved for intensity (top). (d) Comparison of the FB1 response (left) and selectivity (right) between the S5, Q-3 sensors and the F-3 evolved for FB1 sensing (top). The selectivity is calculated as the ratio of the FB1 response to the AFB1 response for the (7,5) chirality. All measurements were taken in the presence of 5 μM of the corresponding toxin following 200 min incubation. The * indicates a significant difference between the means (p < 0.05, two-sample t-test, n = 4). All error bars represent 1 σ standard deviation (n = 4, n=3 if marked by §).

While improving the sensors for AFB1 detection via the (9,4) chirality, we noted a concurrent improvement in the sensors’ wavelength shifting response towards FB1 for the (7,5) and (10,2) chiralities (**Figure 3c, Figure S14**). These chiralities did not exhibit a significant change in intensity in the presence of FB1, once more yielding sensors based on wavelength shifting responses (**Figure S8, Table S4**). We performed three subsequent rounds of mutagenesis on the S5 sensor (**Figure S15**) to improve the FB1 response on the (7,5) and (10,2) chiralities. The final mutant, F-3, exhibited both an increased FB1 response as well as a decreased AFB1 response, resulting in a sensor with increased selectivity to FB1 compared to AFB1 for the (7,5) and (10,2) chiralities (**Figure 3d, Figure S15, Figure S16**). The (7,5) chirality showed the greatest FB1 selectivity owing to a lower response towards AFB1 (**Figure S16**). We therefore chose to monitor the FB1 response exclusively on the (7,5) chirality in the following experiments. In addition, we observed an improved selectivity of the Q-3 sensor towards AFB1 for (9,4) chirality compared to the S5 sequence, due to its reduced sensitivity towards FB1 (**Figure S17**).

Having engineered AFB1 and FB1 sensors, we further applied these sensors for the real-time multimodal detection of mycotoxins in food samples. In particular, the Q-3 and F-3 sensors were used to detect both AFB1 and FB1 at the (9,4) (ca. 1140 nm) and (7,5) (ca. 1050 nm) chirality wavelengths in corn flour media. Calibration curves were constructed to determine the sensitivity of the Q-3 sensor towards AFB1 and the F-3 sensor towards FB1 in presence of corn extract using the (9,4) and (7,5) chirality peaks (**Figures 4a-b**). The calibration curves were used to determine the limit of detection (LOD) of the Q-3 and F-3 sensors towards AFB1 and FB1, respectively (**Figure S20**). Following the addition of AFB1 to the Q-3 and FB1 to the F-3 sensor, respectively, we observed shifts of ≥ 0.24 (AFB1) and ≥ 0.35 nm (FB1) at a concentration of 1 μM (LOD) in corn extract. This LOD, coupled with the relatively easy sample preparation and speed of our sensors, makes the evolved DNA-SWCNT sensors attractive for applications monitoring the presence of these toxins in food and agricultural feed products^21,22^. To further investigate the applicability of our sensors, we demonstrated our sensors for AFB1 and FB1 detection in presence of other sources of toxin contamination such as almond flour (**Figure S21**). We noted that the presence of almond extract had almost no effect on the position of the (9,4) chirality, therefore allowing an efficient sensing of AFB1 in almond products. We evaluated the selectivity of our sensors by analyzing their response to toxin mixtures (**Figure S23**). The selectivity of the response was defined as the difference between the response towards a single toxin versus towards a mixture of toxins. By monitoring the (9,4) chirality of the Q-3 sensor, we were able to detect AFB1 from a mixture of both AFB1 and FB1 with very little interference from the FB1. Similarly, by monitoring the (7,5) chirality for the F-3 sensor, FB1 could be detected from a mixture of AFB1 and FB1, with no significant difference in response compared to the sample containing only FB1. The independence of sensor responsivity between individual and combined toxin solutions indicates the absence of a competitive interaction between the toxins and the nanotube sensor. We further tested the accuracy of our sensors by comparing the concentrations measured by SWCNT fluorescence to the concentrations as determined by HPLC-MS (**Figure S25, Table B.5**) for AFB1 and FB1 extracted from spiked corn flour. For the AFB1 response on the (9,4) chirality, the Q-3 sensor fluorescence showed greater agreement with the values determined by HPLC-MS compared to the F-3 sensor, the latter overestimating the AFB1 concentration due to the shifting contribution from FB1. On the other hand, for the FB1 response on the (7,5) chirality, the F-3 fluorescence gave better concentration estimations compared to the Q-3 sensor, both for single-toxin and mixed-toxin solutions.

**Figure 4:**
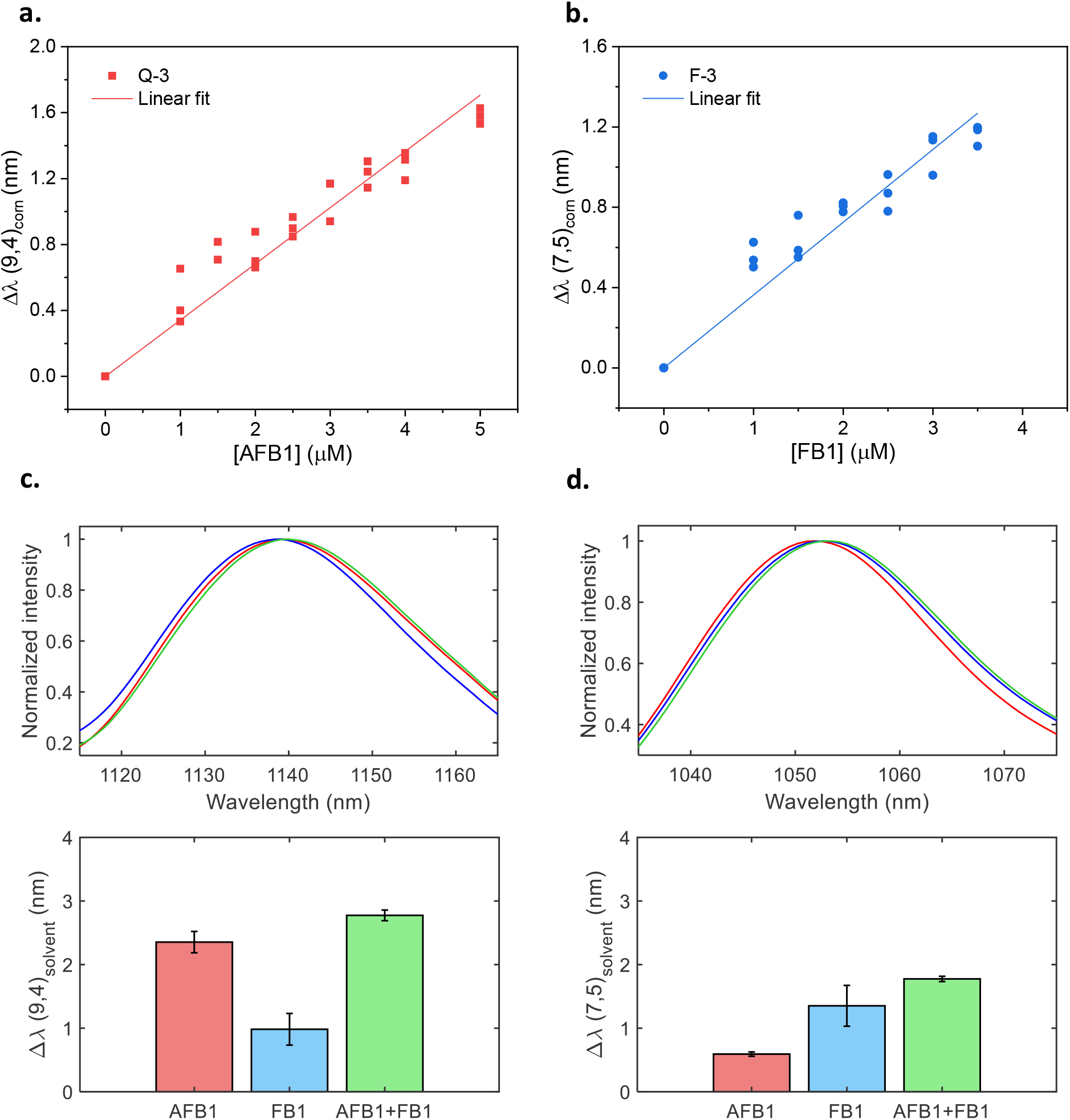
Selectivity of the evolved mycotoxin sensors. Calibration curves for the wavelength shifting response of (a) the (9,4) chirality of the Q-3 sensor and (b) the (7,5) chirality of the F-3 sensor. Measurements were taken for increasing concentrations of toxins suspended in a corn extract solution. The response was normalized to the response of the solvent (60% methanol) containing only corn extract. Measurements were taken following 320 min incubation. (c) The response of the (9,4) chirality of the mixed sensors after incubation with AFB1 (red, 10 μM), FB1 (blue, 10 μM) and the mixture of both toxins (green, 10 μM for each toxin). (d) The response of the the (7,5) chirality of the mixed sensors after incubation with AFB1 (red, 10 μM), FB1 (blue, 10 μM) and the mixture of both toxins (green, 10 μM for each toxin). Top: the normalized emission spectra. Bottom: the shifting response compared to addition of control (DMSO). Measurements were taken following 200 min incubation. Error bars represent 1 σ standard error (n = 3).

We took advantage of the SWCNT chirality-specific responses of the Q-3 and F-3 sensors to perform multimodal sensing of both toxins in a mixture by combining both sensors in a same solution (**Figures 4c-d, Figure S26**). Specifically, we demonstrated the ability to simultaneously detect AFB1 and FB1 by monitoring the (9,4) and (7,5) chiralities. Although previous works have demonstrated multimodal sensing using distinct fluorescence responses of different chiralities^23,24^, these studies required the separation of SWCNT chiralities, a tedious endeavor that is incompatible with scalable applications and limited to only certain nanotube chiralities. In this study, we obviate the need to separate SWCNTs by exploiting the preferential chirality-specificity of each DNA-sequence response. Since batches of SWCNTs were prepared separately with the Q-3 and F-3 sequences and then mixed together, the final suspension yielded (9,4) and (7,5) chiralities wrapped with both sequences. This mixture of on- and off-target DNA sequences on each chirality was shown to yield an overall decrease in selectivity for the mixed sensors compared to the selectivity of the individual sensors towards their specific toxins (**Figure S23**). Therefore, while directed evolution provides a promising avenue for achieving multimodal sensing in the absence of chirality separation, one must also consider possible compromises to sensitivities in mixtures when deciding on the desired minimum sensitivity threshold to achieve using directed evolution.

While our sensors did not require chirality sorting, they may strongly depend on the chirality distribution, which can vary depending on the method used to prepare the DNA-SWCNTs^25^. To examine this, we compared the AFB1 and FB1 responses of the Δ(AG), A-4, and S5 sensors prepared by both surfactant exchange and direct sonication (**Figure S27**). The sonicated samples and exchanged samples were both able to sense AFB1, however the response of the sonicated A-4 sensor was reduced compared to the exchanged sensor. On the other hand, no FB1 response was observed for sensors prepared by sonication. This lack of response for FB1 may be due to both the different wrapping structures that can occur on the SWCNT surface depending on the suspension protocol used^25,26^ and differences in the amount of free DNA present in the exchanged and sonicated samples. To explore the underlying mechanism of the toxin-induced responses of the evolved sensors, we examined the effect of possible changes in the local dielectric environment. Changes in the local dielectric environment can trigger wavelength shifting of nanotube emission^27^. While such a response could arise from the toxin’s direct effect on changing the dielectric constant of the solvent, water in the case of DNA-SWCNTs, previous studies have shown that AFB1 at concentrations used in our study cannot effectively alter the dielectric constant of the water^28^. The DMSO present in the toxin solution could also affect the dielectric constant of the sensor’s solution (final concentration 1%).

However, this effect is accounted for by measuring the shifting response versus a DMSO blank addition. Solvatochromic shifts can also result from changes in the exposed surface area of the nanotube^27,29^. Change in SWCNT surface coverage modulates the accessibility of water on the SWCNT surface, which can consequently alter the local dielectric environment of the SWCNT. Surface coverage-induced wavelength shifting has been observed extensively for various SWCNT sensors including surfactant-covered and DNA-wrapped SWCNTs^30–35^. Herein, we hypothesize that the interaction between the toxins and the DNA-SWCNTs could result in a change in SWCNT surface coverage, leading to the characteristic red-shifting observed in fluorescence and NIR absorbance (**Figure S28**).

In the case of AFB1, we tested this hypothesis by first examining the reversibility of Q-3 sensor. A washing protocol (see **Methods**) was used to remove the toxins from the DNA-SWCNT suspension while allowing a resuspension of the sensors. We applied the washing protocol to the Q-3 sensor after overnight incubation with AFB1 and monitored the position of the (9,4) peak (**Figure 5a, Figure S30**). After washing, we observed that the peak position was the same irrespective of whether the sensor had previously interacted with AFB1 or DMSO (negative control). In addition, we also noted that the washed Q-3 sensor showed a similar response to the unwashed Q-3 sensor upon subsequent additions of AFB1. A similar result was observed for the F-3 sensor (**Figure S30**). These findings suggest that AFB1 interacts with DNA-SWCNT non-covalently and that the interaction between AFB1 and the DNA-SWCNTs is reversible. Interestingly, we note a minimal red-shifting of the peak position as well as a slight decrease in shifting response for the washed samples. We attribute these observations to the removal of free DNA that occurs during the precipitation process (**Figure S34, Figure S35**).

**Figure 5:**
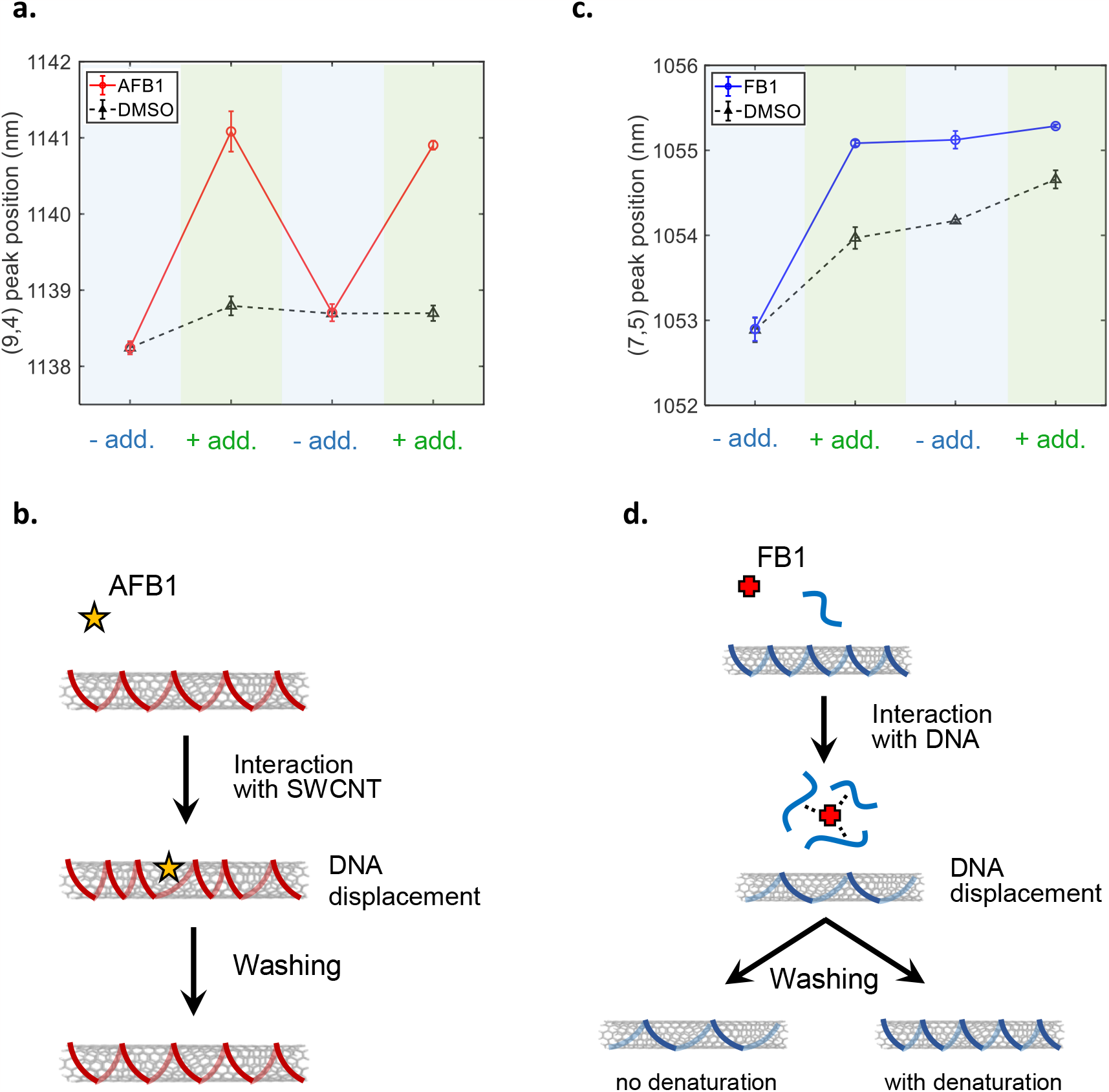
The sensors have two distinct mechanisms of interaction. (a, c) Peak position of the (a) Q-3 and (c) F-3 sensor in the absence (“-add”) and presence (“+add”) of AFB1 (red), FB1 (blue) and DMSO (black) for the (9,4) and (7,5) chiralities. Measurements were taken in the presence of 5 μM of the corresponding toxin following 200 min incubation. Error bars represent 1 σ standard error (n = 3). (b, d) Schematics of the proposed interaction between (b) AFB1 and DNA-SWCNT sensors, and (d) FB1 and DNA-SWCNT sensors. The AFB1 interacts directly with the SWCNT surface resulting in a displacement of the DNA wrapping. Upon washing, the AFB1 is removed from the SWCNT surface and the DNA wrapping recovers its initial conformation. FB1 interacts with the DNA (bound to SWCNT and unbound), leading to a conformational change of the wrapping. Upon washing, the sensor can be reverted back to its initial state only if the toxin-sensor interaction is disrupted by thermal denaturation.

We further examined whether this reversibility stems from a reversible interaction with the SWCNT, the DNA, or both elements. Previous studies have reported the interaction of AFB1 with DNA. In particular, the 8,9-epoxide exo isomer of AFB1 (AFBO) can form covalent adducts with guanine nucleobases, leading to the acute toxicity of the toxin^36^. Since the interaction AFB1 – DNA-SWCNT is reversible, we infer the lack of covalent interaction between the AFB1 and the DNA. In addition, we verified the absence of AFBO in our samples by HPLC-MS (**Figure S37**) and by additional measurements that confirmed a SWCNT fluorescence response towards aflatoxin B2 (AFB2) (**Figure S38**), which cannot form covalent adducts with DNA. In parallel, multiple studies have reported the non-covalent interaction of AFB1 to both double- and single-stranded DNA^37–39^ via AFB1-nucleobase binding. In the DNA-SWCNT hybrids, the DNA nucleobases are considered inaccessible^40^, hence preventing their interaction with AFB1 molecules.

The unlikely covalent and non-covalent means of interacting with DNA therefore suggests that the AFB1 primarily interacts with our sensor directly *via* the SWCNT sidewall. To test this hypothesis, we investigated the fluorescence response of sodium cholate (SC)-SWCNTs in the presence of AFB1 at various SC concentrations (**Figure S39**), equivalent to different surface coverages as previously described ^34^. As expected, no response was observed for the high coverage sample (46 mM SC); however, a strong red-shift of the (9,4) peak was detected upon addition of AFB1 for the sample with the lowest coverage (0.5 mM SC). Interestingly, for the other low-coverage samples (1 and 1.5 mM SC), the (9,4) peak first red-shifted before returning to the initial value over time. We attribute the observed shifting to the destabilization or reorganization of the SC corona upon AFB1 binding to SWCNT, therefore resulting in a change in water accessibility and a red-shifting of the emission. The red-shifting was also linked to a decrease in fluorescence intensity, indicative of a change in the SC corona on the SWCNT surface (**Figure S40**). While this change seemed to be permanent for the 0.5 mM SC sample, the interaction AFB1 – SWCNT might not be thermodynamically favorable compared to the SC – SWCNT interaction for the 1 and 1.5 mM SC samples, explaining the reversibility of the shifting. To verify the interaction between the AFB1 and the SWCNT sidewall, we added low concentrations of sodium dodecylbenzenesulfonate (SDBS) to DNA-SWCNT after interaction with AFB1 to replace the toxin on the SWCNT surface while leaving the DNA wrapping intact (**Figure S43**). At SDBS concentrations below 5 × 10^−3^ %, we observed that only the samples previously reacted with AFB1 would exhibit a blue-shift of the (9,4) peak, indicating a removal of AFB1 from the SWCNT surface. Based on these observations, we propose that the AFB1 interacts with the SWCNT surface, resulting in a displacement of the DNA on the SWCNT surface and the observed red-shift response (**Figure 5b**). These observations of sensor reversibility were further demonstrated in a working flow chamber. The sensors in this flow configuration showed reversible shifting upon washing (**Figure S46**) demonstrating a proof of principle device for the continuous monitoring of AFB1 contamination in corn- and almond-based products.

In the case of FB1, we observed that the (7,5) response of the F-3 sensor was not reversible even following the toxin washing (**Figure 5c, Figure S30**), indicating a relatively strong interaction between FB1 and the DNA-SWCNT. To verify this hypothesis, we designed a modified version of the washing protocol where the samples were heated to 95°C before sample precipitation to denature any strong non-covalent interactions between the toxin and the F-3 sensor (**Figure S33**). With this additional denaturation step, we noted that the FB1 response was reversible, as evidenced by the equivalent (7,5) peak position for both the FB1 and DMSO samples after washing. This heat-activated reversibility therefore implies that FB1 interacts in a strong, non-covalent manner with the DNA-SWCNT. This hypothesis is further supported by the hydrophilicity of the FB1, which would diminish its interaction with the hydrophobic SWCNT surface, as supported by the lack of response towards FB1 for low coverage SC-SWCNT samples (**Figure S41, Figure S42**). The interaction between DNA and FB1 has not been thoroughly studied, but the discovery of aptamers towards FB1^41^ confirms the possibility of a non-covalent interaction such as hydrogen bonding with both the bases and backbone. The role of free DNA in the F-3 sensor response supports the hypothesis that FB1 interacts with the DNA in the hybrid. Upon removal of free DNA, the sensors exhibited a red-shifting of the emission and a much lower FB1 response (**Figure S35, Figure S36**). This decrease in response is similar to the lack of response observed after washing for both the non-denatured and denatured samples (**Figure S30, Figure S33**). This similarity was attributed to the loss of free DNA during the washing step (**Figure S34**), indicating that the free DNA is crucial for the FB1 interaction. Furthermore, we also observed an increase in the concentration of free DNA for the sensors that reacted with FB1 compared to those reacted with AFB1 or DMSO (**Figure S44**). This observation implies that the interaction with FB1 could result in the facilitated removal or weakening of the DNA interaction with the SWCNT surface. Together, these observations suggest that the FB1 interacts non-covalently with wrapped and free DNA in solution, contributing to a displacement of the DNA on the SWCNT surface and a red-shifting of the emission (**Figure 5d**). Upon washing, denaturation can be used to dissociate the DNA from the toxin and restore the initial wrapping (**Figure 5d**).

This hypothesis was further corroborated by the response behavior of the Q-3 sensor towards FB1 post-washing. While the F-3 sensor was unresponsive after toxin washing, the Q-3 sensor exhibited a slight response towards FB1 (**Figure S30, Figure S33**). This observation is in accordance with the fact that the free DNA removal had a lower effect on the FB1 response of the Q-3 compared to the F-3 sensor (**Figure S36**). We hypothesize this difference is linked to the strength of the interaction between the DNA wrapping and the SWCNT surface. To determine the binding affinity of the Q-3 and F-3 sequences to SWCNT, we performed surfactant replacement experiments using SDBS (**Figure S45**). We observed that the surfactant replacement occurred faster for the F-3 sensor compared to the Q-3 sensor, indicating that the F-3 sequence has a lower binding affinity to the SWCNT surface. In addition, measurements of the free DNA concentrations (**Figure S44**) showed that the F-3 sensor had more unbound DNA than the Q-3 sensor, further suggesting a weaker binding of the DNA to the SWCNT surface. These results suggest that the F-3 wrapping is more prone to conformational changes upon free DNA removal, as well as upon toxin interaction.

## Conclusion

In this study, we demonstrated the applicability of directed evolution for the creation of DNA-SWCNT sensors for mycotoxin detection. We showed that it is possible to significantly improve the response of DNA-SWCNT sensors towards two distinct mycotoxins, without requiring any prior knowledge on the interaction mechanisms of these toxins with DNA-SWCNTs. In addition, we showcased how DNA shuffling can be used to drastically accelerate the evolution of these sensors. We further demonstrated that directed evolution can be used to evolve not only the magnitude of response, but also the brightness and selectivity of our sensors. Moreover, we showed that it is possible to independently evolve these sensors for two specific chiralities, (9,4) and (7,5), advantageous for multimodal sensing applications. The applicability of these sensors for toxin detection was also shown in food samples, specifically a dissolved corn flour matrix, without any need for complicated sample preparation. We discovered two distinct sensing interactions that were dependent on the mycotoxin type, highlighting the power of directed evolution in the discovery of new nanotube sensors.

We believe this study can have an important impact on the engineering of DNA-SWCNT sensors. As shown herein, the directed evolution methodology can be used to tune several aspects of DNA-SWCNT sensors in order to optimize their performance for a range of applications. While applied here for the sensing of mycotoxins, the same methodology could be applied to variety of targets including small molecules^6,33^, biomarkers^32,42^ and viruses^43^. Moreover, we believe that the findings of this study will have a more widespread impact on the identification of more rational design rules for DNA-SWCNT sensors. Beyond SWCNT sensors, directed evolution presents a very powerful tool for engineering other kinds of nanocomplexes exhibiting unknown structure-function relationships. As such, this study helps catalyze the fusion of synthetic biology approaches with nanomaterials, with the aim of unlocking the true potential of these hybrid materials.

## Acknowledgements

The authors are thankful for funding support from the Swiss National Science Foundation (SNSF), the European Research Council (ERC) under the European Union’s Horizon 2020 research and innovation program (grant agreement no. 853005) and the Honda Research Institute.

## References

1. Mousavi, S. M., Hashemi, S. A., Zarei, M., Amani, A. M. & Babapoor, A. Nanosensors for Chemical and Biological and Medical Applications. Med. Chem. (Los. Angeles). 08, (2018).

2. Riu, J., Maroto, A. & Rius, F. X. Nanosensors in environmental analysis. Talanta 69, 288–301 (2006).

3. Srivastava, A. K., Dev, A. & Karmakar, S. Nanosensors and nanobiosensors in food and agriculture. Environ. Chem. Lett. 16, 161–182 (2018).

4. Weissleder, R., Kelly, K., Sun, E. Y., Shtatland, T. & Josephson, L. Cell-specific targeting of nanoparticles by multivalent attachment of small molecules. Nat. Biotechnol. 23, 1418–1423 (2005).

5. Zhang, J. et al. Single molecule detection of nitric oxide enabled by d(AT)15 DNA adsorbed to near infrared fluorescent single-walled carbon nanotubes. J. Am. Chem. Soc. 133, 567–581 (2011).

6. Kruss, S. et al. Neurotransmitter detection using corona phase molecular recognition on fluorescent single-walled carbon nanotube sensors. J. Am. Chem. Soc. 136, 713–724 (2014).

7. Beyene, A. G. et al. Ultralarge Modulation of Single Wall Carbon Nanotube Fluorescence Mediated by Neuromodulators Adsorbed on Arrays of Oligonucleotide Rings. Nano Lett. 18, 6995–7003 (2018).

8. Lambert, B. P., Gillen, A. J. & Boghossian, A. A. Synthetic Biology: A Solution for Tackling Nanomaterial Challenges. J. Phys. Chem. Lett. 11, 4791–4802 (2020).

9. Romero, P. A. & Arnold, F. H. Exploring protein fitness landscapes by directed evolution. Nat. Rev. Mol. Cell Biol. 10, 866–876 (2009).

10. Lambert, B., Gillen, A. J., Schuergers, N., Wu, S.-J. & Boghossian, A. A. Directed evolution of the optoelectronic properties of synthetic nanomaterials. ChemComm 55, 3239–3242 (2019).

11. Gillen, A. J. & Boghossian, A. A. Non-covalent Methods of Engineering Optical Sensors Based on Single-Walled Carbon Nanotubes. Front. Chem. 7, 1–13 (2019).

12. Soares, R. R. G. et al. Advances, challenges and opportunities for point-of-need screening of mycotoxins in foods and feeds. Analyst 143, 1015–1035 (2018).

13. Singh, J. & Mehta, A. Rapid and sensitive detection of mycotoxins by advanced and emerging analytical methods: A review. Food Sci. Nutr. 8, 2183–2204 (2020).

14. Eskola, M. et al. Worldwide contamination of food-crops with mycotoxins: Validity of the widely cited ‘FAO estimate’ of 25%. Crit. Rev. Food Sci. Nutr. 60, 2773–2789 (2020).

15. Tola, M. & Kebede, B. Occurrence, importance and control of mycotoxins: A review. Cogent Food Agric. 2, (2016).

16. Rasch, C., Kumke, M. & Löhmannsröben, H. G. Sensing of Mycotoxin Producing Fungi in the Processing of Grains. Food Bioprocess Technol. 3, 908–916 (2010).

17. Jeong, S. et al. High-throughput evolution of near-infrared serotonin nanosensors. Sci. Adv. 5, 1–13 (2019).

18. Mann, F. A., Herrmann, N., Meyer, D. & Kruss, S. Tuning selectivity of fluorescent carbon nanotube-based neurotransmitter sensors. Sensors (Switzerland) 17, (2017).

19. Crameri, A., Raillard, S. A., Bermudez, E. & Stemmer, W. P. C. DNA shuffling of a family of genes from diverse species accelerates directed evolution. Nature 391, 288–291 (1998).

20. Socha, R. D. & Tokuriki, N. Modulating protein stability - Directed evolution strategies for improved protein function. FEBS J. 280, 5582–5595 (2013).

21. Anukul, N., Vangnai, K. & Mahakarnchandkul, W. Significance of regulation limits in mycotoxin contamination in Asia and risk management programs at the national level. J. Food Drug Anal. 21, 227–241 (2013).

22. Rodrigues, I. & Naehrer, K. A three-year survey on the worldwide occurrence of mycotoxins in feedstuffs and feed. Toxins (Basel). 4, 663–675 (2012).

23. Giraldo, J. P. et al. A Ratiometric Sensor Using Single Chirality Near-Infrared Fluorescent Carbon Nanotubes: Application to in Vivo Monitoring. Small 11, 3973–3984 (2015).

24. Nißler, R. et al. Sensing with Chirality-Pure Near-Infrared Fluorescent Carbon Nanotubes. Anal. Chem. 93, 6446–6455 (2021).

25. Gillen, A. J., Lambert, B. P., Antonucci, A., Molina-Romero, D. & Boghossian, A. A. Modulating the properties of DNA-SWCNT sensors using chemically modified DNA. bioRxiv (2021).

26. Yang, Y., Sharma, A., Noetinger, G., Zheng, M. & Jagota, A. Pathway-Dependent Structures of DNA-Wrapped Carbon Nanotubes: Direct Sonication vs. Surfactant/DNA Exchange. J. Phys. Chem. C (2020) doi:10.1021/acs.jpcc.0c00679.

27. Choi, J. H. & Strano, M. S. Solvatochromism in single-walled carbon nanotubes. Appl. Phys. Lett. 90, 88–91 (2007).

28. Muga, F. C., Workneh, T. S. & Marenya, M. O. Electrical properties of maize kernels contaminated with aflatoxin. Agric. Eng. Int. CIGR J. 20, 197–205 (2018).

29. Gillen, A. J. et al. The impact of surface area on the response of SWCNT optical sensors. 41, 1–20.

30. Salem, D. P. et al. Ionic strength mediated phase transitions of surface adsorbed DNA on single-walled carbon nanotubes. J. Am. Chem. Soc. 139, 16791–16802 (2017).

31. Gillen, A. J., Kupis-Rozmysłowicz, J., Gigli, C., Schuergers, N. & Boghossian, A. A. Xeno Nucleic Acid Nanosensors for Enhanced Stability Against Ion-Induced Perturbations. J. Phys. Chem. Lett. 9, 4336–4343 (2018).

32. Harvey, J. D. et al. A carbon nanotube reporter of microRNA hybridization events in vivo. Nat. Biomed. Eng. 1, 1–11 (2017).

33. Harvey, J. D., Baker, H. A., Mercer, E., Budhathoki-Uprety, J. & Heller, D. A. Control of Carbon Nanotube Solvatochromic Response to Chemotherapeutic Agents. ACS Appl. Mater. Interfaces 9, 37947–37953 (2017).

34. Gillen, A. J. et al. Templating colloidal sieves for tuning nanotube surface interactions and optical sensor responses. J. Colloid Interface Sci. 565, 55–62 (2020).

35. Gillen, A. J., Antonucci, A., Reggente, M., Morales, D. & Boghossian, A. A. Distinguishing dopamine and calcium responses using XNA-nanotube sensors for improved neurochemical sensing. bioRxiv (2021).

36. Johnson, W. W. & Guengerich, F. P. Reaction of aflatoxin B1 exo-8,9-epoxide with DNA: Kinetic analysis of covalent binding and DNA-induced hydrolysis. Proc. Natl. Acad. Sci. U. S. A. 94, 6121–6125 (1997).

37. Misra, R. P., Muench, K. F. & Humayun, M. Z. Covalent and Noncovalent Interactions of Aflatoxin with Defined Deoxyribonucleic Acid Sequences. Biochemistry 22, 3351–3359 (1983).

38. Stone, M. P., Gopalakrishnan, S., Harris, T. M. & Graves, D. E. Carcinogennucleic acid interactions:Equilibrium binding studies of aflatoxins b1 and b2. J. Biomol. Struct. Dyn. 5, 1025–1041 (1988).

39. Ma, L., Wang, J. & Zhang, Y. Probing the characterization of the interaction of aflatoxins B1 and G1 with calf thymus DNA in vitro. Toxins (Basel). 9, (2017).

40. Johnson, R. R., Johnson, A. T. C. C. & Klein, M. L. Probing the structure of DNA-carbon nanotube hybrids with molecular dynamics. Nano Lett. 8, 69–75 (2008).

41. McKeague, M. et al. Screening and initial binding assessment of fumonisin B 1 aptamers. Int. J. Mol. Sci. 11, 4864–4881 (2010).

42. Yaari, Z. et al. Machine-Perception Nanosensor Platform to Detect Biomarkers of Ovarian Cancer. bioRxiv (2021).

43. Pinals, R. L. et al. Rapid SARS-CoV-2 Spike Protein Detection by Carbon Nanotube-Based Near-Infrared Nanosensors. Nano Lett. 21, 2272–2280 (2021).

