## Supplementary Information for "Directed evolution of nanosensors for the detection of mycotoxins"

### Supplementary information:
### Directed evolution of nanosensors for the detection of mycotoxins

**Authors:** Benjamin P. Lambert[1], Afsaneh Taheri[2,3], Shang-Jung Wu[1], Alice J. Gillen[1], Mahdi Kashaninejad[2], Ardemis A. Boghossian[1]*

[1]Department of Chemistry and Chemical Engineering, École Polytechnique Fédérale de Lausanne (EPFL); Lausanne, Switzerland

[2]Department of Food Process Engineering, Faculty of Food Science and Technology, Gorgan University of Agricultural Sciences and Natural Resources, Gorgan, Iran

[3]Food, Chemical and Biotechnology Cluster, Singapore Institute of Technology, Singapore

#### Materials and methods

##### *Materials*

All mycotoxins were purchased from Enzo Life Sciences. All DNA oligomers were purchased from Microynth AG. Super-purified HiPco SWCNTs were purchased from NanoIntegris (Batch No. HP32-130). All chemicals were purchased from Sigma-Aldrich, unless otherwise indicated. The corn powder, almond powder and beer used in the experiments were purchased from a local grocery store.

##### *Instruments*

Absorbance measurements were collected using a UV-3600 Plus spectrophotometer (Shimadzu) and quartz suprasil cuvettes (Z600393, Hellma Analytics). Absorbance measurements were also performed in 96- and 384-well plates using a plate reader (Varioskan LUX, Thermo Fisher). The near-infrared fluorescence measurements were acquired on a custom-made microscope as previously described[1]. Briefly, the system is built on a Nikon Ti-E inverted microscope body and illuminated by a supercontinuum laser (SuperK Extreme EXR-15 and SuperK VARIA, NKT Photonics) in the 400-830 nm range. The emitted light is diffracted between 900-1400 nm by a spectrometer (IsoPlane SCT-320, Princeton Instruments) using a 70 lines.mm$^{-1}$ grating and recorded by an InGaAs detector (NIRvana 640 ST, Princeton Instruments). The measurements were recorded automatically using a custom LabVIEW (National Instruments) program. The fluorescence data was analyzed by a custom code in Matlab (R2017b, MathWorks), in which individual fluorescence peaks were fitted with a single Lorentzian curve.

##### *Preparation of SWCNT suspensions*

The sodium cholate (SC)-SWCNT suspension was prepared by mixing 45 mg HiPco SWCNT powder with 45 mL 2% (w/w) SC prepared in deionized (DI) water. This mixture was homogeneized at 5,000 rpm (Polytron PT 1300 D, Kinematica) at room temperature for 20 min, prior to 60 min ultrasonication (amplitude 10, 1/4 in. tip, Q700 Sonicator, Qsonica) in an ice bath (the ice was changed after 30 min). The resulting suspension was centrifuged for 10 min at 15,000 x g at 25°C (Allegra 25R, Beckman Coulter) and the supernatant was further centrifuged for 4 h at 30,000 rpm at 25°C (SW32 Ti, Optima XPN-80, Beckman Coulter). The supernatant (top 90%)

was further concentrated using centrifugation filtration devices (Amicon Ultra-15 3kDa, Merck). Briefly, 10 mL was added to an Amicon device and centrifuged at 4,000 x g for 30 min (Allegra 25R, Beckman Coulter) at 25°C. An additional 10 mL was added to the device and the previous step was repeated. The concentrated nanotube suspension was resuspended using 2% (w/w) SC to a total volume of 6.5 mL and centrifuged at 3,300 x g for 30 min at 25°C (5810R, Eppendorf) to remove aggregates. The supernatant was collected and diluted using 2% (w/w) SC to a final concentration of 108 mg/L (measured using the extinction coefficient $\varepsilon$=0.036 $L.cm^{-1}.mg^{-1}$ at 632 nm[2]). The concentration step was important as most batches could reach only a concentration of approximately 70 mg/L, not enough for our experiments. Moreover, as the batches prepared on different days would not result on the same SWCNT yield, it is important to fix the final concentration in order to improve reproducibility. Nevertheless, we still observed a slight variation of SWCNT chirality distribution between various batches of SC-SWCNT suspensions (**Supplement 1**) which could potentially impact the reproducibility of the DNA-SWCNT suspensions.

The DNA-SWCNT sensors were prepared using a wrapping exchange protocol between the sodium cholate (SC)-SWCNTs and the DNA oligomer as previously described[2]. The DNA solutions were dissolved in DI water to a concentration of 50 µM (measured using Nanodrop 2000, Thermo Scientific). Ratios of 0.2:0.2:0.6 of DNA:SC-SWCNT:methanol were mixed gently (DNA and SC-SWCNT mixed first) and incubated for 2 h at room temperature. The DNA-SWCNTs were then purified to remove the free SC and methanol. First, 1.5 M NaCl was added to the mixture to a final concentration of 0.2 M, followed by the addition of 2.5 volumes of cold ethanol (kept at -20°C) and the solution was gently mixed. The resulting suspension was incubated at -20°C for 1 h and then centrifuged at 21,130 x g (5424R, Eppendorf) for 30 min at 4°C. The supernatant was removed, and the SWCNT-containing pellet was washed with 70% (v/v) ethanol (the pellet should break into small clumps), vortexed briefly, and centrifuged at 21,130 x g for 1 h at 20°C. The supernatant was discarded, and the pellet was air-dried for 12 min at room temperature. The pellet was then resuspended in the desired volume of 0.1 M NaCl, briefly vortexed, and finally centrifuged at 21,130 x g for 30 min at 20°C. The supernatant of this suspension was collected and its concentration was determined by measuring the absorbance of 90 µL of suspension in 96-well plate (Varioskan LUX plate reader, Thermo Scientific). The suspensions were all diluted with 0.1 M NaCl to a final concentration of 7.4 mg/L (corresponding to a background-subtracted absorbance of 0.066). All samples were freshly prepared and used on the same day for experiments in order to avoid variations between samples due to possible aging of the samples. Indeed, we noted a slight aging of our samples which seemed to stabilize after one day at room temperature (**Supplement 1**). DNA-SWCNT prepared by sonication were prepared by mixing 1 mg HiPco SWCNT with 1 mL of DNA solution at 100 μM concentration (≈ 1 mg/mL) followed by 90 min ultrasonication (amplitude 1, cuphorn, Q700 Sonicator, Qsonica) in an ice bath. The resulting suspension was centrifuged for 4 h at 21,130 x g at 4°C (5424R, Eppendorf) and the supernatant was stored at 4°C before use.

*Preparation of the mycotoxin solutions*

Unless otherwise indicated, mycotoxin powders were dissolved in dimethyl sulfoxide (DMSO) in a nitrogen-controlled glovebox (E-line, GS Glovebox). The concentrations of all mycotoxins, except for FB1, were verified using their respective extinction coefficients in the 200-400 nm range (**Supplement 2**). All dissolved mycotoxin samples were aliquoted and stored at -20°C in the dark. The solubility and purity of the toxins (AFB1 mainly) were also assayed (**Supplement 2**).

*DNA mutant library*

Mutants of the DNA sequences were generated by a custom code in Matlab (**Supplement 3**). For the creation of the single mutation library, a random base position (between 1 and 30) was first chosen and then randomly replaced by any of the three other bases with the same probability. For the creation of more than one mutation, the process explained above was repeated on the rest of the non-mutated bases. The initial screening library used to find sensors responsive to AFB1 was composed of repeating sequence variants, constructed by introducing three random mutations in $(AT)_{15}$, $(AG)_{15}$, $(CT)_{15}$, and $(AC)_{15}$ sequences. The choice of repeating sequences with alternating bases was made considering the extensive body of research on SWCNTs functionalized with such sequences, in particular for the creation of sensors[3–6]. As previously observed[2], the introduction of three mutations enables high enough diversity in the starting search library. In this study, we have screened 438 sequences in total: an initial library composed of 100 $(AT)_{15}$, $(AG)_{15}$, $(CT)_{15}$ and $(AC)_{15}$ variant sequences and 338 mutants derived from the Δ(AG) sequence. We have summarized the main sequences used in the study in the **Table S2**.

*DNA shuffling*

DNA shuffling can be used to generate improved mutants with a high degree of fitness by combining the beneficial mutations of one or multiple sequences. To generate the shuffled sequences (**Supplement 3**), the selected sequences are first cut into *n* fragments (*n* equals the length of the sequence divided by the fragment size). The small fragments are then randomly shuffled and the number of fragments necessary to achieve the specified sequence length are selected and combined to create the so-called shuffled sequence. The library of all recombined sequences is given by the permutations of all fragments into sequences composed of *n* fragments. As we performed the sequence shuffling computationally, we had the advantage of being able to choose a fixed fragment length. In this study, we tested fragment sizes of 5, 6 and 10 nucleotides (**Figure S11**). We estimated this range to be optimal for the discovery of mutants as smaller fragments might be too short to contain a recognition site and longer fragments would be too large to enable sufficient sequence diversity. We also ruled out the sequences that contained more than half of unmutated fragments to maximize the presence of beneficial mutations in the sequence.

*Food matrix extraction and spiking*

Corn extract was used alone or in combination with the toxins in order to assay response selectivities. The extraction was performed using a modified version of a previously reported protocol[7]. 5 g of corn powder was mixed with 25 mL 100% methanol and shaked at 3,000 $min^{-1}$ (uniTEXER, LLG) for 1 h. Then, the solution was centrifuged at 3,000 x g for 10 min at 4°C (5810R, Eppendorf) and the supernatant was filtered (Whatman 597). The filtrate was further centrifuged at 21,130 x g for 10 min and stored at room temperature (5424R, Eppendorf). The same protocol was used for the almond flour extract, except it was filtered using a 0.22 µm PTFE filter because of the smaller volume. The beer sample was used directly out of the bottle.

For the spiking experiments, 100 mg of corn powder was mixed in a 1.5 mL Eppendorf tube with 20 µL of toxin solution at a final concentration of 2.5 mM. The solution consisted of 5 µL toxin at 10 mM, 5 µL DMSO (or toxin if there were two toxins at the same time) and 10 µL 60% methanol. Control samples without corn were made by mixing 20 µL of the toxin solution described above with 180 µL of 60% methanol. The powder was vortexed for 15 s and left to incubate with the toxin for 30 min at room temperature. Subsequently, 180 µL of 60% methanol was added and the tube was shaken at 1,500 rpm for 30 min at 20°C (Thermomixer C, Eppendorf). The tube was then centrifuged at 21,130 x g (5424R, Eppendorf) for 1 min. The supernatant was filtered through a 0.22 µm PTFE filter to remove large corn particles. We verified that the AFB1 could go through the PTFE filters by absorbance measurements (**Figure S24**).

*Preparation of the immobilized DNA-SWCNTs*

Films of DNA-SWCNTs were created in order to test the reversibility of the sensors. First 20 µL of DNA-SWCNTs was added to a 96-well plate with glass bottom and the sensors were dried at 70°C for 1 h. This method allows to obtain films with a SWCNT concentration sufficient to perform spectroscopy. Then, 40 µL of warm 2% (w/v) agarose (in 0.1 M NaCl) was added on top of the dried SWCNTs in order to avoid sample removal from the focal plane upon analyte addition. 100 µL 0.1 M NaCl was added and the fluorescence was monitored for 20 min (time interval of 150 s with laser off between measurements). The NaCl solution was removed and replaced with 100 µL of 20 µM AFB1 solution in 0.1 M NaCl and the fluorescence was monitored for 1 h (time interval of 150 s with laser off between measurements).

*Mycotoxin removal protocol*

The washing process to remove mycotoxins from DNA-SWCNT suspensions is illustrated in **Figure S30**. The DNA-SWCNTs were first mixed with the toxins and the response was measured. Following overnight incubation with the toxin, the DNA-SWCNTs were precipitated by salt-assisted ethanol precipitation following the same protocol described previously for the purification of the DNA-SWCNT suspensions. The samples were incubated with ethanol at -20°C for 1 h and centrifuged at 21,130 g for 30 min at 4°C. The DNA-SWCNT pellet was washed with 70% ethanol and centrifuged at 21,130 g for 1h at 20°C. The supernatant was discarded and the pellet was air-dried for 12 min. Finally, the DNA-SWCNT pellet was resuspended in 0.1 M NaCl and centrifuged at 21,130 g for 30 min at 20°C to remove aggregates and the response of the sensor towards mycotoxins was measured again. For washing under denaturing conditions (**Figure S33**), the DNA-SWCNTs were heated to 95°C for 5 min (Thermomixer C, Eppendorf) immediately before the salt-assisted ethanol precipitation step. We verified that our DNA-SWCNT washing procedure was effectively removing the toxin both in the absence (**Table S6**) and presence of DNA-SWCNTs (**Figure S31**), using HPLC-MS and UV absorbance, respectively.

*Surfactant replacement assays*

The DNA wrapping was replaced using both sodium deoxycholate (SDC) and sodium dodecylbenzenesulfonate (SDBS) surfactants following previously reported procedures[8,9]. The SDC was dissolved to a concentration of 1% (w/w) in DI water. 45 µL of DNA-SWCNT suspension was mixed with 5 µL of 1% SDC (final concentration of 0.1 %) in a 384-well plate (MaxiSorp, Nunc) and incubated at room temperature. The fluorescence of the suspension was recorded before, 15 min and after SDC addition and after an overnight incubation. The SDBS was dissolved in DI water and diluted to concentrations ranging from 0.0138 to 0.55% (w/w). 20 µL (or 50 µL) of DNA-SWCNT suspension was mixed with 2 µL (or 5 µL) of SDBS solution, with final SDBS concentrations ranging from $1.25 \times 10^{-3}$ to 0.05%, and were incubated at room temperature before the fluorescence was recorded.

*Free DNA removal*

The free DNA was removed using Amicon Ultra-0.5 100 kDa devices. The membrane of the device was first rinsed with 500 µL DI water at 14,000 x g for 5 min. Then, DNA-SWCNT suspension and DI water were added to the device for a total volume of 500 µL and the device was centrifuged at 4,000 x g for 2 min. If analysis of the free DNA concentration is required, the flow through can be stored for further processing. DI water was added to the device to reach a final volume of 500 µL and the DNA-SWCNTs stuck on the membrane were resuspended by pipetting up and down. The samples were centrifuged, washed and resuspended again 5 additional times in order to remove all traces of free DNA. The last centrifugation was done for 5 min instead of 2 min. Finally, the samples were resuspended in 0.1 M NaCl and centrifuged to remove SWCNT aggregates. The final suspensions were diluted to the same concentration as the initial suspensions before filtering ($A_{632}$(90 µL) ≈ 0.066 after blank correction).

*DNA quantification*

The DNA concentrations of the flow-throughs of centrifugation devices and supernatants of the DNA-SWCNTs were assayed using the fluorescence of the Sybr Gold nucleic acid dye. For the ethanol supernatants, ethanol was removed by filtrating the supernatant (500 µL) with Amicon Ultra-0.5 10kDa centrifugation devices (pre-rinsed) two times with DI water followed by resuspension of the DNA in the filter in DI water (50 µL). For the measurement, 10 µL of DNA sample was added to 40 µL 1X Sybr Gold in 10 mM HEPES buffer (pH 8). The HEPES buffer was used to keep the pH within the optimized range for the dye (between pH 7.0 and 8.5). The samples were then incubated in the dark for 10 min and the fluorescence at 537 nm under 495 nm excitation was recorded with an exposure of 100 ms (Varioskan LUX, Thermo Fisher). We constructed calibration curves of Sybr Gold fluorescence and DNA concentration for each sequence (**Figure S34**) and used the slopes to find the concentration of DNA in our samples.

*Measurements*

Fluorescence measurements were acquired with an exposure time of 5 s, a laser power of 100% and an illumination bandwidth of 10 nm. For the monitoring of the (7,5) and (7,6) chiralities, an excitation of 660 nm was used while the monitoring of the (10,2) and (9,4) chiralities was done with an excitation of 745 nm. 49.5 µL aliquots of diluted DNA-SWCNT suspension were added to a 384-well plate (Nunc Maxisorp, Thermo Scientific) and fluorescence spectra were collected prior to the addition of any analyte. Following the initial measurement, 0.5 µL of analyte solution (concentration specified in the experiment) was added to the well and briefly mixed by pipetting up and down. Spectra were acquired at several time intervals, starting 80 min post-addition and repeated every 120 min for a total of a total of at least 200 min. All measurements were taken at room temperature and the laser was switched off in between measurements. The measurements were also done with sealing tape (CLS6524, Corning) on the plates to avoid evaporation. As the kinetics showed that the steady state of the response for both toxins occurred after approximately 200 min (**Figure S49**), most measurements were compared 200 min post-addition. For the experiments in presence of corn extract, the measurements were compared 320 min post-addition, as this was the time point where the background response due to the corn extract stabilized (**Figure S18**). The shifting responses labelled as "$\Delta\lambda_{solvent}$", "Peak shift" or "Wavelength shift" were calculated as the difference in peak position between the toxin and the negative control (DMSO or methanol depending on the sample). The shifting responses labelled as "$\Delta\lambda_{corn}$" correspond to the difference in peak position between the toxin and the negative control both containing corn extract. The shifting responses labelled as "$\Delta\lambda_{t0}$" was calculated compared to the peak position before addition (t0).

#### Supplement 1: DNA-SWCNT suspensions

While we obtained a good reproducibility in the preparation of HiPco SWCNTs with sodium cholate (SC), we observed variations in the absorbance spectra of different batches of HiPco-2% SC (**Figure S1**). We attribute these variations to variations in sonication conditions due to the lack of automation in the process. We hypothesize these differences could lead to variations in the behavior of DNA-SWCNT suspensions prepared with different HiPco-SC batches.

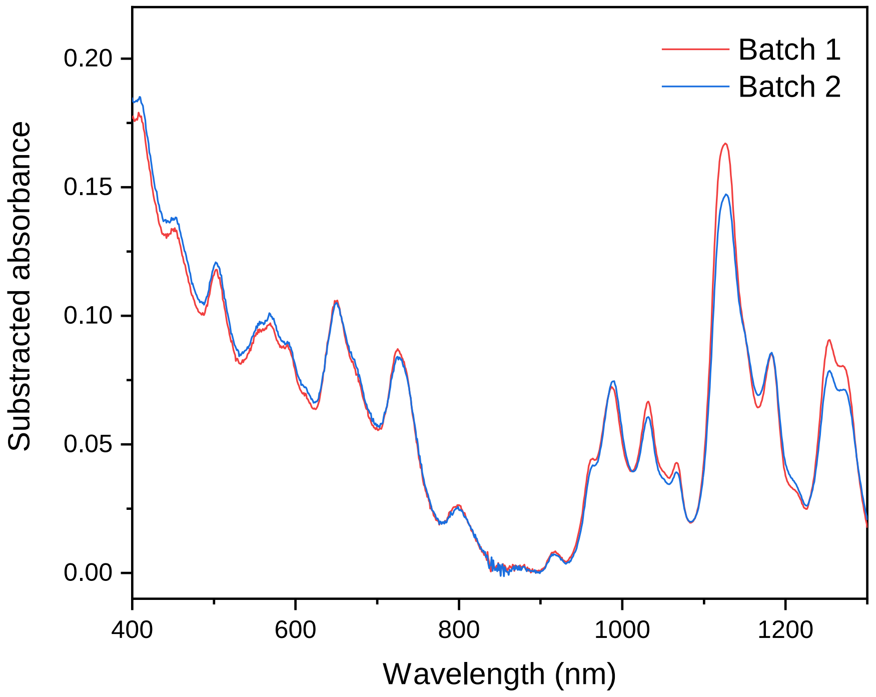


**Figure S1** - Differences between HiPco-SC batches. Absorbance spectra of HiPco-SC (2 % SC) batches. The spectra are both subtracted by the background value at 894 nm.

We observed an aging of the DNA-SWCNT complexes. This aging was characterized mostly by a significant decrease in fluorescence intensity, but did not seem to strongly affect the response of the sensors (**Figure S2**). We attribute this aging to the reorganization of the DNA wrapping from a meta-stable stable (post-resuspension) to an equilibrium (after one day), consistent to the conclusions of Yang *et al.*[10].

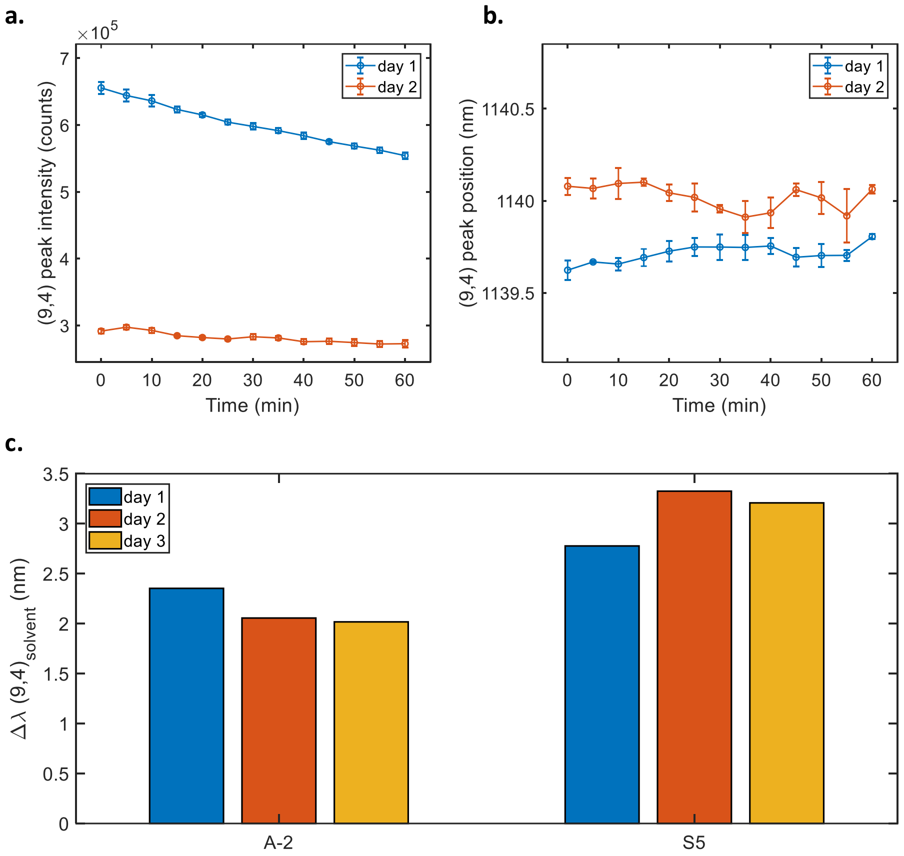


**Figure S2** - Aging of the DNA-SWCNT complexes. Effect of aging on (a) the fluorescence intensity, (b) peak position and (c) AFB1 shifting response of the sensors for the (9,4) chirality. The kinetics (a,b) were performed on the F-3 sensor. The dependence on AFB1 shifting response was performed on the A-2 and S5 sensors after a post-addition time of 200 min and with an AFB1 concentration of 10 μM. Although aging affects significantly the intensity of the sensors, it does not affect strongly the position of the emission peak or the analyte response.

#### Supplement 2: Mycotoxin solutions

The concentrations of all mycotoxins, except for FB1, were verified using their respective extinction coefficients in the 200-400 nm range listed in the **Table S1**. It is important to note that different batches of mycotoxins were used over the course of the study (mainly for AFB1 and FB1). Although the concentrations were fixed by absorbance for AFB1, slight variations may arise between batches. Such variations may then result in slight variations in DNA-SWCNT response observed between different batches of experiments.

**Table S1 -** List of the extinction coefficients ε at the respective wavelengths for the mycotoxins used in the study. The toxins were diluted in the indicated solvent before UV-VIS measurement.

| Mycotoxin | ε ($L.cm^{-1}.mM^{-1}$) | Wavelength (nm) | Solvent | Ref. |
|---|---|---|---|---|
| AFB1 | 21.8 | 363 | Ethanol | 11 |
| OTA | 5.5 | 333 | Methanol | 12 |
| ZEN | 6.02 | 316 | Methanol | 13 |
| DON | 5.28 (6.81) | 235 (221) | Acetonitrile | Estimated from 14,15 |

The toxin batches prepared before December 2019 were not prepared in an inert atmosphere and could therefore present traces of toxin oxidation. The most striking evidence of such toxin degradation occurred for one AFB1 batch which changed color from transparent to light pink (**Figure S3**).

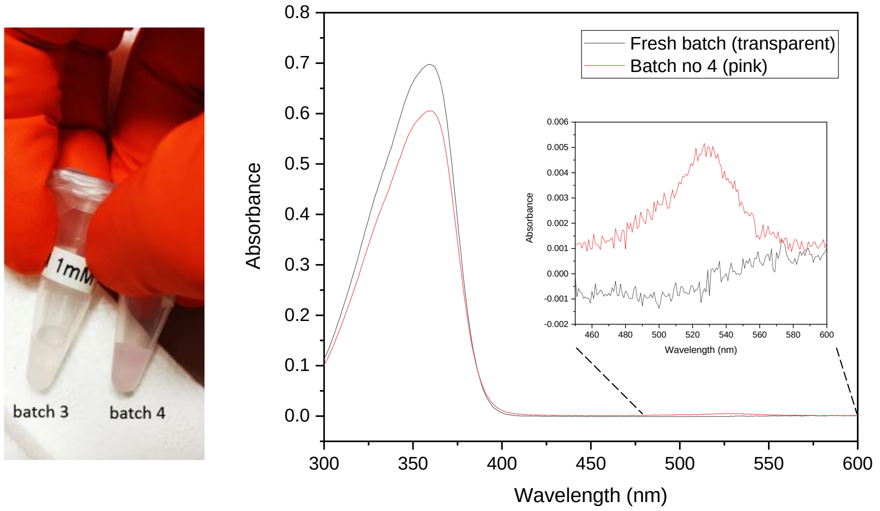


**Figure S3 –** Color change of the AFB1 batches. (left) Picture of the fresh ("batch 3") and old ("batch 4") AFB1 stocks. (right) Absorbance spectrum of the fresh and old AFB1 stocks. Both solutions present the 363 nm peak characteristic of AFB1, however the old solution also exhibits a peak around 530 nm in accordance with the visible pink color.

Although we have not found any relationship between the change of color and the composition of the toxin batch, we performed toxin characterization by high-performance liquid chromatography–tandem mass spectrometry (HPLC-MS) and noted the presence of AFB1 exo-8,9-epoxide (AFBO) for the colored batch and the absence of such compound for batches dissolved under inert atmosphere (**Figure S4**).

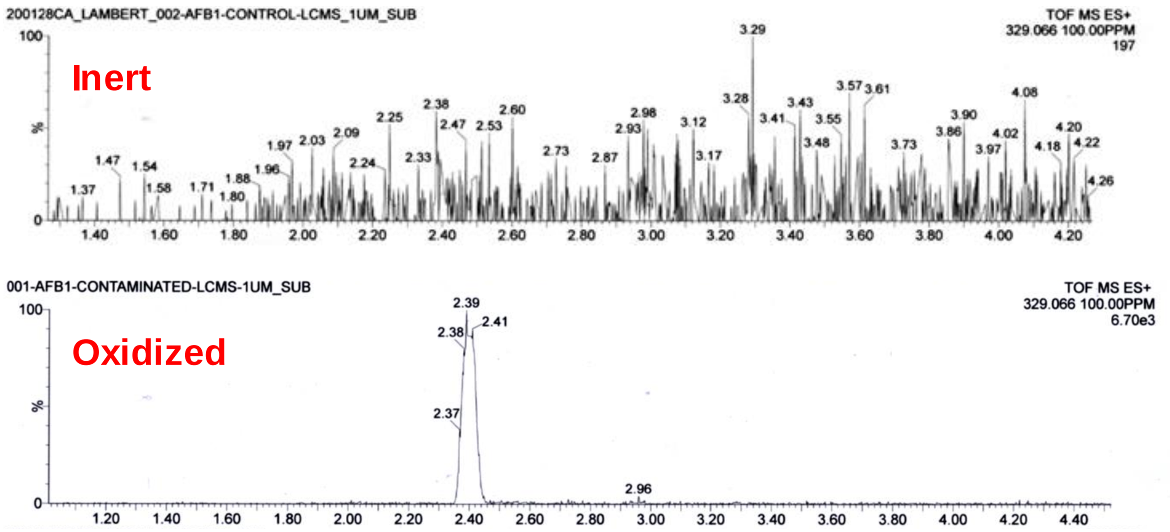


**Figure S4 -** HPLC-MS chromatograms of the AFB1 batches. Chromatograms for the aflatoxin B1 exo-8,9-epoxide ($C_{17}H_{12}O_7$) (m/z equal to 329.066) for (top) inert and (bottom) oxidized ("batch 4") batches of AFB1. The HPLC was performed on a $C_{18}$ column and the MS analysis by electrospray in positive mode.

In addition, we noted that the purity of the toxin batch can affect the response of the DNA-SWCNT sensors. We tested the Δ(AG) and A-2 sensors in presence of the oxidized batch and a batch dissolved under inert atmosphere (**Figure S5**). We observed that the response was higher in presence of the oxidized batch for the same concentration of toxin. In order to avoid possible misleading comparisons between two toxin batches of different purity, we indicate clearly in the figure legend if the result was obtained with a batch dissolved under non-controlled atmosphere. Furthermore, some results were obtained with toxins dissolved in methanol instead of DMSO. Indeed, while some toxins are more soluble in methanol than DMSO, DMSO was finally chosen in order to have a unique solvent for all toxins and limit further variations. We therefore also indicate in the figure legend if the toxin was dissolved in methanol instead of DMSO.

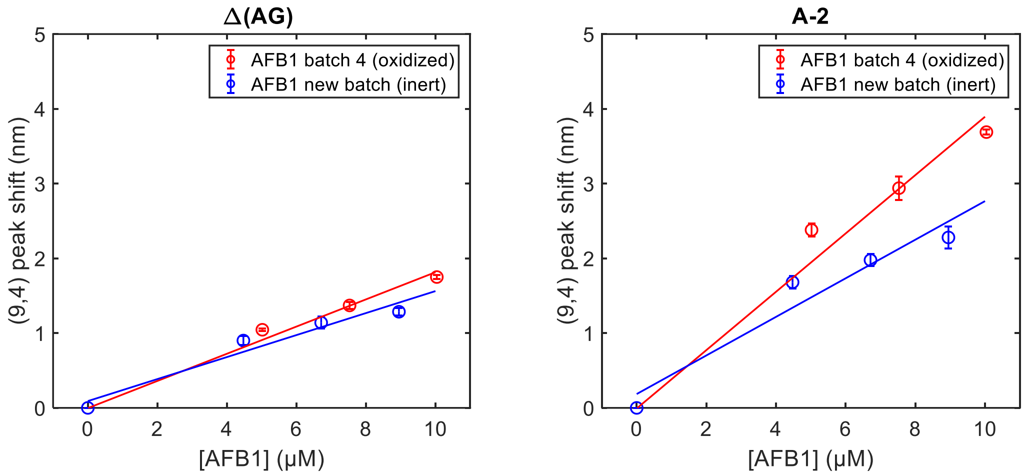


**Figure S5** - Response towards the oxidized AFB1. Peak positions of the (9,4) chirality for (left) the Δ(AG) and (right) the A-2 sensors in presence of inert (blue) and oxidized (red) AFB1 batches. The final AFB1 concentrations were comprised between 0 and 10 µM. All measurements were performed 200 min post-addition. The error bars represent 1 σ (n = 3). The calibration curves were fitted with linear fits.

While FB1 is freely soluble in water, AFB1 has a reported solubility limit of only approximately 60 µM in aqueous media. We tested this solubility limit by measuring the concentration of AFB1 in solution whether in DMSO or in a 1:100 DMSO:0.1 M NaCl to mimic our experimental conditions (**Figure S6**). For that, solutions of AFB1 were prepared by mixing 1 µL of AFB1 stock solution (10 or 20 mM in DMSO) in 99 µL of DMSO (blue) or 0.1 M NaCl (orange). The samples were incubated at room temperature for 1 h and centrifuged at 21,130 x g for 5 min in order to pellet possible aggregates. A small portion of the supernatant (20 µL) was taken, mixed with ethanol (180 µL) and the concentration was measured by absorbance spectroscopy using the extinction coefficient listed in the **Table S1**. We noted that the concentrations of the AFB1 solution in aqueous medium were not higher than 46 µM, therefore confirming a solubility limit of approximately 46 µM for AFB1 in our experimental conditions. This solubility limit was too low to reach the saturation of our sensors, preventing us from calculating an equilibrium dissociation constant for our sensors.

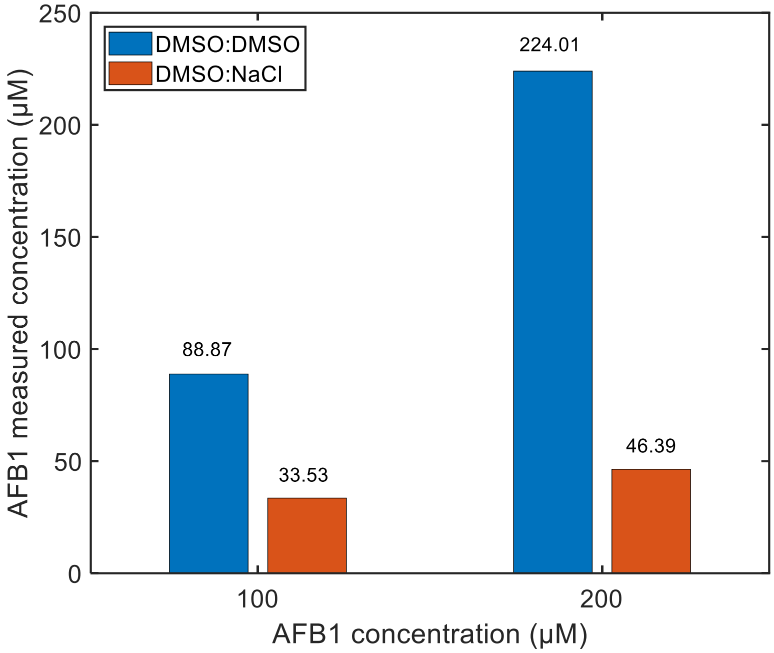


**Figure S6** - Solubility of the AFB1 solutions. Concentrations measured from absorbance spectroscopy for AFB1 diluted in DMSO (blue) or in 0.1 M NaCl (orange) as a function of the theoretical AFB1 concentrations.

#### Supplement 3: DNA library generation

The codes were written for Matlab R2017b.

Script for DNA mutagenesis (1 mutation)

```
%DNA creation program
%Benjamin Lambert (2019)
%Define 4 bases: A,C,G,T respectively as 1,2,3,4

%%%%%%%%%%%%%%%%%%%%%%%%%%%%%%
%LOADING INPUT DATA
%The sequence to mutate should be added in FASTA format in the file 'input_sequence.txt'

fid = fopen('input_sequence.txt'); %open the text file
data = textscan(fid,'%s%s%s');
fclose(fid);

DNA=zeros(size(str2mat(data{1}{2:2:end})')); %transforms A to 1, C to 2, G to 3 and T to 4
seq=size(DNA,2); %number of sequence in the input
length_seq=size(DNA,1); %length of the DNA sequences

k=2;
for i=1:seq
    for j=1:length_seq
        for  l=1:length(base)
            if data{1}{k}(j)=='A'
            DNA(j,i)=1;
            elseif data{1}{k}(j)=='C'
            DNA(j,i)=2;
            elseif data{1}{k}(j)=='G'
            DNA(j,i)=3;
            elseif data{1}{k}(j)=='T'
            DNA(j,i)=4;
            end
        end
    end
    k=k+2;
end

%%%%%%%%%%%%%%%%%%%%%%%%%%%%%%

tot_seq=length_seq*3; %total number of mutants for 1 mutation
mut_seq=zeros(length_seq,tot_seq);

for i=1:tot_seq
    mut_seq(:,i)=DNA(:,1);
end

%Do only 1 mutation
j=1;
for i=1:length_seq
    if DNA(i,1)==1
        mut_seq(i,j)=2;
        mut_seq(i,j+1)=3;
        mut_seq(i,j+2)=4;
    elseif DNA(i,1)==2
        mut_seq(i,j)=1;
        mut_seq(i,j+1)=3;
        mut_seq(i,j+2)=4;
    elseif DNA(i,1)==3
        mut_seq(i,j)=1;
        mut_seq(i,j+1)=2;
        mut_seq(i,j+2)=4;
    elseif DNA(i,1)==4
        mut_seq(i,j)=1;
```

```
        mut_seq(i,j+1)=2;
        mut_seq(i,j+2)=3;
    end
j=j+3;
end

%%%%%%%%%%%%%%%%%%%%%%%%%%%%%%
%mix the mutants randomly + transform in sequences

idx=randperm(tot_seq);
mut_seq_mix(:,:)=mut_seq(:,idx);

sequence_new=cell(length_seq,tot_seq);
sequence_new2=strings(tot_seq,1);

for j=1:tot_seq
    for i=1:length_seq
        if mut_seq_mix(i,j)==1
            sequence_new{i,j}="A";
        elseif mut_seq_mix(i,j)==2
            sequence_new{i,j}="C";
        elseif mut_seq_mix(i,j)==3
            sequence_new{i,j}="G";
        elseif mut_seq_mix(i,j)==4
            sequence_new{i,j}="T";
        end
    end
    sequence_new2(j)=strcat(sequence_new{:,j});
end

%%%%%%%%%%%%%%%%%%%%%%%%%%%%%%
%remove sequences that are not desired

fid2 = fopen('seq_old.txt');
data2 = textscan(fid2,'%s%s%s');
fclose(fid2);

for i=1:size(data2{1},1)
DNA_old(i,:)=data2{1}{i};
seq_old(i,:)=convertCharsToStrings(DNA_old(i,:));
end

sequence_new3=sequence_new2;
for l=1:length(sequence_new3)
    if sum(sequence_new3(l,:)==seq_old(:,:))>0
    sequence_new3(l,:)=NaN;
    end
end
sequence_new4=rmmissing(sequence_new3);

%%%%%%%%%%%%%%%%%%%%%%%%%%%%%%
%save the mutant sequences to a text file

fid=fopen(strcat(regexprep(data{1}{1}, '^.', '' ),'_mutant_sequences.txt'),'w');
for i=1:length(sequence_new4(:,1))
s3='%s\n';
formatSpec=strcat(s3);
fprintf(fid,formatSpec,sequence_new4{i,:});
end
fclose(fid);
```

<u>Script for DNA mutagenesis (1 to 3 simultaneous mutations)</u>

```
%DNA creation program
%Benjamin Lambert (2019)
%Define 4 bases: A,C,G,T respectively as 1,2,3,4


fid = fopen('input_sequence.txt');
data = textscan(fid,'%s%s%s');
fclose(fid);

fid = fopen('input_sequence.txt'); %open the text file
data = textscan(fid,'%s%s%s');
fclose(fid);

DNA=zeros(size(str2mat(data{1}{2:2:end})')); %transforms A to 1, C to 2, G to 3 and T to 4
seq=size(DNA,2); %number of sequence in the input
length_seq=size(DNA,1); %length of the DNA sequences

k=2;
for i=1:seq
    for j=1:length_seq
        for  l=1:length(base)
            if data{1}{k}(j)=='A'
            DNA(j,i)=1;
            elseif data{1}{k}(j)=='C'
            DNA(j,i)=2;
            elseif data{1}{k}(j)=='G'
            DNA(j,i)=3;
            elseif data{1}{k}(j)=='T'
            DNA(j,i)=4;
            end
        end
    end
    k=k+2;
end

%%%%%%%%%%%%%%%%%%%%%%%%%%%%%%

%number mutations
mut_id=3;

all_seq=3^mut_id*nchoosek(length_seq,mut_id); %total number of possibilities
number_seq=5000; %number of generated sequences

%%%%%%%%%%%%%%%%%%%%%%%%%%%%%%

ref_seq=zeros(number_seq,length_seq);

%All the sequences are the same as the input sequence
for i=1:number_seq
    ref_seq(i,:)=DNA(:,seq);
end

mut_seq=ref_seq;
for l=1:number_seq

nb=randperm(30);           %create an array of randomly permuted numbers between 1 and 30
pos1=datasample(nb,1);     %find a random position for the first mutation
if ref_seq(l,pos1)==3
    mut_seq(l,pos1)=datasample([1,2,4],1);
elseif ref_seq(l,pos1)==4
    mut_seq(l,pos1)=datasample([1,2,3],1);
elseif ref_seq(l,pos1)==1
    mut_seq(l,pos1)=datasample([2,3,4],1);
elseif ref_seq(l,pos1)==2
    mut_seq(l,pos1)=datasample([1,3,4],1);
end
```

```
if mut_id>1
nb(nb==pos1)=[];           %remove the position of mutation 1 from the list of possible positions
pos2=datasample(nb,1);     %find a random position for the second mutation
if ref_seq(l,pos2)==3
    mut_seq(l,pos2)=datasample([1,2,4],1);
elseif ref_seq(l,pos2)==4
    mut_seq(l,pos2)=datasample([1,2,3],1);
elseif ref_seq(l,pos2)==1
    mut_seq(l,pos2)=datasample([2,3,4],1);
elseif ref_seq(l,pos2)==2
    mut_seq(l,pos2)=datasample([1,3,4],1);
end
end

if mut_id>2
nb(nb==pos2)=[];           %remove the position of mutation 2 from the list of possible positions
pos3=datasample(nb,1);     %find a random position for the third mutation
if ref_seq(l,pos3)==3
    mut_seq(l,pos3)=datasample([1,2,4],1);
elseif ref_seq(l,pos3)==4
    mut_seq(l,pos3)=datasample([1,2,3],1);
elseif ref_seq(l,pos3)==1
    mut_seq(l,pos3)=datasample([2,3,4],1);
elseif ref_seq(l,pos3)==2
    mut_seq(l,pos3)=datasample([1,3,4],1);
end
end

if mut_id>3
disp('Please input a mutation rate between 1 and 3');
end

end

%%%%%%%%%%%%%%%%%%%%%%%%%%%%%%
%transform in sequences

sequence_new=cell(number_seq,length_seq);
for k=1:length(mut_seq(:,1))
    for i=1:length_seq
        if mut_seq(k,i)==1
            sequence_new{k,i}="A";
        elseif mut_seq(k,i)==2
            sequence_new{k,i}="C";
        elseif mut_seq(k,i)==3
            sequence_new{k,i}="G";
        elseif mut_seq(k,i)==4
            sequence_new{k,i}="T";
        end
    end
    sequence_new2(k,1)=strcat(sequence_new{k,:});
end

%%%%%%%%%%%%%%%%%%%%%%%%%%%%%%
%remove redundant sequences

sequence_new3=sequence_new2;
    if sum(sequence_new3(1,:)==sequence_new3([2:number_seq],:))>0
    sequence_new3(1,:)=NaN;
    end
for l=2:number_seq-1
    if sum(sequence_new3(l,:)==sequence_new3([1:l-1 l+1:number_seq],:))>0
    sequence_new3(l,:)=NaN;
    end
end
    if sum(sequence_new3(number_seq,:)==sequence_new3([1:number_seq-1],:))>0
    sequence_new3(number_seq,:)=NaN;
    end
sequence_new4=rmmissing(sequence_new3);
```

```
%%%%%%%%%%%%%%%%%%%%%%%%%%%%%%
%remove sequences that are not desired

fid2 = fopen('seq_old.txt');
data2 = textscan(fid2,'%s%s%s');
fclose(fid2);

for i=1:size(data2{1},1)
DNA_old(i,:)=data2{1}{i};
seq_old(i,:)=convertCharsToStrings(DNA_old(i,:));
end

sequence_new4b=sequence_new4;
for l=1:length(sequence_new4)
    if sum(sequence_new4b(l,:)==seq_old(:,:))>0
    sequence_new4b(l,:)=NaN;
    end
end
sequence_new4c=rmmissing(sequence_new4b);

%%%%%%%%%%%%%%%%%%%%%%%%%%%%%%
%save the mutant sequences to a text file

fid=fopen(strcat(regexprep(data{1}{1}, '^.', '' ),'_mutant_sequences.txt'),'w');
for i=1:length(sequence_new4(:,1))
s3='%s\n';
formatSpec=strcat(s3);
fprintf(fid,formatSpec,sequence_new4{i,:});
end
fclose(fid);
```

Script for DNA shuffling

(requires the custom package “npermutek”, see here:

https://www.mathworks.com/matlabcentral/fileexchange/69507-npermutek )

```
%DNA shuffling program
%Benjamin Lambert (2019)
%Define 4 bases: A,C,G,T respectively as 1,2,3,4

%%%%%%%%%%%%%%%%%%%%%%%%%%%%%%
%LOADING INPUT DATA
%The sequences used for the shuffling should be added in FASTA format in the file 'input_sequences.txt'

fid = fopen('input_sequences.txt'); %open the text file
data = textscan(fid,'%s%s%s');
fclose(fid);

base=unique(str2mat(data{1}{2:2:end})); %find the bases used
DNA=zeros(size(str2mat(data{1}{2:2:end})')); %transforms A to 1, C to 2, G to 3 and T to 4
num_seq=size(DNA,2); %number of sequences to shuffle
len_seq=size(DNA,1); %length of sequences

k=2;
for i=1:num_seq
    for j=1: len_seq
        for l=1:4
            if data{1}{k}(j)==base(l)
            DNA(j,i)=l;
            end
        end
    end
    k=k+2;
end

%%%%%%%%%%%%%%%%%%%%%%%%%%%%%%
%PARAMETERS%

cutting_size=5; %choose cutting size
tot_seq=100; %choose the total number of sequences to generate

%%%%%%%%%%%%%%%%%%%%%%%%%%%%%%
%Cutting of the sequences

if mod(len_seq,cutting_size)==0
DNA_cut=zeros(cutting_size+1,len_seq/cutting_size,num_seq);
p=0;
for k=1:num_seq
    m=0;
for i=1:size(DNA_cut,2)
    DNA_cut(1,i,k)=i+p; %labels of the fragments (from 1 to 6 for example)
    DNA_cut(2:cutting_size+1,i,k)=DNA(1+m:cutting_size+m,k); %contents of the fragment (bases contained in
the fragment)
    m=m+cutting_size;
end
p=p+size(DNA_cut,2);
end
else
disp('insert a fragment size that divides the sequence length');
end

%put all fragments all together
DNA_cut2=[DNA_cut(:,:,1) DNA_cut(:,:,2)];

%systematically combine the fragments
permut=npermutek(DNA_cut2(1,:),len_seq/cutting_size);
```

```
%%%%%%%%%%%%%%%%%%%%%%%%%%%%%%
%To remove some specific unwanted DNA fragments (uncomment this section if desired)

% range=[1 3 5]; %put here the unwanted fragments
%permut2=permut;
% for i=1:length(permut(:,1))
%     k=0;
%     for j=1:len_seq/cutting_size
%         if sum(ismember(range,permut(i,j)))>0
%             k=k+1;
%         end
%     end
%     if k>3
%         permut2(i,:)=NaN;
%     end
% end
% permut3=rmmissing(permut);
%%%%%%%%%%%%%%%%%%%%%%%%%%%%%%

if exist(permut3,'var') == 0
permut3=permut;
end

%%%%%%%%%%%%%%%%%%%%%%%%%%%%%%
%randomly pick a certain number of shuffled sequences from permut3 and assemble as sequences

DNA_shuffled=zeros(len_seq,tot_seq);
DNA_motif=zeros(len_seq/cutting_size,tot_seq);
row_id=datasample(1:length(permut3(:,1)),tot_seq,'Replace',false);

for m=1:tot_seq
index=permut3(row_id(m),:);
p=1;
for i=1:length(index)
    DNA_shuffled(p:p+(cutting_size-1),m)=DNA_cut2(2:cutting_size+1,index(i)); %final DNA sequences shuffled
    p=p+cutting_size;
end
    DNA_motif(:,m)=index; %organization of the DNA fragments as sequences (numer fragments)

end

sequence_new=cell(len_seq,tot_seq); %transforms numbers to letters
%numbers to string
for k=1:tot_seq
    for i=1:len_seq
        if DNA_shuffled(i,k)==1
            sequence_new{i,k}="A";
        elseif DNA_shuffled(i,k)==2
            sequence_new{i,k}="C";
        elseif DNA_shuffled(i,k)==3
            sequence_new{i,k}="G";
        elseif DNA_shuffled(i,k)==4
            sequence_new{i,k}="T";
        end
    end
    sequence_new2(k)=strcat(sequence_new{:,k}); %concatenate all letters into sequences
end


%%%%%%%%%%%%%%%%%%%%%%%%%%%%%%
%creates a file with all the shuffled sequences

fid=fopen(strcat('mutants_shuffled',num2str(cutting_size),'_',num2str(tot_seq),'.txt'),'w');
for i=1:length(sequence_new2(:,1))
s3='%s\n';
formatSpec=strcat(s3);
fprintf(fid,formatSpec,sequence_new2{i,:});
end
fclose(fid);
```

```
%%%%%%%%%%%%%%%%%%%%%%%%%%%%%%
%creates a file with all the shuffled fragments

fid=fopen(strcat('mutants_motif',num2str(cutting_size),'_',num2str(tot_seq),'.txt'),'w');
for i=1:tot_seq
s3='%s\n';
formatSpec=strcat(s3);
fprintf(fid,formatSpec,num2str(DNA_motif(:,i)'));
end
fclose(fid);
```

**Table S2 -** List of the main sequences discussed in this study. The red underlined letters represent the mutation compared to the parent sequence.

| Name | Sequence (5' to 3') |
|---|---|
| (AG)15 | AGAGAGAGAGAGAGAGAGAGAGAGAGAGAG |
| Δ(AG) | AGAGAGAGAGAGAGAGGAGAGAGCGGGAGAG |
| A-1 | AGAGAGAGAGAGAGAGGGAGTGAGCGGGAGAG |
| A-2 | AGAGAGAGAGAGAGAGGGAGTGAGCGGGAGGG |
| A-3 | TGAGAGAGAGAGAGAGGGAGTGAGCGGGAGGG |
| A-4 | TGAGAGAGAGAGGGGGAGTGAGCGGGAGGG |
| B-1 | AGAGAGAGGGAGAGGGAGAGAGCGGGAGAG |
| B-2 | AGAGAGAGGGAGAGGGAGGGAGCGGGAGAG |
| C-1 | AGCGAGAGAGAGAGGGAGAGAGCGGGAGAG |
| C-2 | AGCGAGAGAGAGAGGGAGAGAGCGGGAGCG |
| S5 | GAGGGGAGGGAGAGGGAGGGGAGAGGAGTG |
| Q-1 | GAGGGGAGGGAGAGGTAGGGGAGAGGAGTG |
| Q-2 | GAGGCGAGGGAGAGGTAGGGGAGAGGAGTG |
| Q-3 | GAGGCGTGGGAGAGGTAGGGGAGAGGAGTG |
| F-1 | GAGGGGAGGAAGAGGGAGGGGAGAGGAGTG |
| F-2 | GAGGGGAGAAAGAGGGAGGGGAGAGGAGTG |
| F-3 | GAGGGGAGAAAGAGGAAGGGGAGAGGAGTG |

### Supplement 4: DNA-SWCNT responses towards mycotoxins

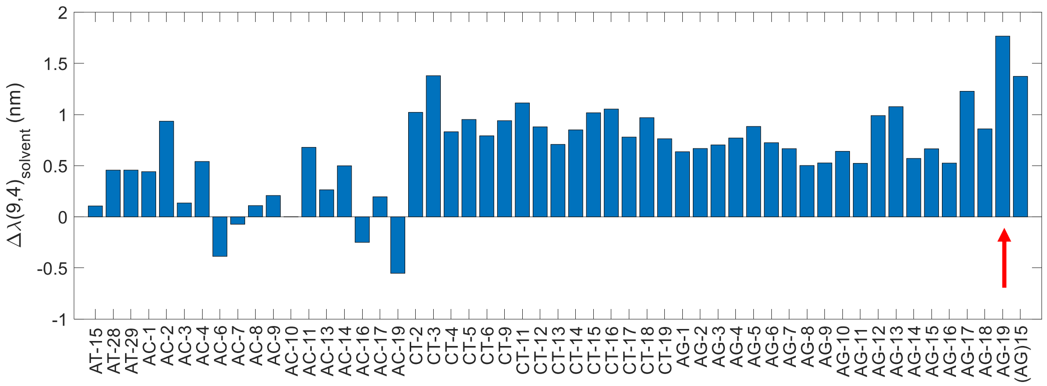


**Figure S7 -** Initial screening towards AFB1. Shifting of the (9,4) peak for 53 complexes from the first screening library towards AFB1 (10 µM final). The Δ(AG) mutant is indicated by the red arrow ("AG-19"). The measurements were taken after 20 min. The AFB1 batches prepared under non-inert conditions in DMSO.

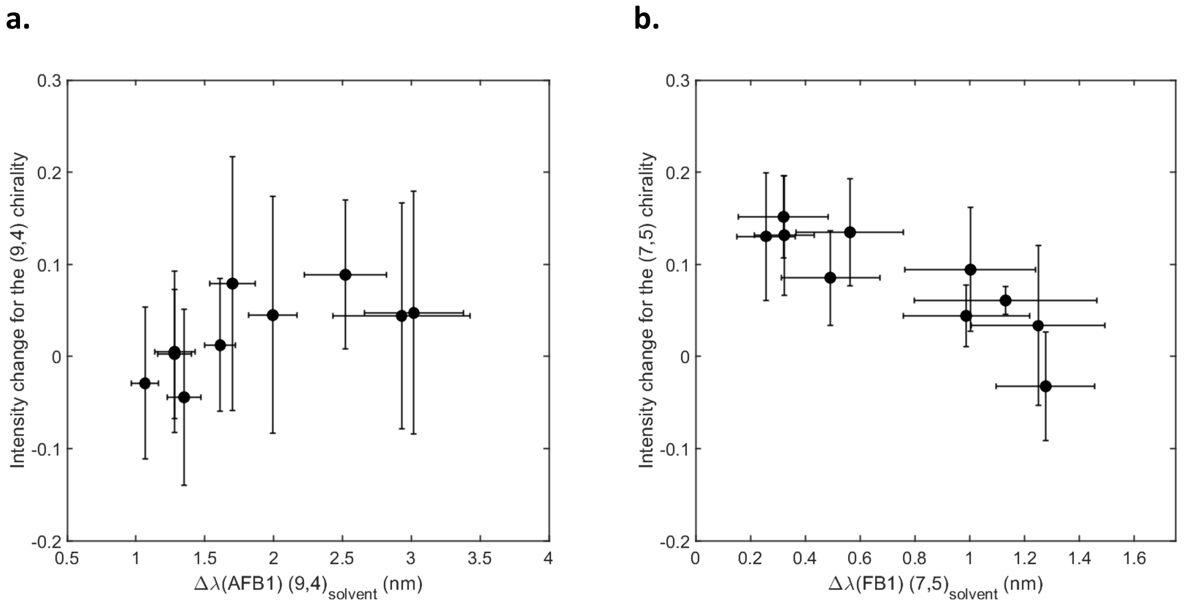


**Figure S8 -** Intensity change as a function of the shift in peak position (with respect to the solvent) for (a) the (9,4) chirality in presence of AFB1 and (b) the (7,5) chirality in presence of FB1. The intensity change corresponds to $(I-I_0)/I_0$ with I the intensity of the sensor in presence of toxin and $I_0$ the intensity of the sensor in the solvent. The toxins are at a final concentration of 5 µM. The error bars represent 1 σ (n = 4). The measurements are performed after a post-addition incubation time of 200 min.

**Table S3 -** Summary of the shifting responses (mean and standard deviation) in nm of the (9,4) chirality for the Δ(AG) mutants in presence of AFB1, FB1, OTA, ZEN and DON (all 10 μM) after a post-addition incubation time of 200 min.

| Name | AFB1 | FB1 | OTA | ZEN | DON |
|---|---|---|---|---|---|
| $(AG)_{15}$ | 0.78 ± 0.09 | -0.46 ± 0.04 | -0.06 ± 0.58 | 0.45 ± 0.03 | 0.02 ± 0.03 |
| Δ(AG) | 1.65 ± 0.16 | -0.07 ± 0.07 | -0.17 ± 0.05 | 0.69 ± 0.07 | 0.00 ± 0.06 |
| A-1 | 2.12 ± 0.33 | 0.08 ± 0.11 | -0.06 ± 0.08 | 0.85 ± 0.11 | 0.02 ± 0.04 |
| B-1 | 2.27 ± 0.26 | 0.18 ± 0.12 | 0.04 ± 0.10 | 0.82 ± 0.10 | 0.03 ± 0.06 |
| C-1 | 2.01 ± 0.16 | 0.13 ± 0.12 | 0.03 ± 0.04 | 0.85 ± 0.08 | 0.07 ± 0.03 |
| A-2 | 3.18 ± 0.29 | 0.59 ± 0.28 | 0.38 ± 0.10 | 1.30 ± 0.19 | 0.01 ± 0.07 |
| B-2 | 2.62 ± 0.24 | 0.23 ± 0.20 | 0.17 ± 0.14 | 0.98 ± 0.12 | 0.00 ± 0.11 |
| C-2 | 2.56 ± 0.22 | 0.23 ± 0.21 | 0.08 ± 0.17 | 1.10 ± 0.11 | 0.08 ± 0.14 |
| A-3 | 4.29 ± 0.46 | 0.43 ± 0.19 | 0.35 ± 0.12 | 1.55 ± 0.08 | 0.01 ± 0.13 |
| A-4 | 5.20 ± 0.71 | 0.39 ± 0.15 | 0.18 ± 0.15 | 1.74 ± 0.17 | -0.06 ± 0.10 |

**Table S4 -** Summary of the shifting responses (mean and standard deviation) in nm of the (7,5) chirality for for the Δ(AG) mutants in presence of AFB1, FB1, OTA, ZEN and DON (all 10 μM) after a post-addition incubation time of 200 min.

| Name | AFB1 | FB1 | OTA | ZEN | DON |
|---|---|---|---|---|---|
| $(AG)_{15}$ | 0.00 ± 0.07 | -0.49 ± 0.06 | -0.27 ± 0.21 | 0.02 ± 0.02 | -0.02 ± 0.02 |
| Δ(AG) | -0.07 ± 0.03 | 0.22 ± 0.11 | 0.16 ± 0.09 | 0.08 ± 0.02 | 0.03 ± 0.06 |
| A-1 | -0.19 ± 0.04 | 0.52 ± 0.22 | 0.44 ± 0.22 | 0.09 ± 0.08 | 0.03 ± 0.06 |
| B-1 | -0.06 ± 0.07 | 0.66 ± 0.27 | 0.52 ± 0.19 | 0.07 ± 0.07 | 0.04 ± 0.07 |
| C-1 | -0.03 ± 0.03 | 0.36 ± 0.16 | 0.28 ± 0.09 | 0.08 ± 0.03 | 0.03 ± 0.03 |
| A-2 | 0.17 ± 0.17 | 1.51 ± 0.27 | 1.18 ± 0.18 | 0.01 ± 0.11 | 0.10 ± 0.13 |
| B-2 | 0.18 ± 0.20 | 1.22 ± 0.28 | 0.95 ± 0.17 | 0.07 ± 0.07 | 0.01 ± 0.15 |
| C-2 | -0.12 ± 0.05 | 0.38 ± 0.16 | 0.29 ± 0.15 | 0.04 ± 0.03 | -0.01 ± 0.05 |
| A-3 | 0.20 ± 0.23 | 1.41 ± 0.39 | 1.10 ± 0.29 | -0.28 ± 0.51 | -0.09 ± 0.11 |
| A-4 | 0.52 ± 0.19 | 1.40 ± 0.23 | 0.93 ± 0.16 | 0.15 ± 0.12 | 0.02 ± 0.15 |

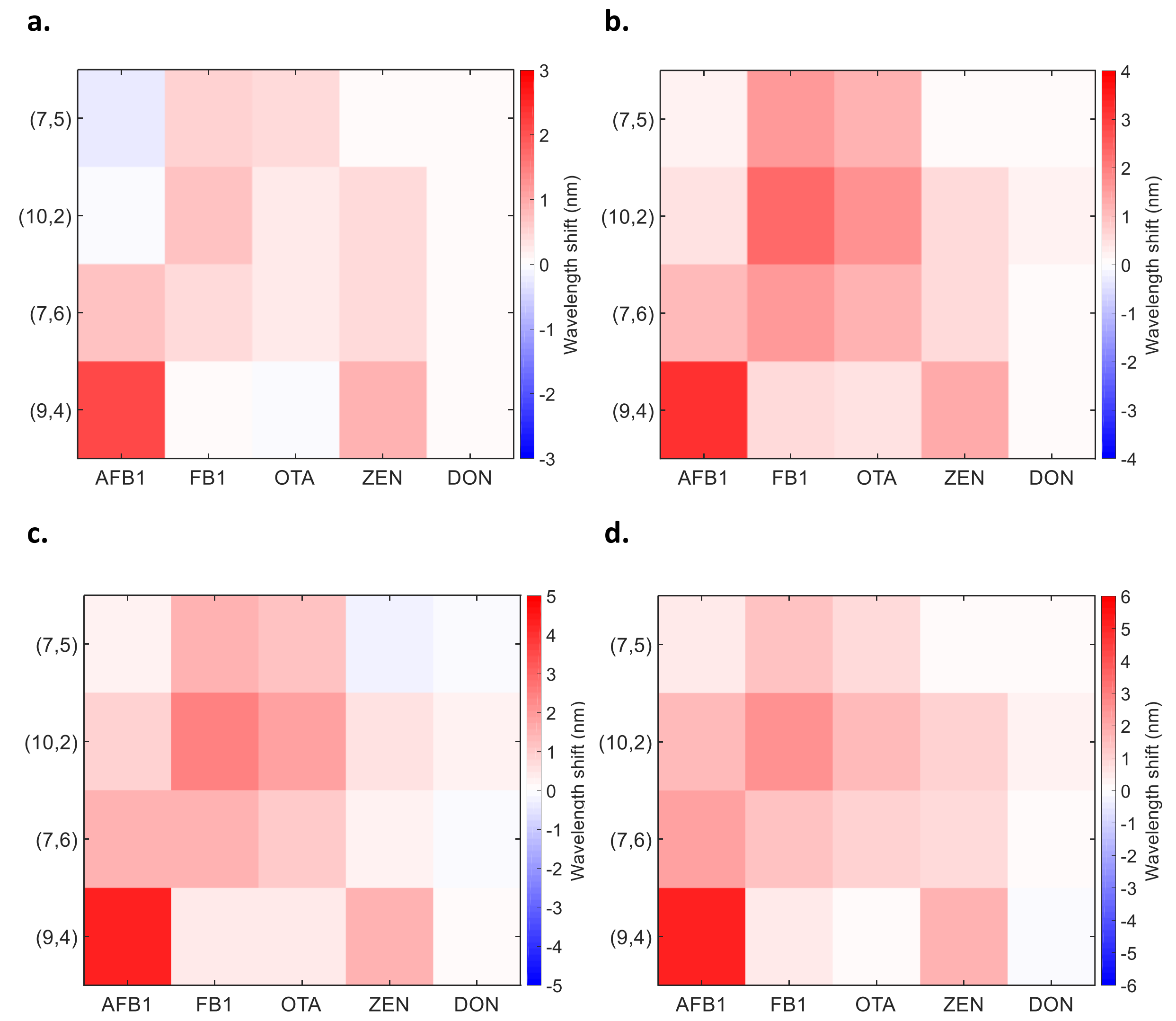


**Figure S9** - Response heatmaps of the (a) A-1, (b) A-2, (c) A-3 and (d) A-4 sensors in presence of AFB1, FB1, OTA, ZEN and DON (all 10 μM). The measurements are performed after a post-addition incubation time of 200 min.

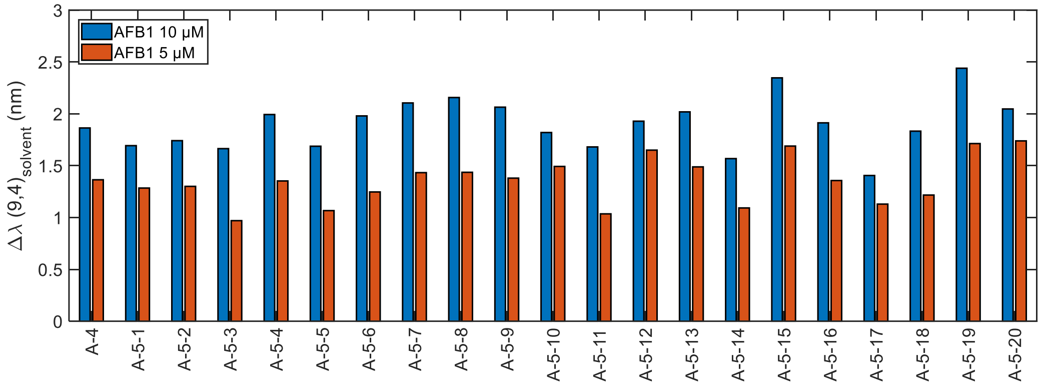


**Figure S10 -** Response of the mutants of the A-4 sensor towards AFB1. Shifting response of the (9,4) chirality in presence of AFB1 for the mutants of the fifth round. The measurements were performed 200 min after addition with AFB1 concentrations of 5 and 10 µM.

#### Supplement 5: Shuffled mutants

**a.**

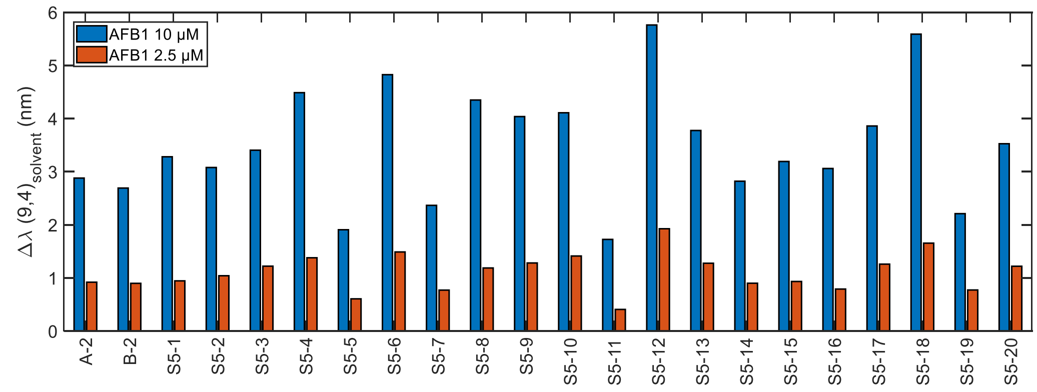


**b.**

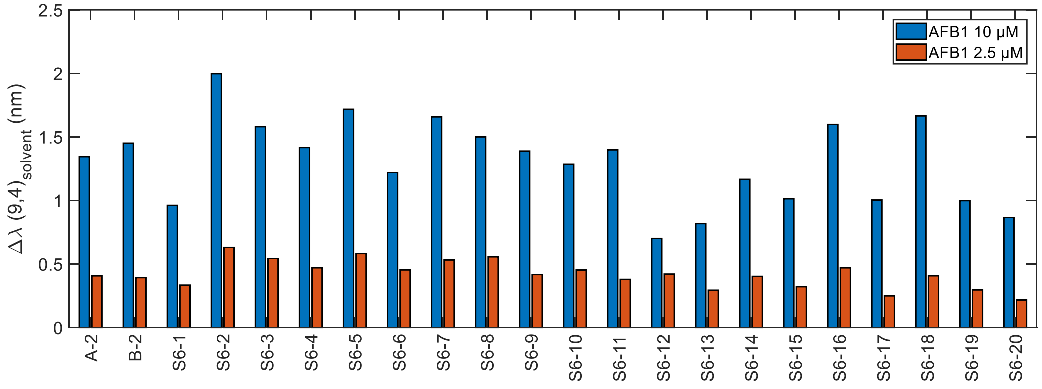


**c.**

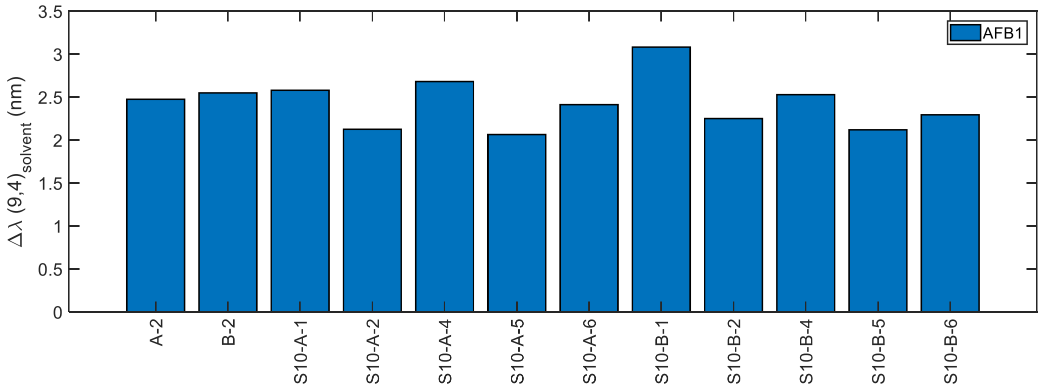


**Figure S11 -** Response towards AFB1 for the shuffled mutants. Shifting of the (9,4) peak for the shuffled mutants of the A-2 and B-2 sequences with cut size of (a) 5 and (b) 6 nucleotides. (c) Shuffled mutants of A-2 or B-2 individual sequences with cut size of 10 nucleotides. All measurements were performed 200 min post-addition. The AFB1 was at a final concentration of 10 µM (blue) or 2.5 µM (orange). The variations observed for the responses of A-2 and B-2 can be explained by variations of concentration between AFB1 batches. The AFB1 batches prepared under non-inert conditions in DMSO.

#### Supplement 6: Fluorescence intensity improvement

In order to counterbalance the decrease in starting intensity observed for the latest generations of mutants, we mutated the S5 sequence to find mutants exhibiting increased intensity of the (9,4) peak (**Figure S12**). In the first round, we found the Q-1 which exhibited an increased intensity (data not shown). In the second round, we identified the Q-2 sequence with increased intensity but also a decreased response. In the third round, we corrected the decrease in response previously observed for the Q-2 sequence with the Q-3 sequence which exhibited a similar response as the Q-1 but an increased intensity.

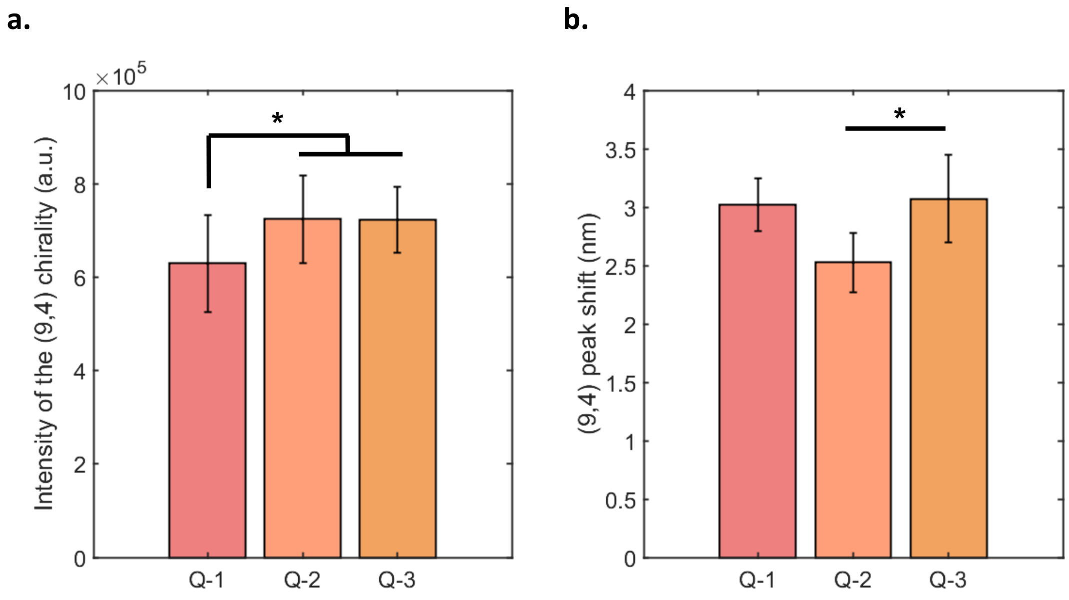


**Figure S12 -** Evolution of the fluorescence intensity mutants. (a) Initial intensity of the (9,4) peak for the Q-1, Q-2 and Q-3 sensors. The symbol * show a significant difference between the means for $p < 0.05$ (two-sample t-test, n = 18). The error bars represent 1 $\sigma$ (n = 18). (b) Shift of the (9,4) chirality in presence of AFB1 for the Q-1, Q-2 and Q-3 sensors. The AFB1 is at a final concentration of 10 μM. *$p < 0.05$ (two-sample t-test, n = 3). The error bars represent 1 $\sigma$ (n = 3). The measurements are performed after a post-addition incubation time of 200 min.

We replaced the DNA wrapping using sodium deoxycholate (SDC, final concentration 0.1%) in order to identify the origin of the increase in intensity (**Figure S13**), as previously demonstrated[2]. We collected fluorescence spectra after both 15 min and overnight incubation. We noted that the complete replacement only occurred after overnight incubation. Interestingly, the increase in intensity was retained post-SDC replacement, indicating that the intensity change is not due to a quantum yield effect but rather due to a change in the SWCNT chirality distribution.

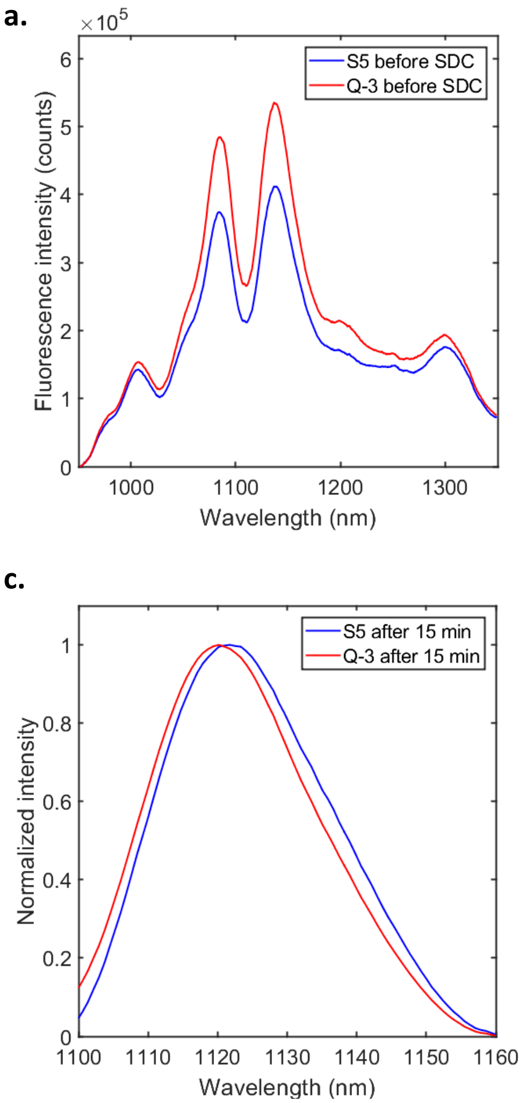


**Figure S13** - The differences of fluorescence intensity observed are not due a change in quantum yield but rather a result of changes in chirality distributions. Fluorescence spectra for the S5 (blue) and Q-3 (red) sensors (a) before and (b) after wrapping replacement with sodium deoxycholate (SDC). The "after o/n" measurements are performed after an overnight incubation (> 12 h). Normalized spectra of the (9,4) peak for the S5 (blue) and Q-3 (red) (c) 15 min after SDC addition and (d) after overnight incubation with SDC. The successful completion of the replacement is evidenced by the overlap of the peak positions after overnight incubation. The final concentration of SDC is 0.1%. The lines represent an average of 3 replicates.

#### Supplement 7: Analyte selectivity improvement

In addition to the correlation observed between the AFB1 response on the (9,4) chirality and the FB1 response on the (7,5) chirality, we observed a correlation between the (9,4) AFB1 and (10,2) FB1 response (**Figure S14**), as well as between the (9,4) peak position and the (9,4) AFB1 response.

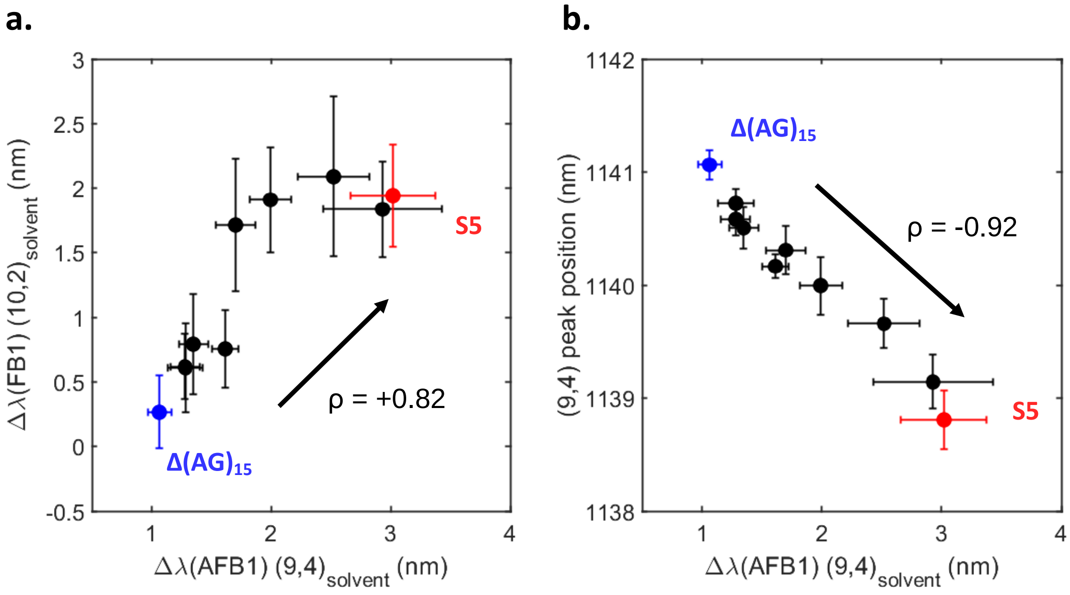


**Figure S14** - Correlation of the (a) (10,2) FB1 response (b) (9,4) peak position pre-addition with the (9,4) AFB1 response. Spearman correlation tests were done for both datasets revealing a positive ($\rho = +0.82$, $p < 0.05$) and negative ($\rho = -0.92$, $p < 0.05$) correlation for the (a) and (b) panels, respectively. Measurements were taken 200 min post-addition of the relevant toxin (5 μM). Error bars represent 1 σ standard error (n = 4).

The improved sensors for FB1 were evolved from the S5 sequence (**Figure S15**). In the first round, the response of the (7,5) and (10,2) peaks increased for the F-1 mutant compared to the S5 sequence (data not shown). While no improved FB1 sensitivity was obtained in the second round, the F-2 mutant exhibited a decreased response towards AFB1 for both the (7,5) and (10,2) chiralities. In the third round, we identified the F-3 mutant which both retained the decreased AFB1 response of the F-2 mutant and exhibited an increased FB1 response.

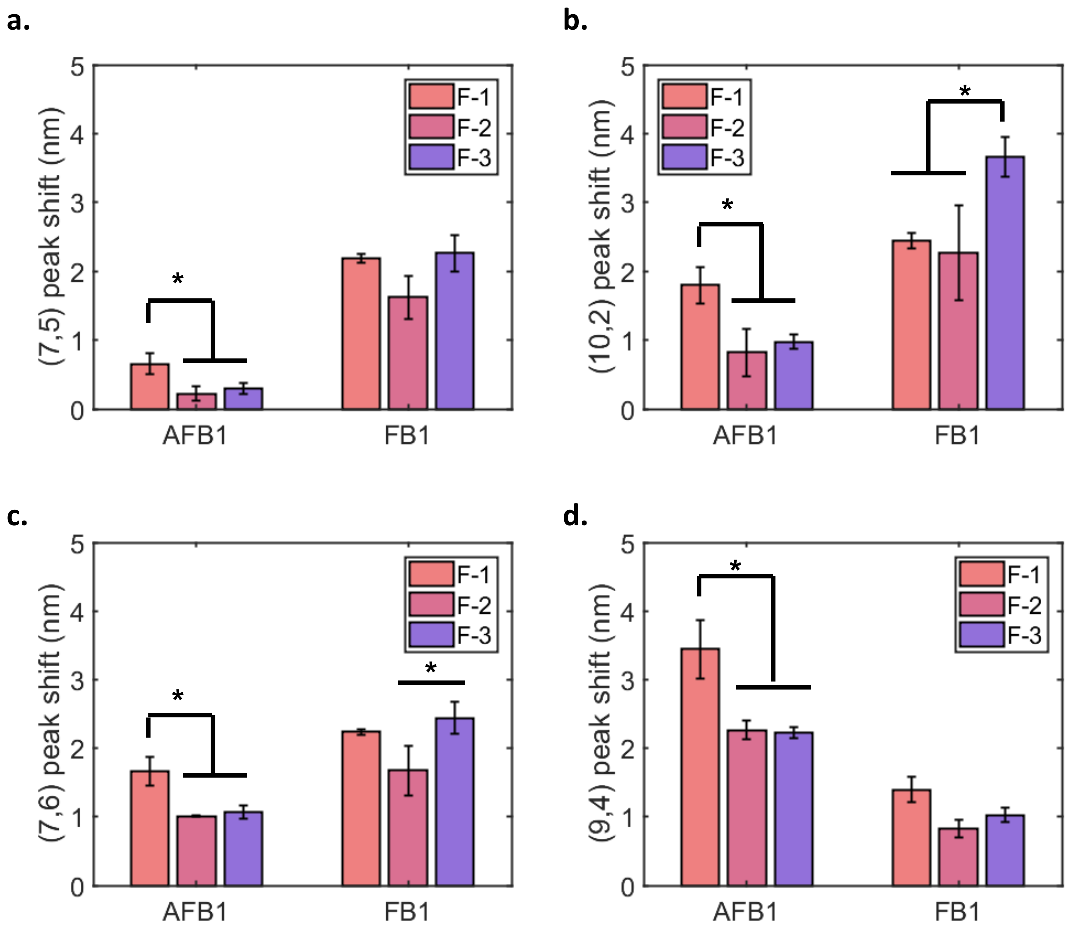


**Figure S15** - Shifting response of the F-1, F-2, and F-3 mutants in presence of AFB1 (10 µM) and FB1(10 µM) for the (a) (7,5), (b) (10,2), (c) (7,6), and (d) (9,4) chiralities. * $p < 0.05$ (two-sample t-test, $n = 3$). Error bars represent 1 σ standard error ($n = 3$). All measurements were performed 200 min post-addition.

As the F-3 sensor exhibits increased FB1 response and decreased AFB1 response, we observe an increased selectivity for the FB1 compared to the AFB1 (**Figure S16**). This effect is even more exacerbated for the (7,5) chirality for which the AFB1 response is almost zero.

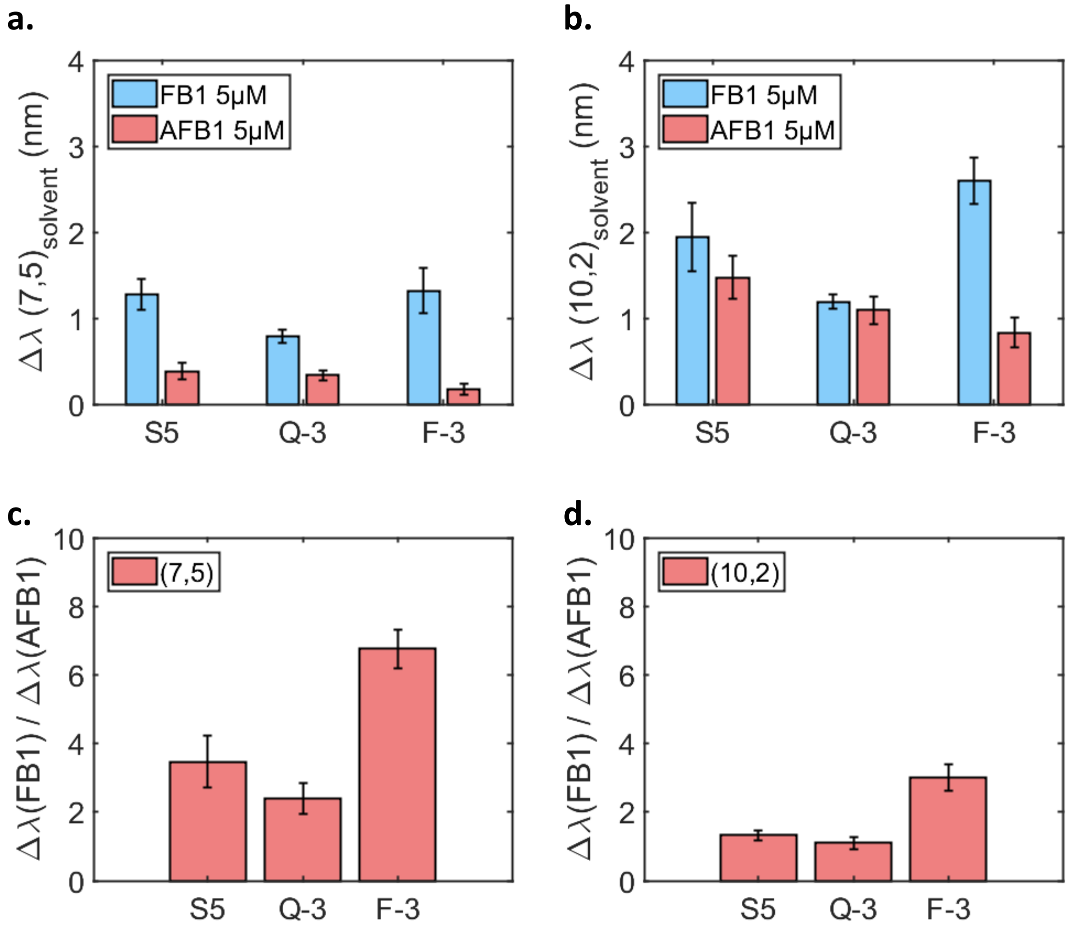


**Figure S16** - Selectivity of the FB1 response. Shifting response of the S5, Q-3, and F-3 sensors in presence of AFB1 (5 μM final concentration) and FB1 (5 μM final concentration) for the (a) (7,5) and (b) (10,2) chiralities. Ratios of the FB1 and AFB1 responses for the (c) (7,5) and (d) (10,2) chiralities. Error bars represent 1 σ standard error (n = 4). Measurements were performed 200 min post-addition.

The Q-3 and F-3 sensors both show an increased selectivity towards AFB1 versus FB1 compared to the S5 sensor (**Figure S17**). Although the Q-3 sensor exhibits a greater AFB1 response, it shows an AFB1 selectivity versus FB1 similar to the F-3 sensor for the (9,4) chirality. Yet, the Q-3 sensor also exhibits a greater AFB1 selectivity versus corn extract compared to the F-3 sensor for the (9,4) chirality, therefore making it a better AFB1 sensor overall.

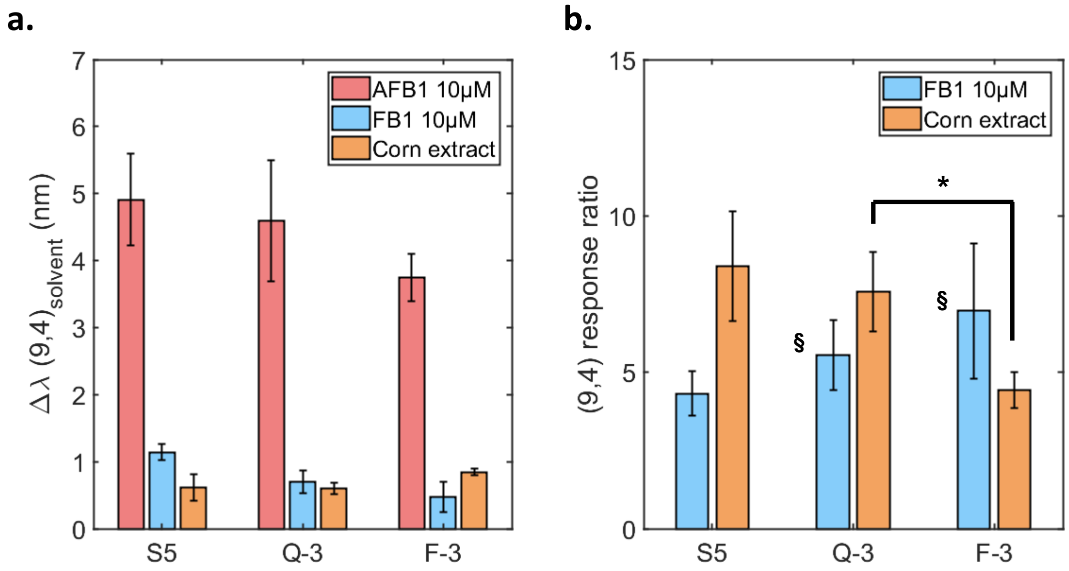


**Figure S17** - Selectivity of the AFB1 response. (a) Shifting response of the (9,4) chirality for the S5, Q-3, and F-3 sensors in presence of AFB1 (5 µM final concentration), FB1 (5 µM final concentration), and corn extract. (b) Ratios of the (9,4) chirality response to AFB1 versus the FB1 and corn extract responses (calculated as Δλ(AFB1)/Δλ(FB1) or Δλ(AFB1)/Δλ(corn)). All measurements were performed 200 min post-addition. *$p < 0.05$ (two-sample t-test, n = 4). Error bars represent 1 σ standard deviation (n = 4 for all except the bars indicated by § for which n=3).

#### Supplement 8: Influence of corn extract

The corn flour used in our study contains a variety of compounds including different proteins and starch. Even post-centrifugation, the supernatant still retained a strong yellow color indicating the presence of additional compounds, which have the potential of affecting the DNA-SWCNT response. We observed fluorescence shifting following the addition of corn extract for both the (7,5) and (9,4) chiralities (**Figure S18**). This shifting stabilized after approximately 320 min incubation time. Owing to this, all measurements performed in the corn matrix were incubated for a minimum of 320 min to allow sufficient time for the background to stabilize. In addition, the shifting responses towards the toxins were calculated with respect to the negative control containing corn extract.

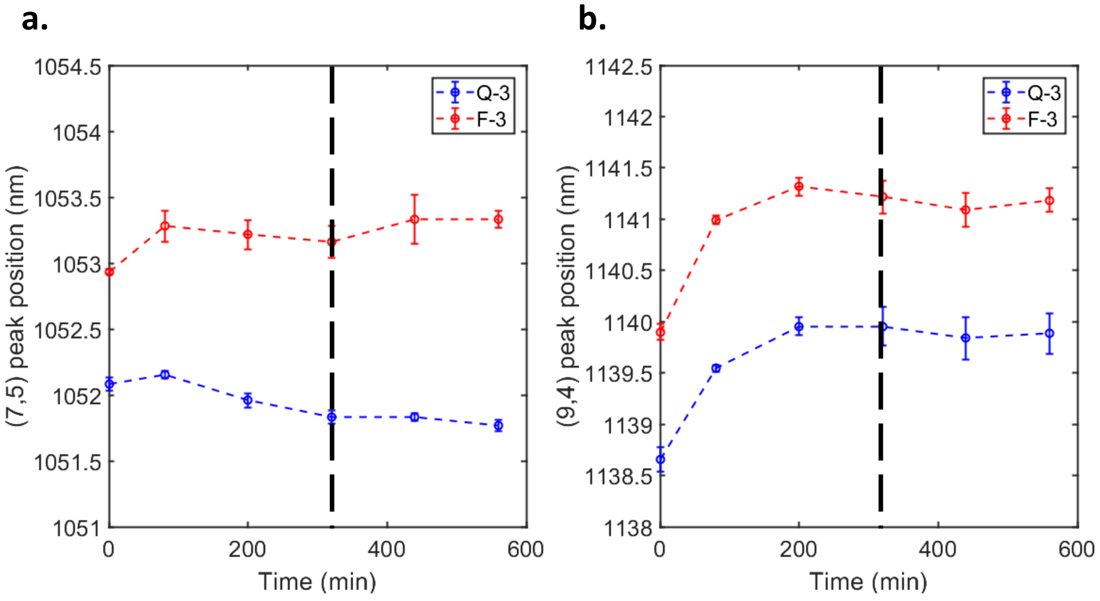


**Figure S18** - Evolution of the emission wavelength with the extracted corn blank in 60% methanol. Evolution of the (a) (7,5) and (b) (9,4) peak positions for the Q-3 (blue) and F-3 (red) sensors as a function of time. The peak position stabilized after 320 min (black horizontal line). The error bars represent 1 σ standard deviation (n = 3).

#### Supplement 9: Calibration curves and limit of detection

In our experiments, we fitted the calibration curve of the (7,5) chirality for the F-3 sensor towards FB1 with a linear fit as we were mainly interested by concentrations below 4 µM but noted that a single exponential fit could fit the response data better at concentrations above 4 µM (**Figure S19**).

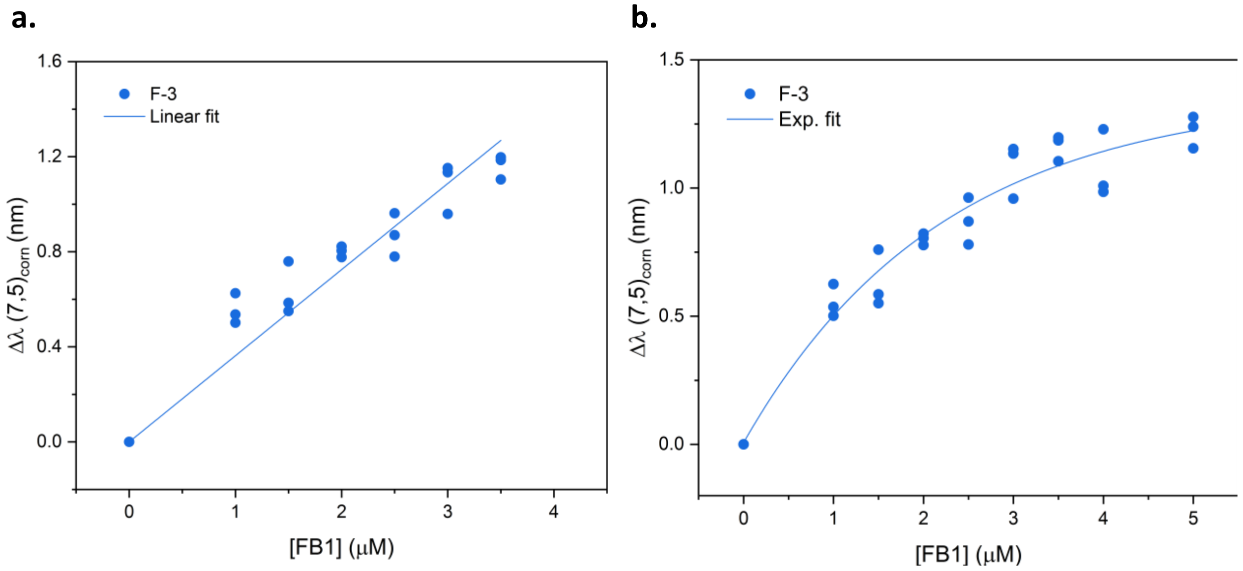


**Figure S19 -** Calibration curves for the (7,5) chirality of the F-3 sensor in presence of FB1 fitted with a (a) linear fit and (b) single exponential fit. All measurements were performed 320 min post-addition in the presence of corn in 60% methanol.

We used calibration curves performed in presence of corn extract to determine the limit of detection (LOD) of the Q-3 and F-3 sensors towards AFB1 and FB1, respectively. In our experiments, we obtained a peak position accuracy of approximately 0.2 nm. Following the addition of AFB1 and FB1 to the Q-3 and F-3 sensor, respectively, we observed a shift Δλ of ≥0.24 nm (AFB1 1 μM) and ≥0.35 nm (FB1 1 μM) versus the control, indicating that a LOD of 1 μM can be considered for both sensors in the presence of corn (**Figure S20**).

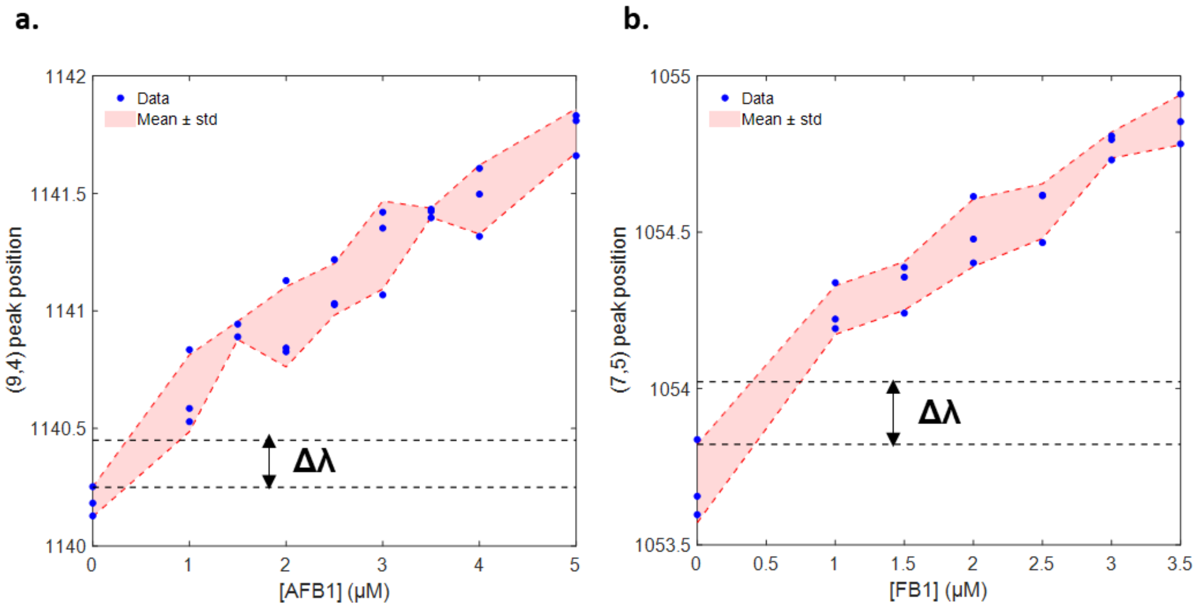


**Figure S20** - Limit of detection for the AFB1 and FB1 sensing. Calibration curves of the (a) (9,4) chirality (Q-3 sensor) towards AFB1 and (b) (7,5) chirality (F-3 sensor) towards FB1. The blue dots represent measured data and the red shaded area represents the mean of the data ± the standard deviation. Horizontal lines indicate the minimal shift Δλ necessary to identify a peak shift when compared to the negative control (0.2 nm). At a toxin concentration of 1 μM, the Q-3 sensor exhibits a shift ≥0.24 nm towards AFB1 and the F-3 sensor a shift ≥0.35 nm towards FB1. All measurements were recorded 320 min post-addition incubation in presence of corn in 60% methanol.

#### Supplement 10: Response in presence of other food products

We tested the performance of the S5 and Q-3 sensors towards the mycotoxins in presence of almond flour extract (**Figure S21**). We noted that the presence of almond extract had almost no effect on the position of the (9,4) chirality, therefore allowing an efficient sensing of AFB1 in almond products.

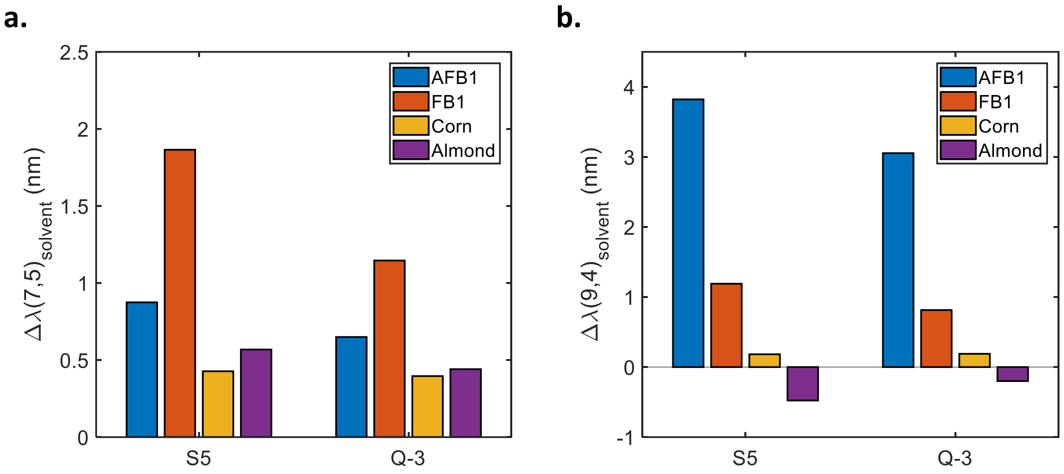


**Figure S21 -** Response of the sensors in presence of almond extract. Shifting response of (a) the (7,5) and (b) the (9,4) chiralities in presence of AFB1, FB1, corn flour extract and almond flour extract. The measurements were performed with a toxin concentration of 5 µM and 200 min after addition.

In addition, we tested the mycotoxin response of the S5, Q-3 and F-3 sensors in presence of beer (**Figure S22**). We observed a large red-shifting upon addition for both the (7,5) and (9,4) chiralities, preventing the detection of toxins in this medium with our current sensors.

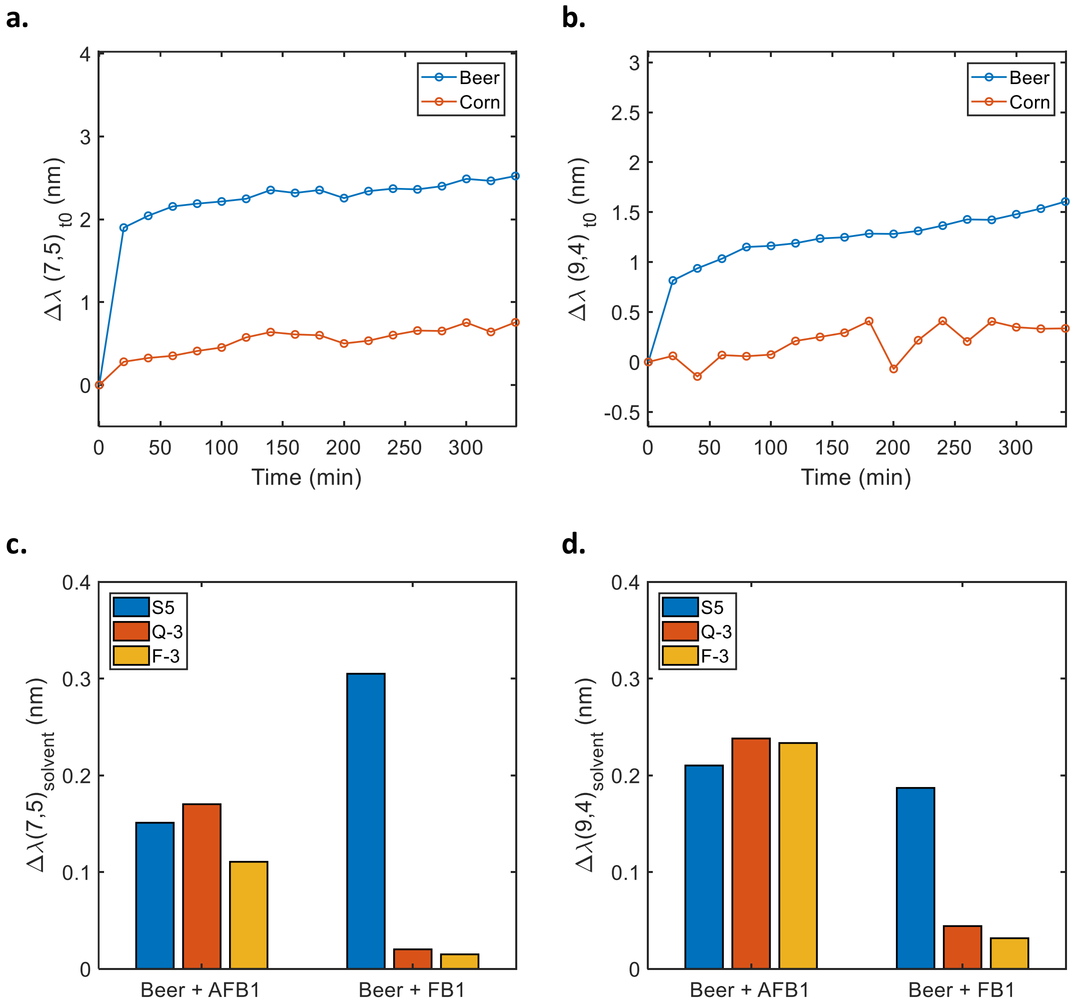


**Figure S22** - Response of the sensors in presence of beer. (a,b) Shifting of the (7,5) and (9,4) chiralities caused by corn extract (red) and beer (blue) as a function of time for the S5 sensor. (c,d) Shifting response of the (7,5) and (9,4) chiralities towards AFB1 and FB1 in presence of beer for the S5, Q-3 and F-3 sensors. The measurements were performed with a toxin concentration of 5 µM and 200 min after addition.

#### Supplement 11: Evaluation of the selectivity of the sensors

The selectivity of the responses was determined by differences between the response towards a single toxin versus towards the mixture of toxins. The (9,4) chirality of the Q-3 sensor showed no significant difference for the response towards AFB1 or the mixture of AFB1 and FB1. Similarly, the (7,5) chirality of the F-3 sensor exhibited no significant response difference towards FB1 or the mixture of toxins.

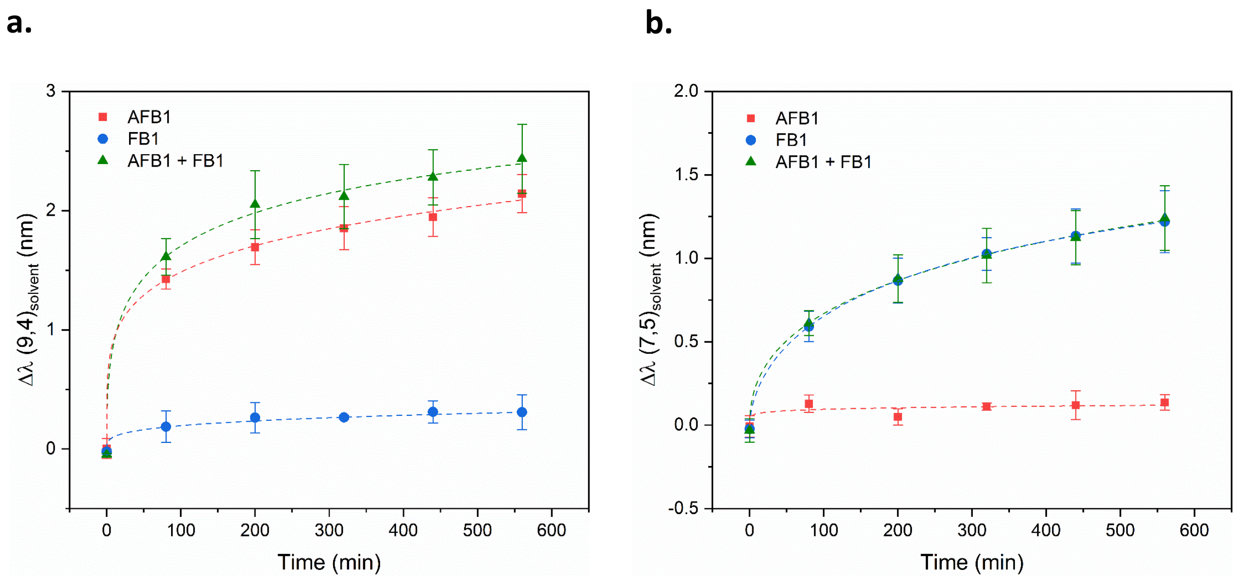


**Figure S23 -** The (9,4) and (7,5) chirality response of the (c) Q-3 and (d) F-3 sensors, respectively, towards AFB1 (2.5 µM, red), FB1 (2.5 µM, blue), and the toxin mixture (2.5 µM of each toxin, green). Error bars represent 1 σ standard error (n = 3). All measurements were recorded 320 min post-addition.

The sensors were tested in the presence of extracted corn matrix spiked with AFB1 and FB1. We first verified that the toxins were not affected by the extraction and purification procedure, such as the effect of the PTFE filters (**Figure S24**). Concentrations of toxins in spiked corn samples were determined by fluorescence using our sensors (**Figure S25**) and verified using HPLC-MS (**Table S5**). These results confirmed that the Q-3 sensor was performing better for AFB1 detection by monitoring the (9,4) chirality and the F-3 sensor for FB1 detection by monitoring the (7,5) chirality.

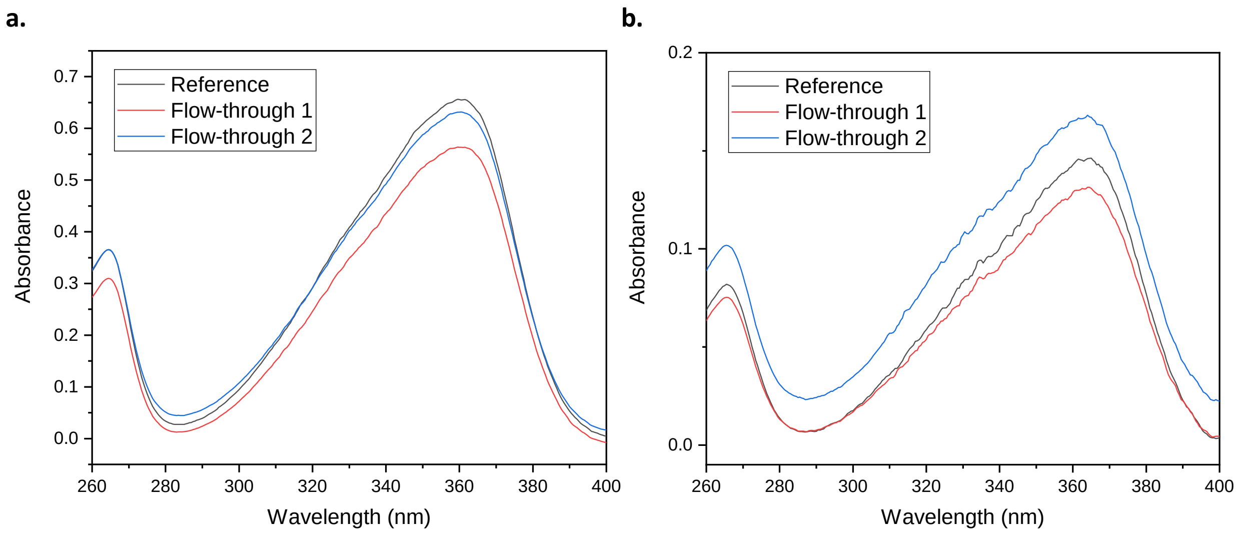


**Figure S24 -** Purification of AFB1 samples through PTFE filters. Absorbance spectra of the AFB1 in (a) 100% methanol and (b) 60% methanol before (reference) and after filtration through 0.22 µm PTFE filter. This control confirms that the purification of spiked corn samples with PTFE filters does not affect the final concentration of ABF1.

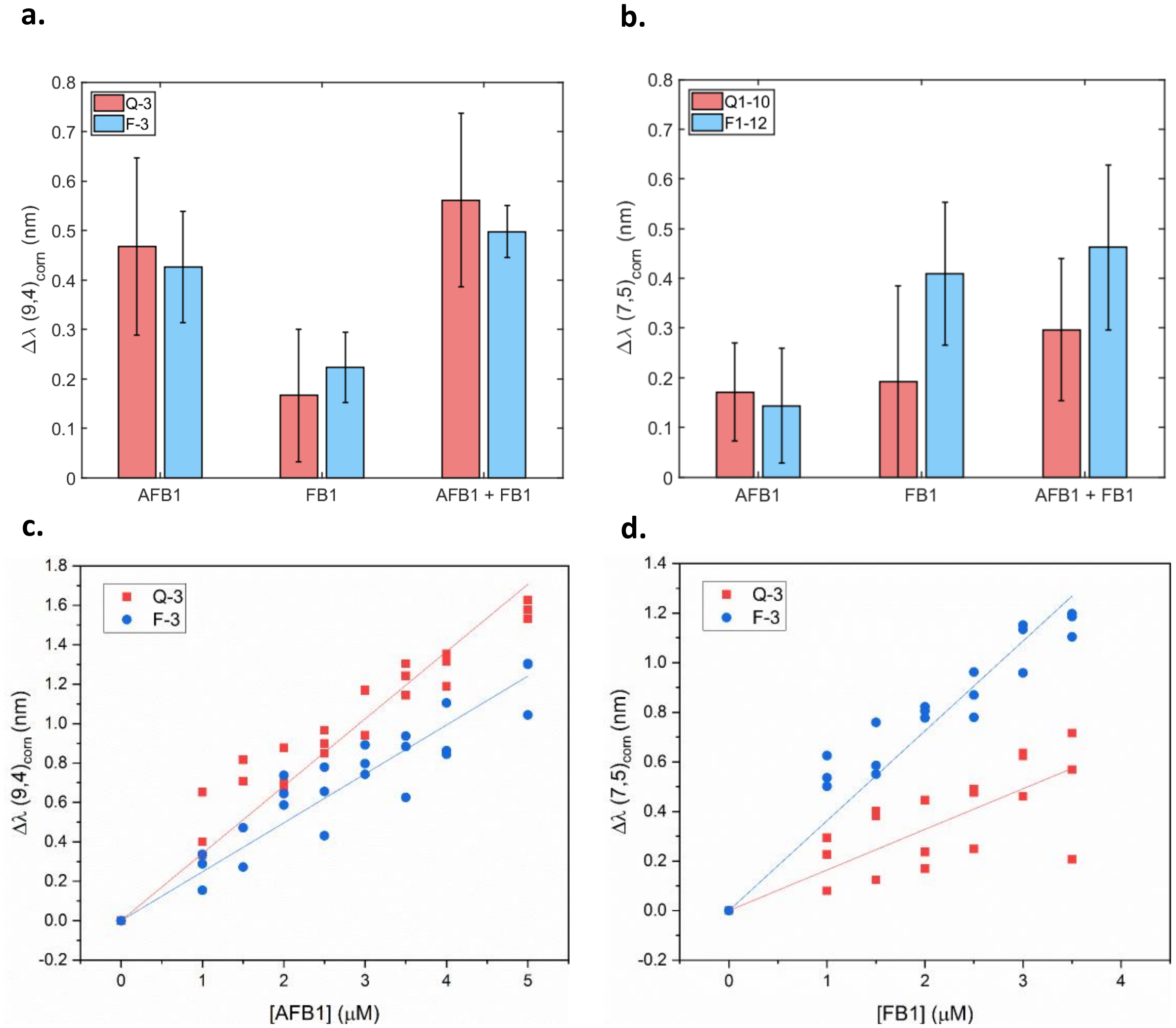


**Figure S25** - Detection of AFB1 and FB1 in extracted spiked corn. Wavelength shift of the (a) (9,4) and (b) (7,5) chiralities following the addition of AFB1 (2.5 μM), FB1 (2.5 μM), or AFB1 and FB1 (2.5 μM of each toxin) for the Q-3 (red) and F-3 (blue) sensors. Error bars represent 1 σ standard deviation (n = 6). Calibration curves for the (c) (9,4) chirality towards AFB1 and (d) (7,5) chirality towards FB1 for the Q-3 (red) and F-3 (blue) sensors. All measurements were performed 320 min post-addition.

**Table S5 -** Determination of toxin concentration in corn. The slope refers to the linear fit of the calibration curves shown in **Figure S25**. The shifting response of the sensor is given here for the (9,4) chirality (AFB1) and (7,5) chirality (FB1). The concentration values were calculated from the slope and compared to values measured by HPLC-MS. Red indicates where there was too high error or disagreement between the concentrations measured by fluorescence and HPLC-MS. Green indicates good agreement between the concentrations measured by fluorescence and HPLC-MS.

| Chirality | Sensor | Sample | Slope (nm.μM$^{-1}$) | Response (nm) | Calculated (μM) | Measured (μM) |
|---|---|---|---|---|---|---|
| (9,4) - AFB1 | Q-3 | AFB1 | $a = 0.34 \pm 0.01$ | $0.47 \pm 0.18$ | $1.37 \pm 0.53$ | $1.63 \pm 0.06$ |
| | | AFB1 + FB1 | $a = 0.34 \pm 0.01$ | $0.56 \pm 0.18$ | $1.64 \pm 0.52$ | $1.53 \pm 0.15$ |
| | F-3 | AFB1 | $a = 0.25 \pm 0.01$ | $0.43 \pm 0.11$ | $1.72 \pm 0.46$ | $1.63 \pm 0.06$ |
| | | AFB1 + FB1 | $a = 0.25 \pm 0.01$ | $0.49 \pm 0.05$ | $2.01 \pm 0.22$ | $1.53 \pm 0.15$ |
| (7,5) - FB1 | Q-3 | FB1 | $a = 0.16 \pm 0.01$ | $0.19 \pm 0.19$ | $1.17 \pm 1.18$ | $1.28 \pm 0.12$ |
| | | AFB1 + FB1 | $a = 0.16 \pm 0.01$ | $0.29 \pm 0.14$ | $1.81 \pm 0.88$ | $1.26 \pm 0.12$ |
| | F-3 | FB1 | $a = 0.36 \pm 0.01$ | $0.41 \pm 0.14$ | $1.13 \pm 0.53$ | $1.28 \pm 0.12$ |
| | | AFB1 + FB1 | $a = 0.36 \pm 0.01$ | $0.46 \pm 0.17$ | $1.28 \pm 0.61$ | $1.26 \pm 0.12$ |

#### Supplement 12: Multimodal sensing of mycotoxins

To further improve the selectivity of the mixed sensors, we therefore wrapped HiPco SWCNTs (rich in the (9,4) chirality) with the Q-3 sequence and CoMoCAT SWCNTs (rich in the (7,5) chirality) with the F-3 sequence to bias the distribution of chiralities towards their respectively engineered wrapping in the final mixture (**Figure S26**). We were able to confirm an uncompromised (9,4) response towards AFB1 even in the presence of the residual F-3 wrapped (9,4) SWCNTs from the CoMoCAT solution in the mixture. By contrast, the (7,5) sensitivity towards the FB1 response was still affected by the presence contaminating Q-3 wrapped (7,5) SWCNTs, as HiPco SWCNTs contain a substantial portion of (7,5) SWCNTs.

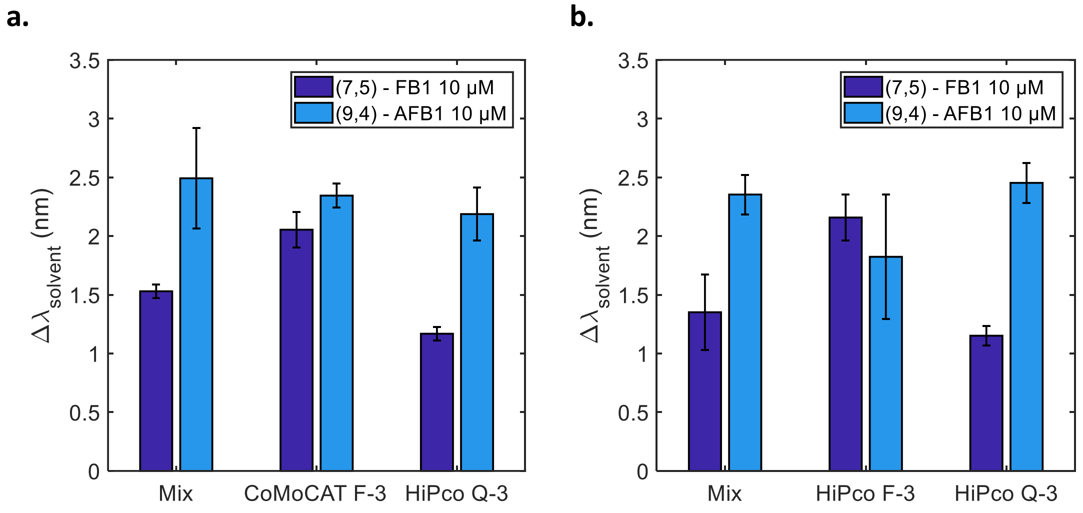


**Figure S26** - Comparison of the mixtures made of HiPco and HiPco-CoMoCAT sensors. Shifting response of the (7,5) (dark blue) and (9,4) (light blue) chiraclities for (a) the HiPco-CoMoCAT and (b) the HiPco sensors in presence of FB1 and AFB1. The measurements were performed 200 min after addition and with a toxin concentration of 10 µM. The use of CoMoCAT for the FB1 sensors does not strongly improve the selectivity of the mixed sensors compared to the individual sensors.

#### Supplement 13: Impact of preparation method on the response behaviour of the DNA-SWCNT sensors

We compared the AFB1 and FB1 responses for the Δ(AG), A-4, and S5 sensors prepared by both surfactant exchange and direct sonication (**Figure S27**). The sonicated samples and exchanged samples were both able to sense AFB1, however the response of the sonicated A-4 sensor was reduced. On the other hand, no FB1 response was observed for sensors prepared by sonication.

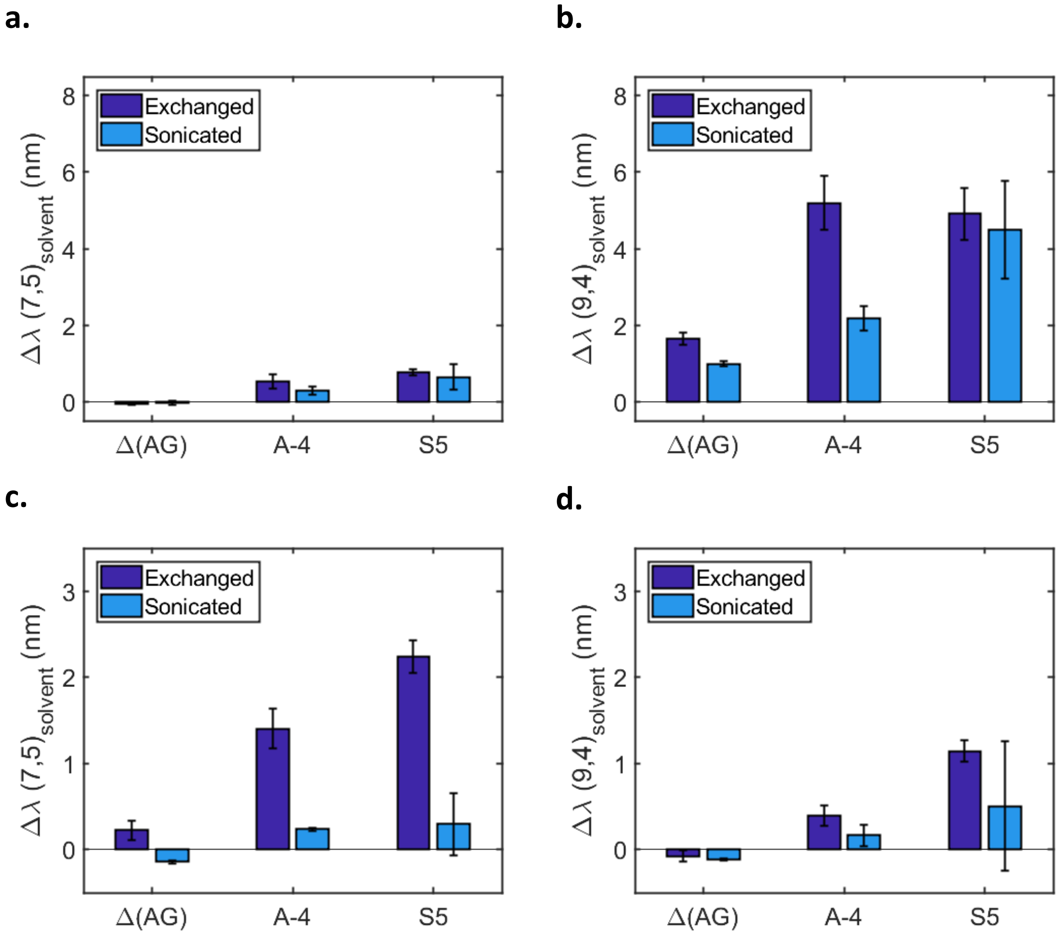


**Figure S27** - Wavelength shifting response of the (7,5) and (9,4) chiralities for the Δ(AG), A-4, and S5 sensors prepared by exchange or direct sonication towards (a,b) AFB1 (10 µM) and (c,d) FB1 (10 µM). All measurements were performed 200 min post-addition. The error bars represent 1 σ standard deviation (n = 3).

#### Supplement 14: NIR absorbance of the sensors

The near-infrared (NIR) absorbance of the Q-3 and F-3 sensors was monitored in presence of AFB1 and FB1 and compared to the solvent control (**Figure S28**). In the presence of AFB1, the shifting response of the (9,4) peak was convoluted with the (7,6) peak, however we still noted a broadening of the peak at 1150 nm, which could be indicative of the shifting response. In the presence of FB1, a clear shift of the (7,5) peak was observed (indicated by the black dotted line).

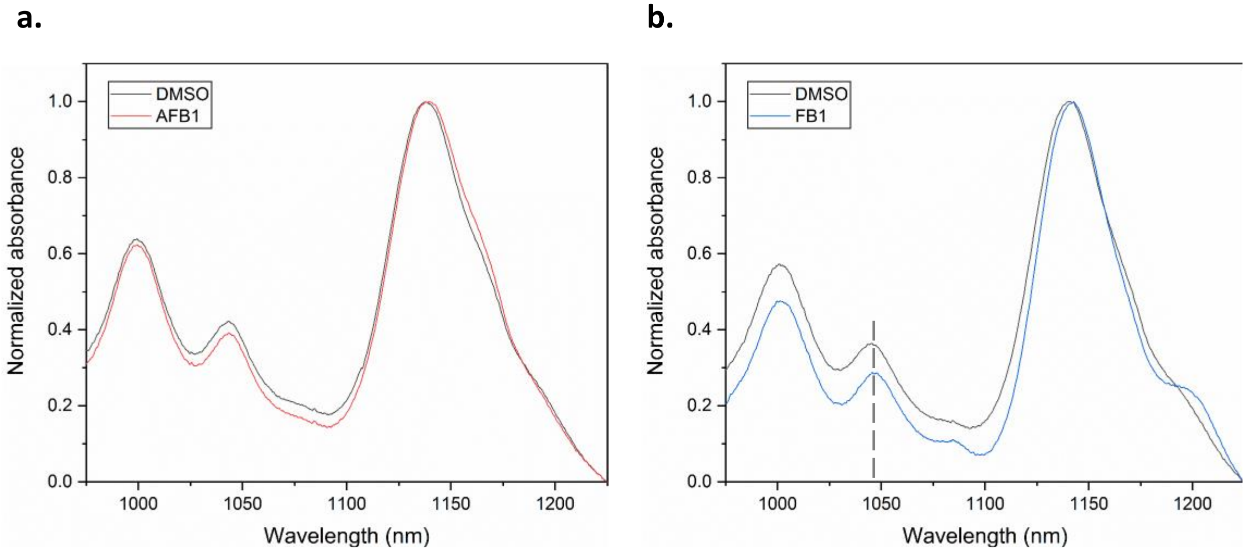


**Figure S28** - Absorbance spectra of the (a) Q-3 and (b) F-3 sensors in presence of 10 μM AFB1 (red) and FB1 (blue), as well as DMSO (black).

#### Supplement 15: Toxin removal

We initially attempted to study the reversibility of the immobilized DNA-SWCNTs on a glass substrate, but the sensors exhibited a loss of responsivity when immobilized (**Figure S29**).

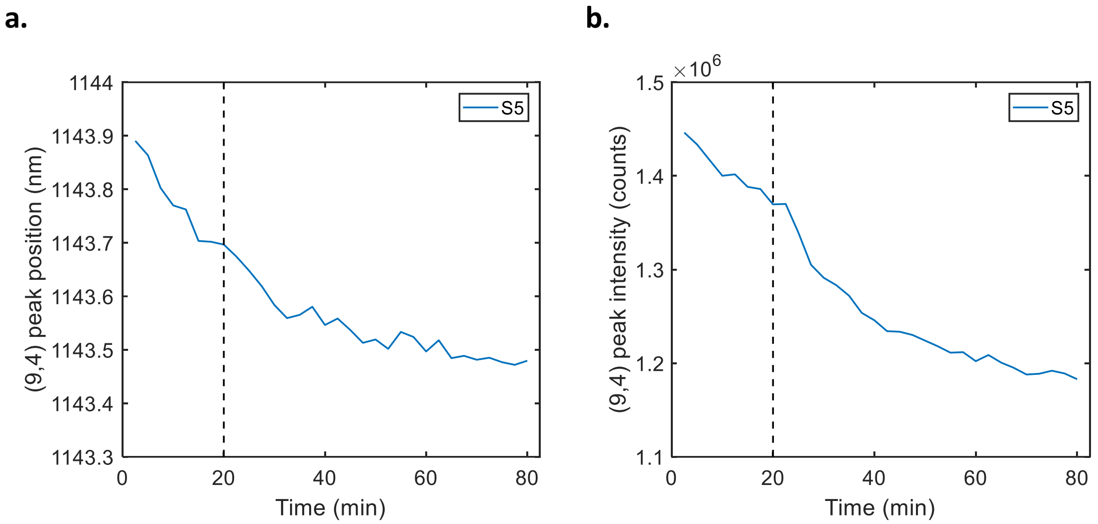


**Figure S29 -** Response of immobilized sensors towards AFB1. (a) Peak position and (b) peak intensity of the (9,4) chirality for the immobilized S5 sensor as a function of time in presence of 0.1M NaCl (0 to 20 min) and in presence of 20 µM AFB1 (20 to 80 min).

We ultimately developed a liquid-state washing protocol, illustrated in **Figure S30**, that showed improved AFB1 sensitivity compared to immobilized samples. Briefly, reacted DNA-SWCNT sensors were incubated overnight with the toxin. The DNA-SWCNTs were then precipitated using a salt-assisted ethanol precipitation and resuspended in 0.1 M NaCl (as described in the **Materials and methods**

). The response of the (9,4) and (7,5) chiralities were then tested following both AFB1 and FB1 addition for the resuspended sensors.

<u>Standard washing protocol</u>

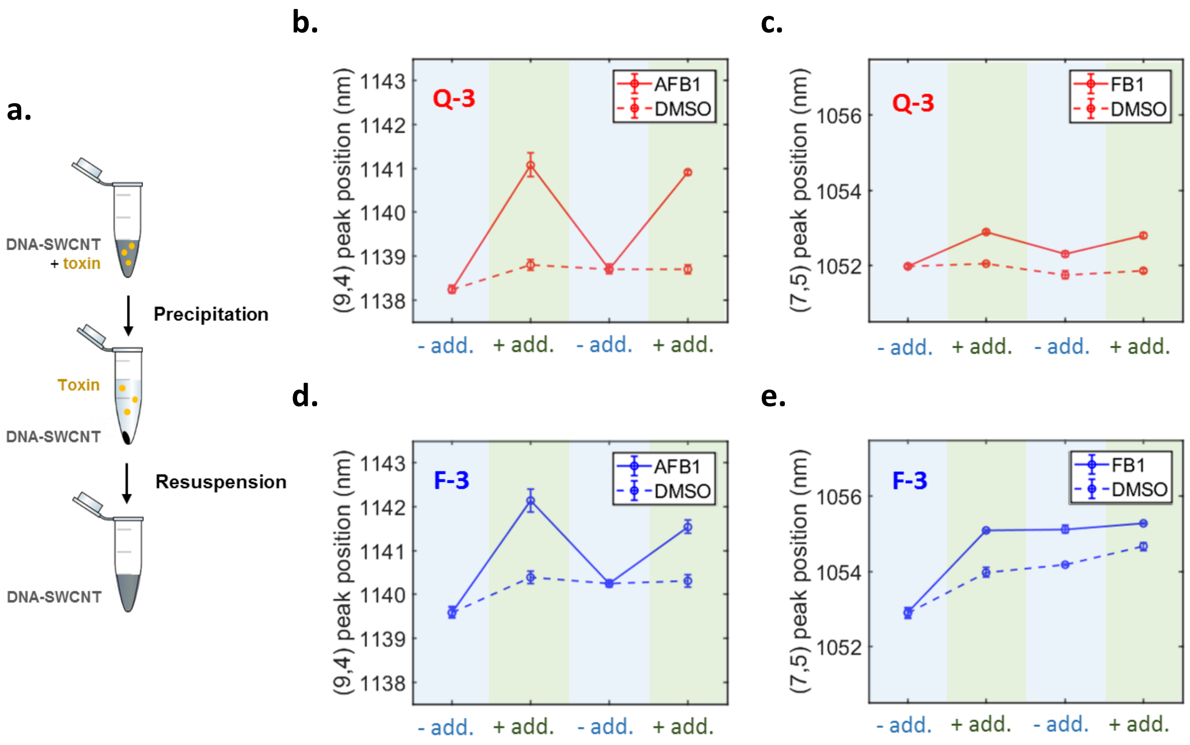


**Figure S30** - Reversibility of the Q-3 and F-3 sensors. (a) Schematic of the simple washing procedure. (b,c) Peak positions of the (9,4) and (7,5) chiralities for the Q-3 sensor, respectively, following the addition of toxin solution (10 µM). (d,e) Peak positions (9,4) and (7,5) chiralities for the F-3 sensor, respectively, following the addition of toxin solution (10 µM). All measurements were performed 200 min post-addition. The error bars represent 1 σ standard deviation (n = 3).

We verified that our DNA-SWCNT washing procedure was effectively removing the toxin both in the absence (**Table S6**) and presence of DNA-SWCNTs (**Figure S31**), using HPLC-MS and UV absorbance, respectively.

**Table S6** - Removal of toxin during ethanol precipitation was verified using HPLC-MS for AFB1 (red) and FB1 (blue) by calculating the relative concentration of toxin (in %) before and after ethanol precipitation.

| Name | AFB1 | FB1 |
|---|---|---|
| Before | 100 | 100 |
| After | 0.019 | 0.025 |

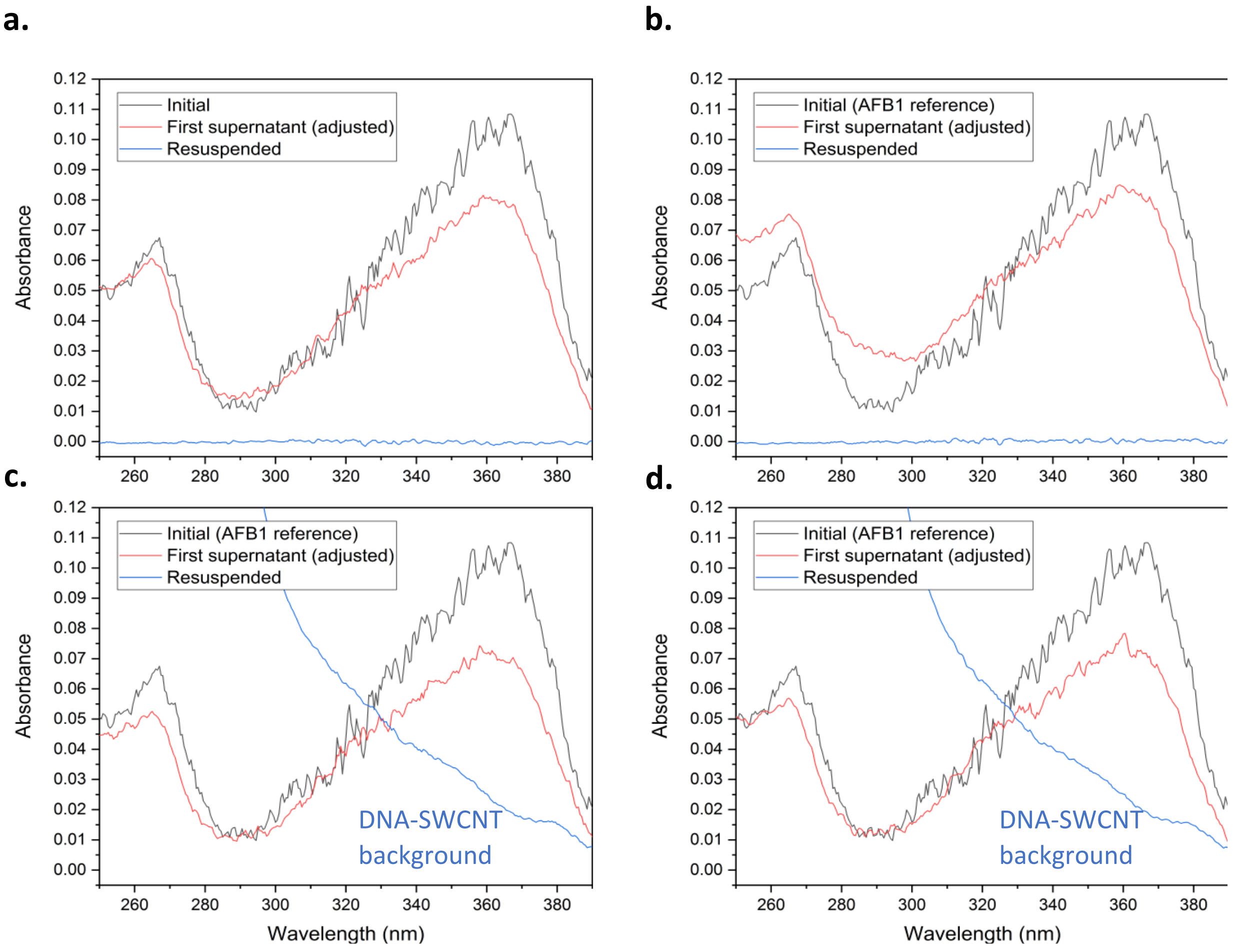


**Figure S31** - Removal of AFB1 during ethanol precipitation verified by UV-VIS. (a) Without nanotubes. (b) With 2% SC-SWCNTs. (c) With A-4 DNA-SWCNT. (d) With $\Delta(S5)_{16}$ DNA-SWCNT (16$^{th}$ mutant of S5). The first supernatant of the precipitation procedure (ethanol fraction, not containing SWCNTs) is shown here to contain most of the AFB1, hence proving the validity of this technique for the removal of toxins from DNA-SWCNT sensors.

We also tested the use of Amicon Ultra-0.5 3kDa centrifugal devices in order to remove the toxins (**Figure S32**). For that, we mixed 1 µL AFB1 with 99 µL 0.1 M NaCl or DNA-SWCNT in 0.1 M NaCl and 400 µL DI water and ran the mixture in the device (pre-rinsed) at 14,000 x g for 10 min. The absorbance of the first flow-through was assayed, which showed absence of any AFB1 absorbance peak. We washed a second time the device with 400 µL 50% ethanol (the toxin being more soluble in ethanol but also avoiding damage to the filter membrane) and the device was centrifuged at 14,000 x g for 20 min. Again, the flow-through was assayed, exhibiting traces of AFB1. Finally, the filter was washed a third time with 50% ethanol and centrifuged at 14,000 x g for 30 min. The flow-through of this fraction did not show any presence of AFB1. Finally, the filter was run three times with DI water (in order to replace the ethanol) and the filter fraction was assayed, demonstrating the absence of AFB1. While the filtering seems to result in removal of the toxins, it necessitates the use of organic solvent in the device which can damage the membrane and contaminate the sample with undesired compounds from the filter. In addition, the filtering method results in significant losses of DNA-SWCNTs which are limited when using the precipitation technique. Finally, the precipitation method offers a greater flexibility in the volume and concentration of the DNA-SWCNT used (a too large concentration of SWCNT can block the filter membrane).

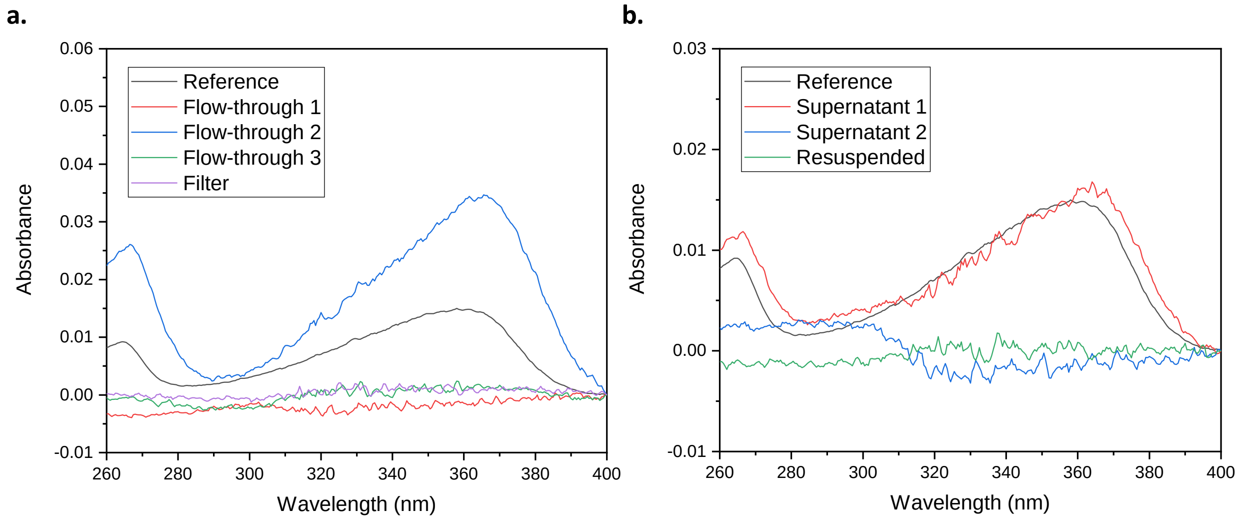


**Figure S32** - Comparison of AFB1 removal with filtering and precipitation. (a) Absorbance spectra of the samples processed by filtering with an Amicon Ultra-0.5 3kDa filter. The reference indicates the initial sample containing AFB1. DI water was added to the sample for the first centrifugation, 50% ethanol was added for the second and third centrifugations. The filter fraction was washed several times with DI water to remove traces of ethanol and resuspended in 0.1M NaCl. (b) Absorbance spectra of the samples processed by precipitation. The reference indicates the initial sample containing AFB1. The first precipitation was performed with 100% ethanol and the second precipitation with 70% ethanol. The samples were resuspended in 0.1 M NaCl.

As the FB1 sensor could not be reversed, we modified the washing protocol to insert an additional denaturing step (see **Materials and methods**

) in order to disrupt interactions between the FB1 and the DNA as illustrated in **Figure S33**.

Heat-denatured washing protocol

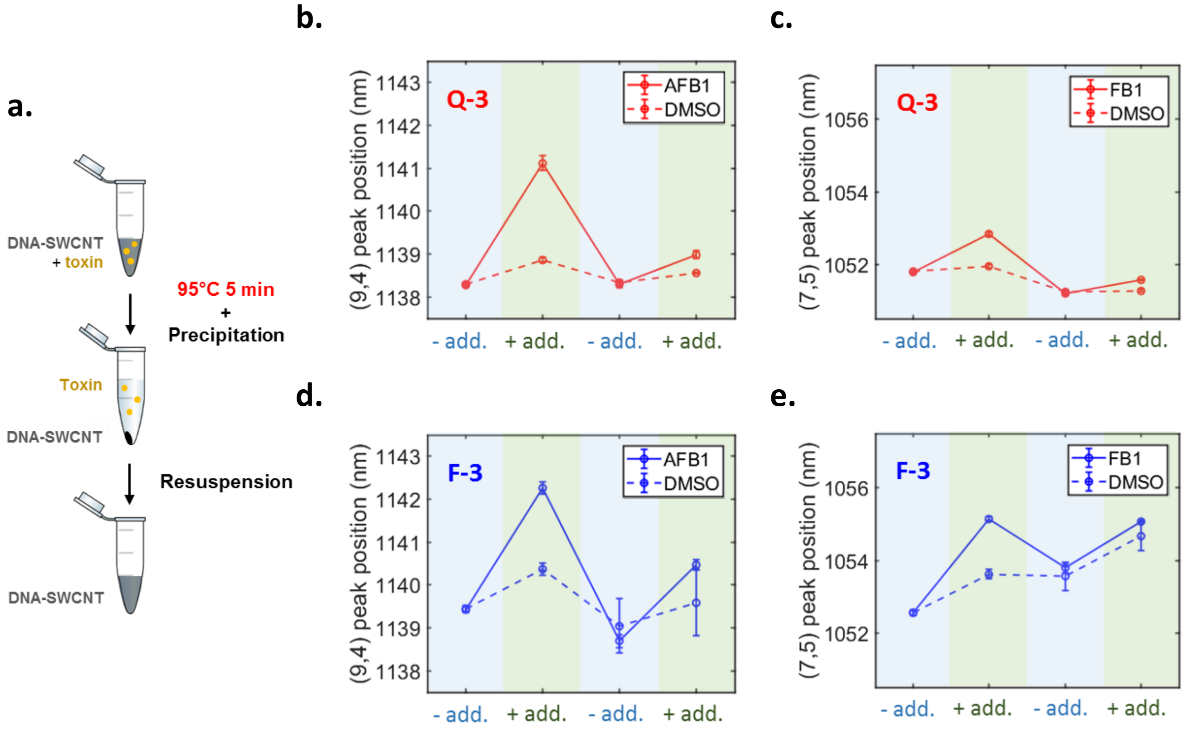


**Figure S33** - Reversibility of the Q-3 and F-3 sensors when using an additional denaturation step. (a) Schematic of the modified washing procedure with the additional denaturation step to disrupt the interaction between the toxin and the DNA-SWCNTs. (b,c) Peak positions of the (9,4) and (7,5) chiralities for the Q-3 sensor, respectively, following the addition of toxin solution (10 µM). (d,e) Peak positions (9,4) and (7,5) chiralities for the F-3 sensor, respectively, following the addition of toxin solution (10 µM). All measurements were performed 200 min post-addition. The error bars represent 1 σ standard deviation (n = 3).

#### Supplement 16: Free DNA loss during precipitation

The amount of DNA lost during the ethanol precipitation of DNA-SWCNT was estimated using the fluorescence of a Sybr Gold DNA dye assay. We first calibrated our assay to relate the fluorescence signal of the dye with DNA concentration (**Figure S34**). The concentration of DNA was determined for the supernatants from the first and second ethanol precipitations. Ethanol from the supernatants was removed using Amicon Ultra 0.5 3kDa centrifugation devices and the concentrated DNA was then resuspended in DI water. A significant amount of DNA was lost during the precipitation process, especially during the second precipitation step (the 70% ethanol fraction).

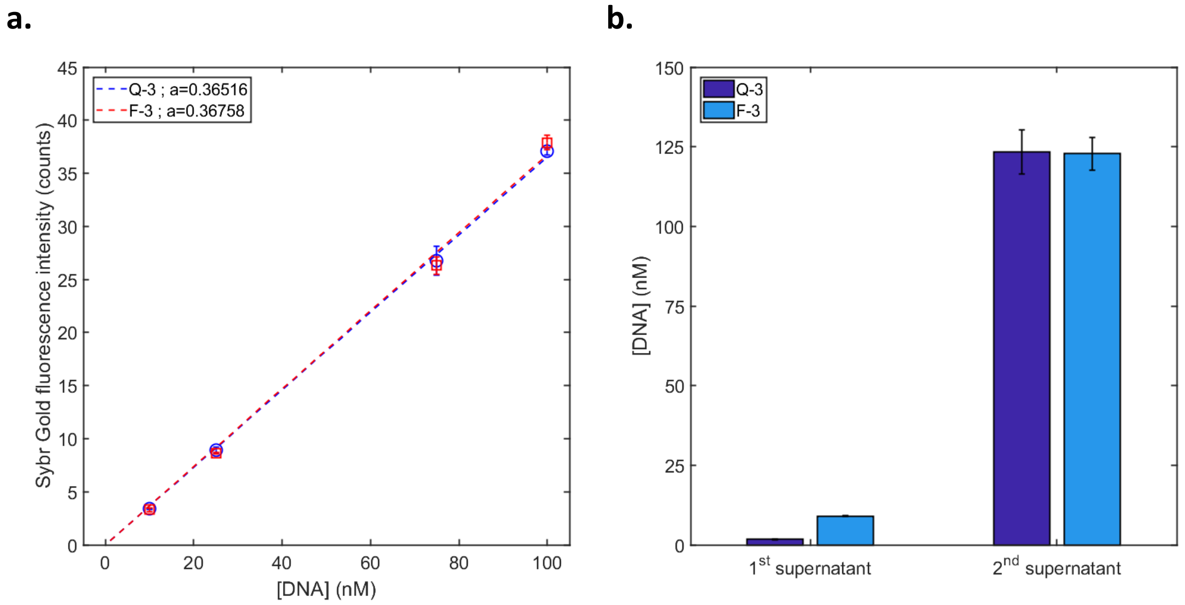


**Figure S34** - Determination of the amount of DNA lost during DNA-SWCNT precipitation. (a) The calibration curve of the Q-3 and F-3 DNA to the fluorescence of the Sybr Gold dye was used to determine the free DNA concentration. (b) Determined concentration of DNA in supernatants from the DNA-SWCNT precipitation protocol. Error bars represent 1 σ standard deviation (n = 3).

#### Supplement 17: Effect of free DNA on the properties of the sensors

We investigated the effect of removing the free DNA on the peak position (**Figure S35**) and the shifting response (**Figure S36**) of the Q-3 and F-3 sensors. To remove free DNA, we used Amicon Ultra 0.5 100 kDa centrifugation devices, washing the nanotube solution six times with DI water to remove all traces of free DNA before resuspension. We noted a chirality dependent red-shift in the peak position following the removal of free DNA, with greater shifting observed for the F-3 sensor compared to the Q-3 sensor. The removal of free DNA also resulted in a reduced response for both sensors towards AFB1 and FB1. Interestingly, the response of the Q-3 sensor towards FB1 was less affected by the removal of free DNA than the F-3 sensor.

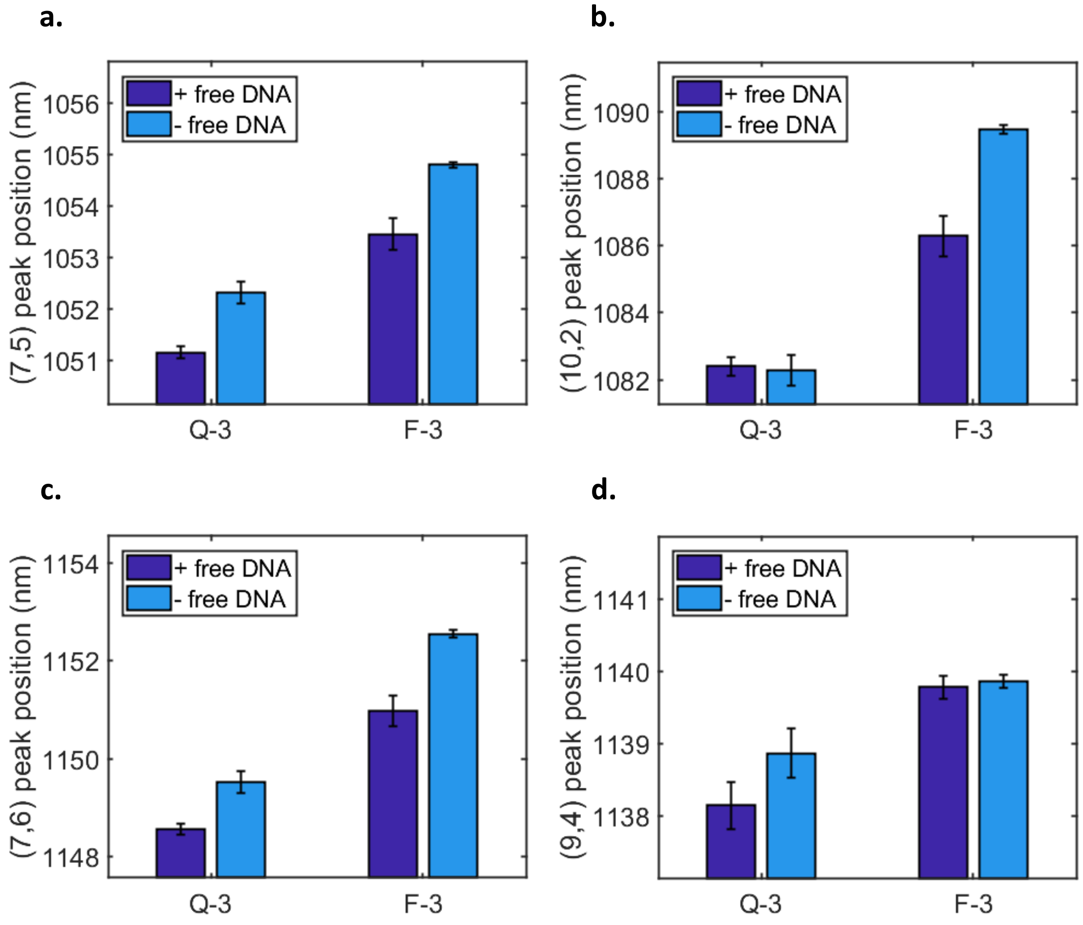


**Figure S35** - Peak position of the Q-3 and F-3 sensors with and without free DNA for the (a) (7,5), (b) (10,2), (c) (7,6) and (d) (9,4) chiralities. The error bars represent 1 σ standard deviation (n = 3).

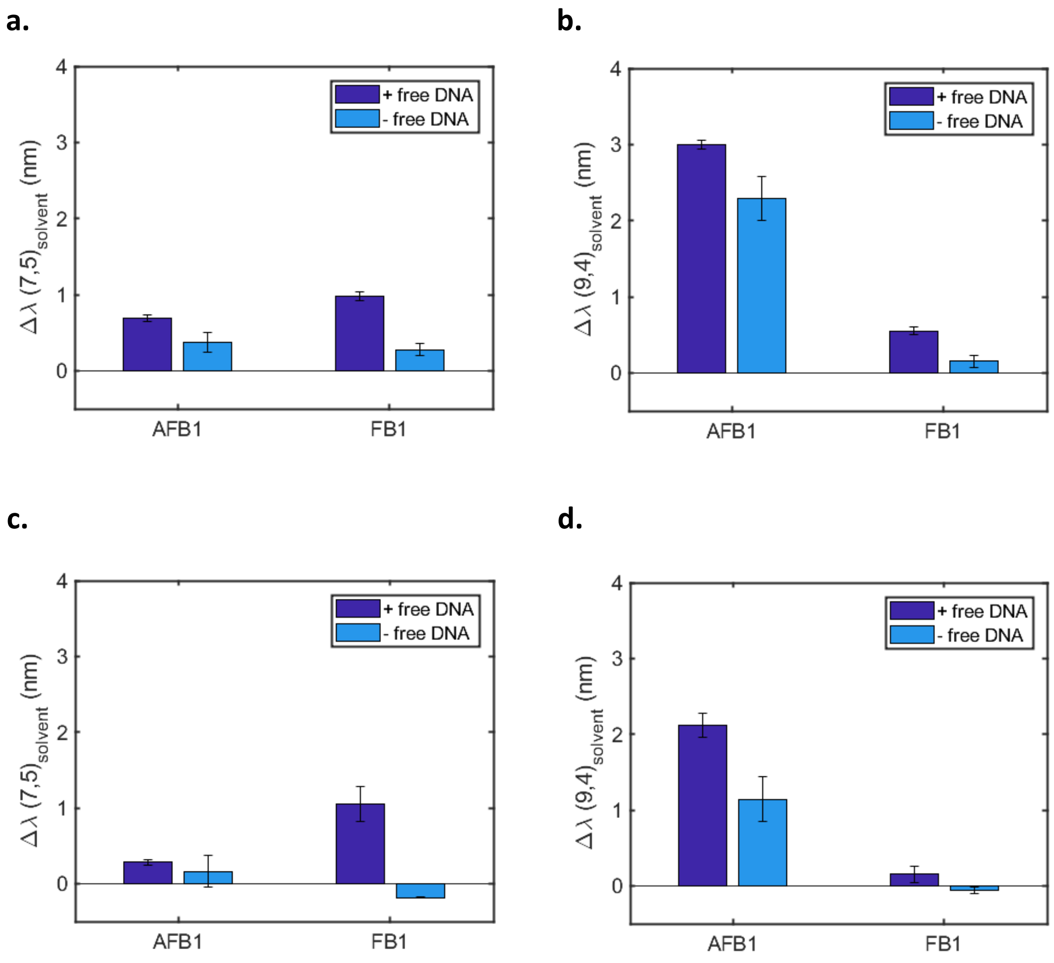


**Figure S36 -** Wavelength shifting response of the (a,b) Q-3 and (c,d) F-3 sensors with and without free DNA towards AFB1 (10 μM) and FB1 (10 μM) for the (7,5) and (9,4) chiralities. All measurements were performed 200 min post-addition. The error bars represent 1 σ standard deviation (n = 3).

#### Supplement 18: Absence of covalent interaction

The 8,9-epoxide exo isomer of AFB1 (AFBO) can form covalent adducts with guanine nucleobases. The AFBO is typically formed by the P450 cytochrome in the human metabolism[16], but additional studies have also confirmed the production of AFBO through the photo-oxidation of AFB1[17]. We verified the absence of AFBO in our samples by HPLC-MS (**Figure S37**). While we observed a clear peak for AFB1 using HPLC-MS, no peaks corresponding to the m/z of aflatoxin B1 exo-8,9-epoxide or aflatoxin B1 exo-8,9-dihydrodiol were detected, confirming their absence from our solutions.

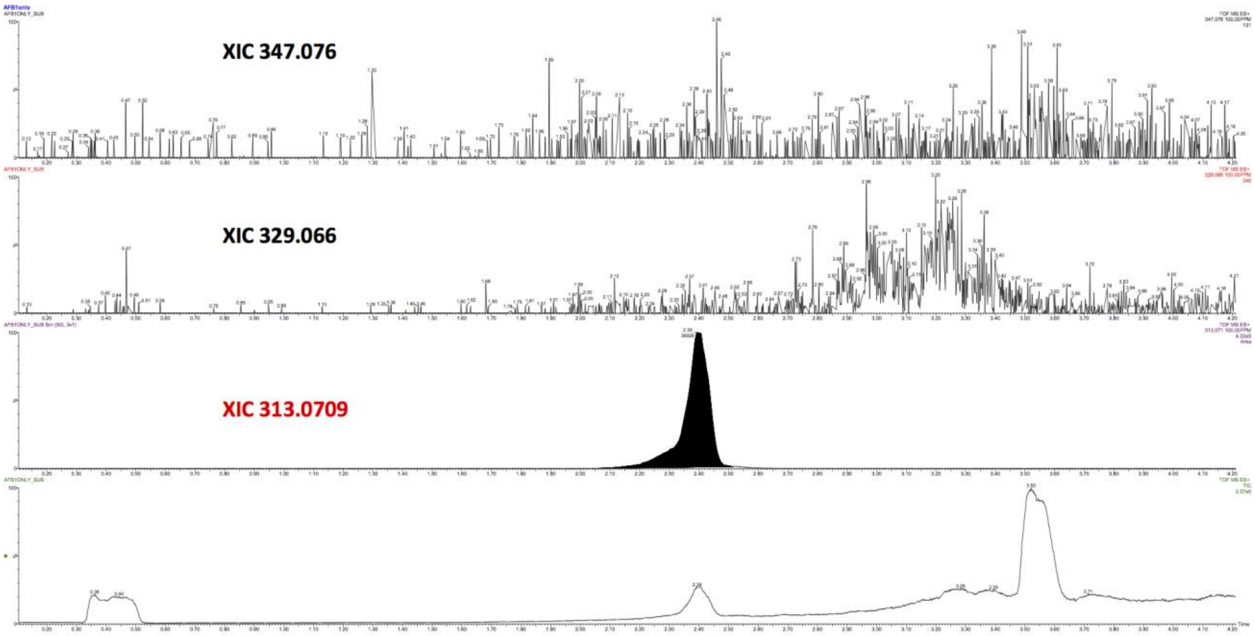


**Figure S37 -** HPLC-MS chromatograms for compounds of m/z equal to (from top to bottom) 347.076, 329.066, 313.0709, and all ions. The m/z of 347.076 corresponds to aflatoxin B1 exo-8,9-dihydrodiol ($C_{17}H_{14}O_8$), the m/z of 329.066 corresponds to aflatoxin B1 exo-8,9-epoxide ($C_{17}H_{12}O_7$), and the m/z of 313.0709 corresponds to aflatoxin B1.

In addition, we tested our sensors in presence of aflatoxin B2 (AFB2) in order to demonstrate the absence of covalent adducts between the toxin and the DNA (**Figure S38**). AFB1 and AFB2 are structurally similar, however while AFB1 can be metabolized into AFBO and form a covalent adduct with DNA bases, AFB2 cannot. We noted that the Q-3 and F-3 sensors exhibited similar wavelength shifting for the (9,4) peak towards both AFB1 and AFB2. We therefore concluded that the observed response does not result from the formation of covalent adducts between the AFB1 and the DNA wrapping. Moreover, this generalizes the use of our sensors for the detection of both aflatoxins.

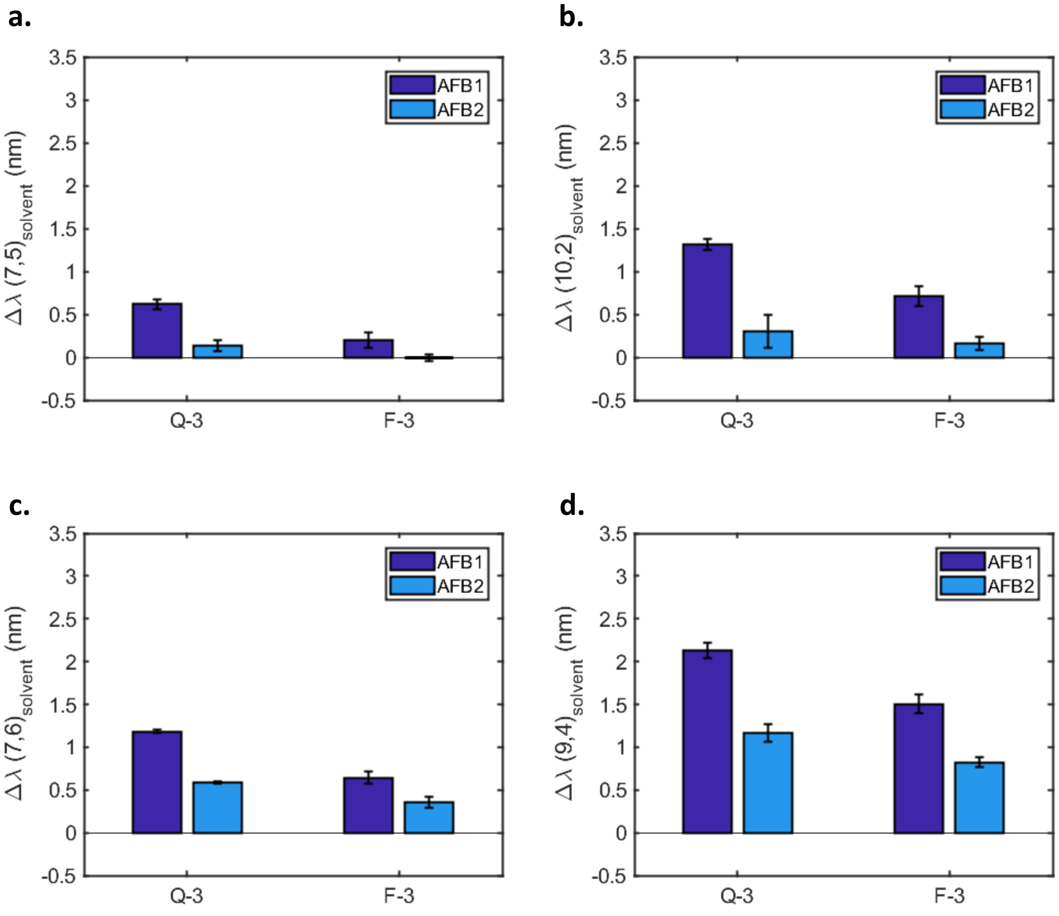


**Figure S38 -** Response of the Q-3 and F-3 sensors to AFB1 (10 µM) and AFB2 (10 µM) for the (a) (7,5), (b) (10,2), (c) (7,6), and (d) (9,4) chiralities. All measurements were performed 200 min post-addition. The error bars represent 1 σ standard deviation (n = 3).

#### Supplement 19: Interaction between the toxins and the SWCNT surface

We tested the interaction of the AFB1 and FB1 toxins with exposed SWCNT surface using sodium cholate (SC)-wrapped SWCNTs. Nanotube suspensions with different surface coverages were prepared using different concentrations of SC, between 0.5 mM and 46 mM, as previously described[18] ; lower SC concentrations resulted in lower the surface coverage. Upon addition of AFB1 (**Figure S39**), we did not observe any change in the wavelength position for the high coverage sample (46 mM SC) but observed a strong red-shift of the (9,4) peak for the sample with the lowest coverage (0.5 mM SC). This indicated that AFB1 was able to interact directly with the SWCNT surface. This red-shifting was also linked to a decrease in fluorescence intensity, indicating that the SC was being removed from the surface of the SWCNT (**Figure S40**). In the case of FB1 (**Figure S41**), we did not observe any shifting of the (7,5) chirality, even for the low concentration SC-SWCNTs (0.5 mM) confirming the absence of any interaction between the FB1 and the SWCNT surface. The absence of any interaction was confirmed by the steady fluorescence intensity recorded even post-addition of FB1 (**Figure S42**).

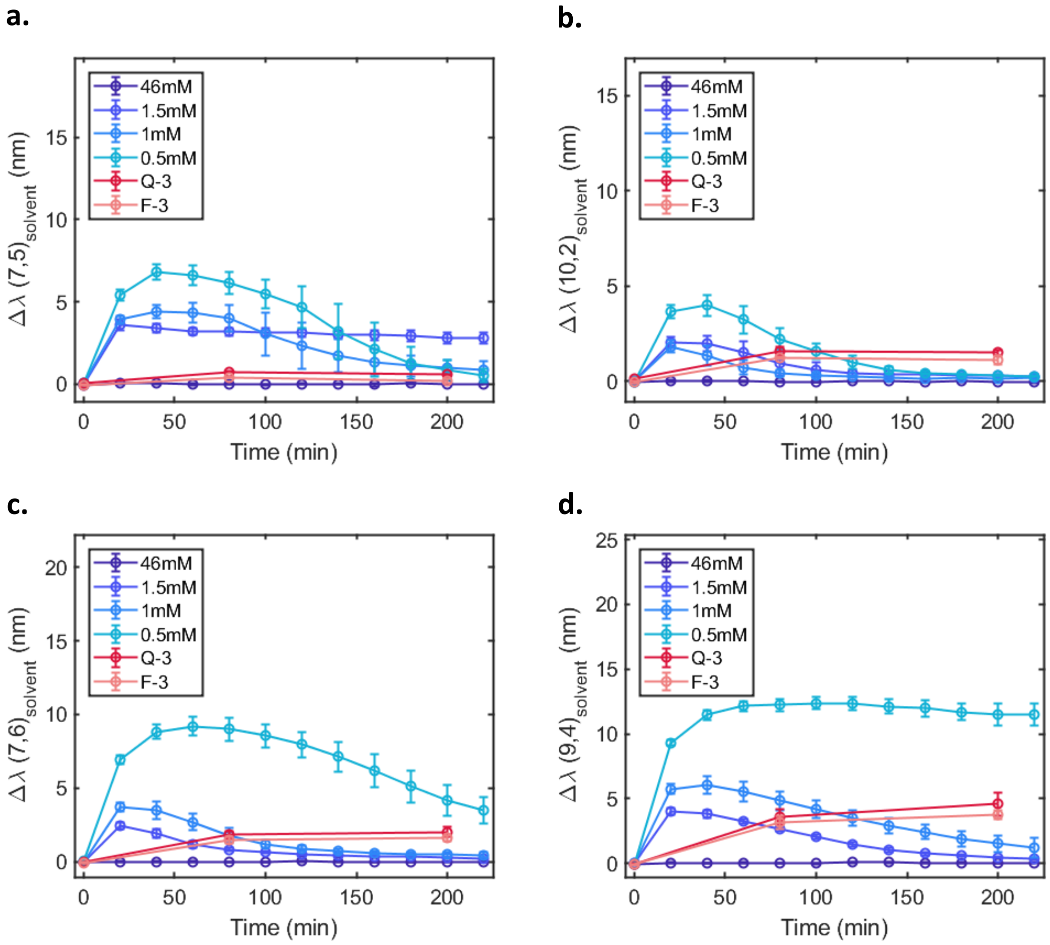


**Figure S39 -** Comparison of the wavelength shifting response of surfactant-wrapped SWCNTs (blue) and DNA-SWCNTs (red) following the addition of AFB1 (10 μM) for the (a) (7,5), (b) (10,2), (c) (7,6), and (d) (9,4) chiralities as a function of time. The error bars represent 1 σ standard deviation (n = 3).

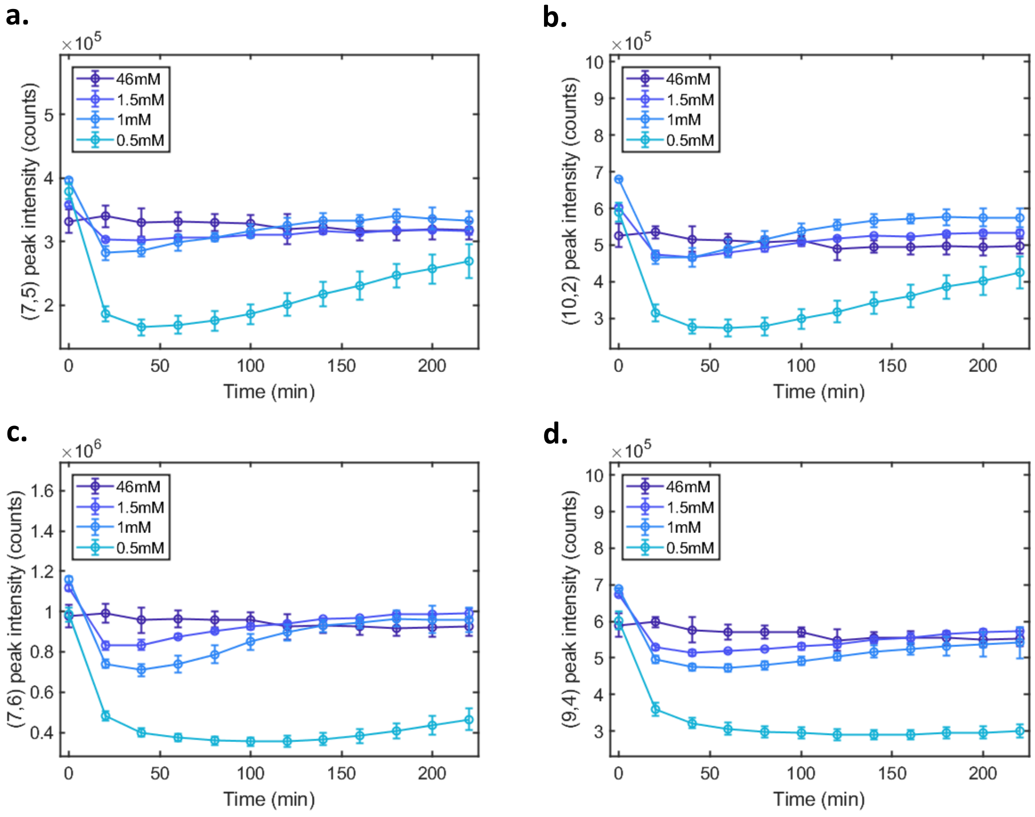


**Figure S40 -** Comparison of the fluorescence intensity response of surfactant-wrapped SWCNTs (blue) and DNA-SWCNTs (red) following the addition of AFB1 (10 μM) for the (a) (7,5), (b) (10,2), (c) (7,6), and (d) (9,4) chiralities as a function of time. The error bars represent 1 σ standard deviation (n = 3).

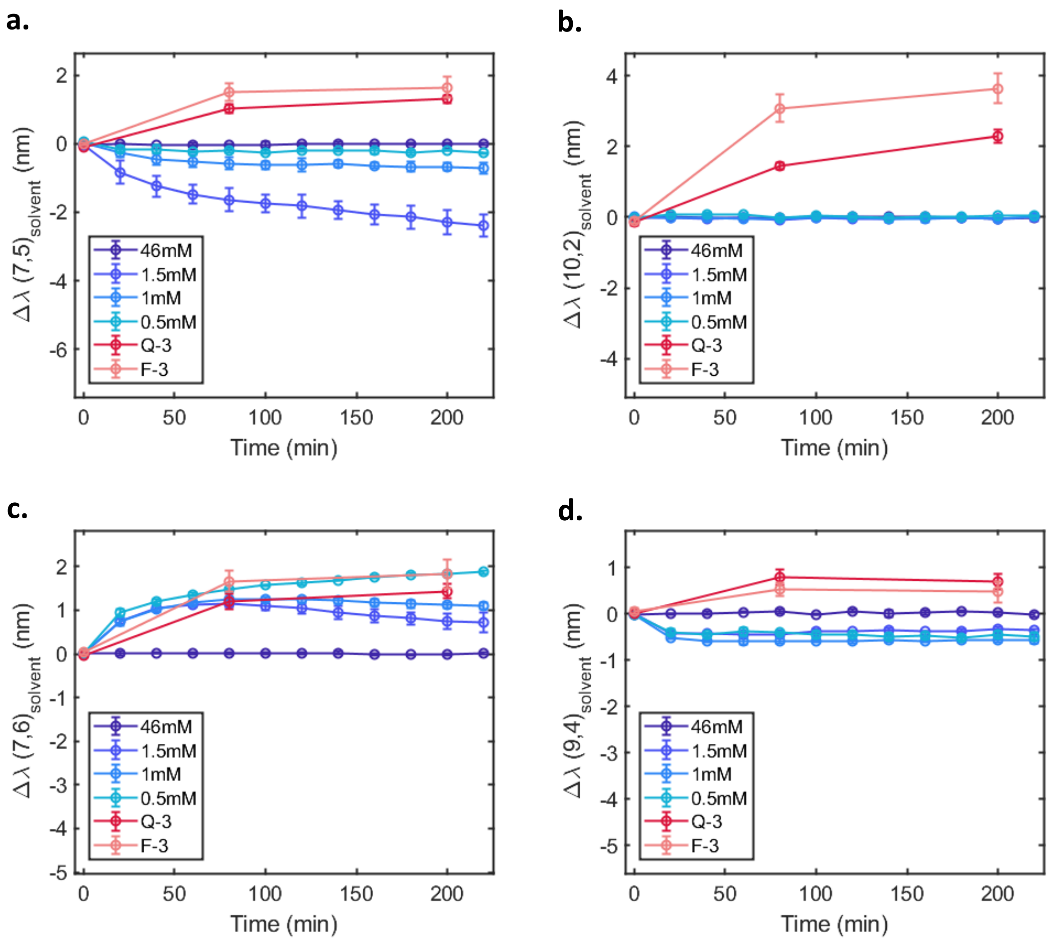


**Figure S41 -** Comparison of the wavelength shifting response of surfactant-wrapped SWCNTs (blue) and DNA-SWCNTs (red) following the addition of FB1 (10 μM) for the (a) (7,5), (b) (10,2), (c) (7,6), and (d) (9,4) chiralities as a function of time. The error bars represent 1 σ standard deviation (n = 3).

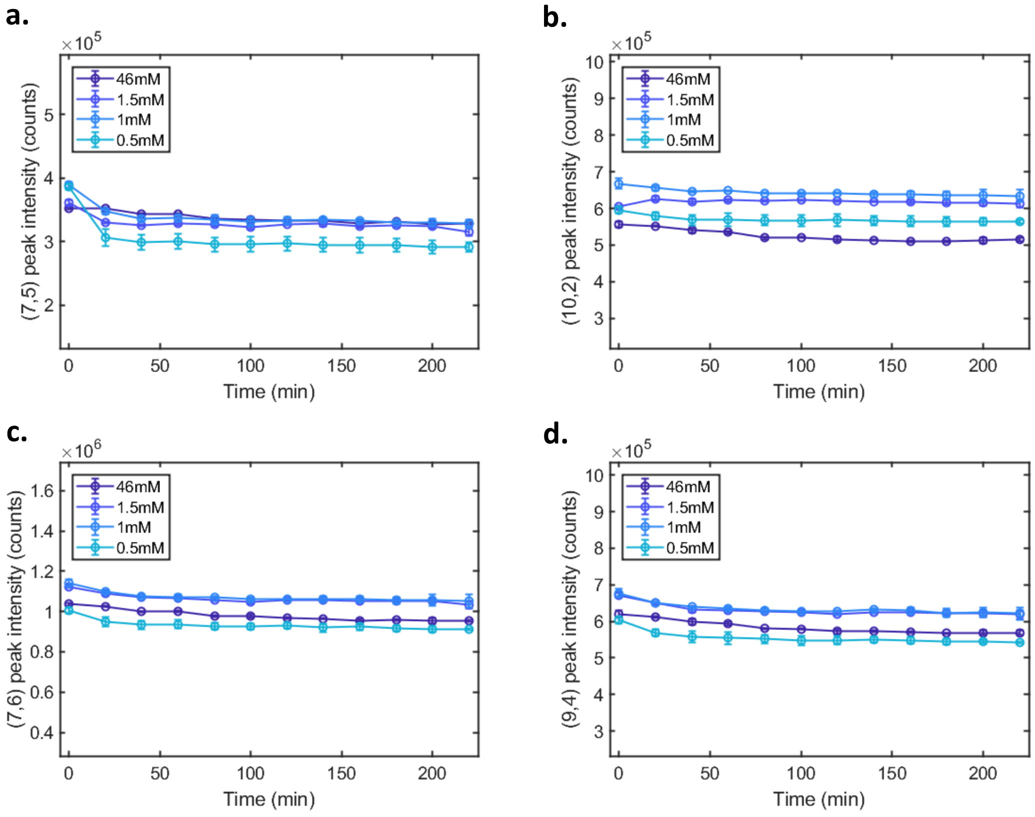


**Figure S42 -** Comparison of the fluorescence intensity response of surfactant-wrapped SWCNTs (blue) and DNA-SWCNTs (red) following the addition of FB1 (10 μM) for the (a) (7,5), (b) (10,2), (c) (7,6), and (d) (9,4) chiralities as a function of time. The error bars represent 1 σ standard deviation (n = 3).

#### Supplement 20: Removal of AFB1 from the SWCNT surface

Titrating the toxin-SWCNT mixture with low concentrations of SDBS surfactant was used to revert the red-shifting caused by the addition of AFB1. This further supported that the AFB1 interacts with the SWCNT surface.

**a.**

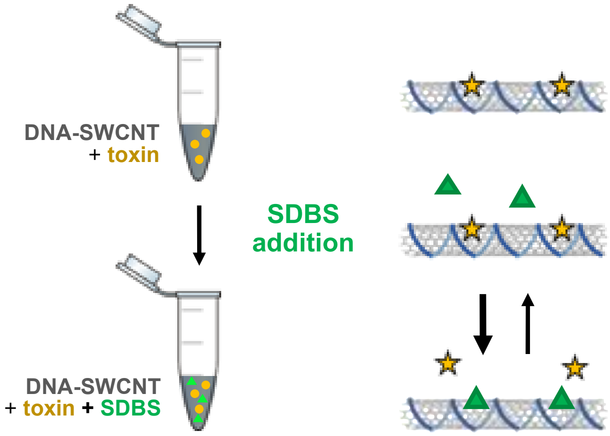


**b.**

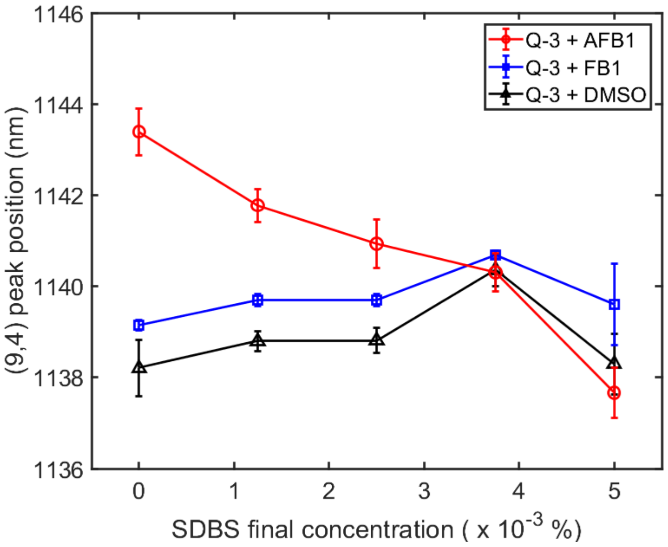


**Figure S43 -** Removal of AFB1 from DNA-SWCNT sensors. (a) Schematic of the protocol used to remove AFB1 from the surface of the DNA-SWCNTs using SDBS. The SDBS molecules replace the AFB1 molecules on the SWCNT surface, shifting the emission back to its initial position. (b) The position of the (9,4) peak for the Q-3 sensor incubated with AFB1 (10 µM, red), FB1 (10 µM, blue), or DMSO (black) in presence of varying concentrations of SDBS: 0, 1, 2.5, 3.75, 5 x $10^{-3}$ %. The error bars represent 1 σ standard deviation (n = 3).

#### Supplement 21: DNA concentration removed after analyte interaction

The concentration of free DNA before and after incubation with AFB1, FB1, and DMSO was estimated for the Q-3 and F-3 sensors (**Figure S44**). The sensors were washed using 100kDa Amicon centrifugation devices and the concentration of the free DNA present in the flow-through of the filtering devices was estimated using the Sybr Gold assay described in **Figure S34**. Interestingly, the free DNA concentrations were 3-5 times higher for the F-3 sensor compared to the Q-3 sensor. For both sensors, incubation with FB1 resulted in an increase in the concentration of free DNA compared to the control (solvent). On the other hand, incubation with AFB1 resulted in either no difference or decreased concentrations of free DNA versus the solvent control. Moreover, in both cases, the free DNA concentration was lower following incubation with DMSO compared to the samples pre-addition.

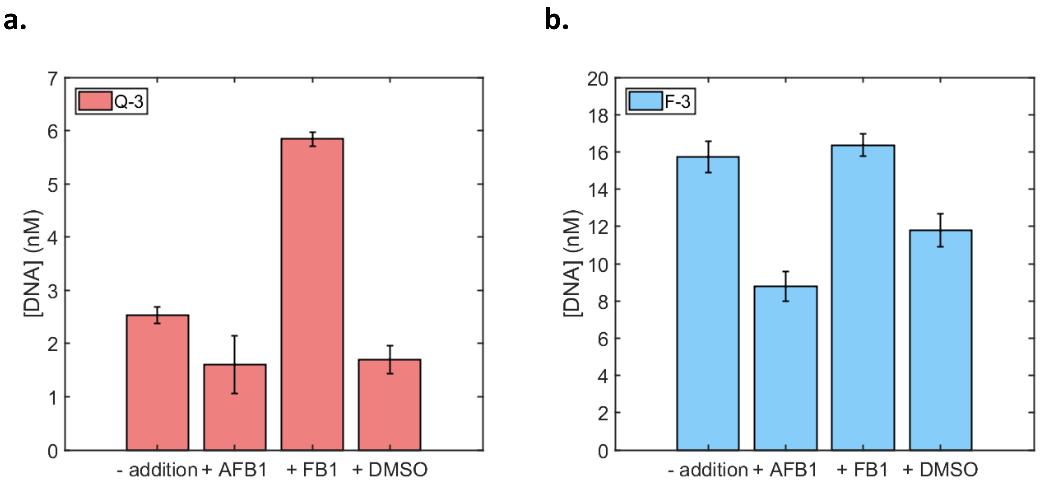


**Figure S44** - Concentration of free DNA for (a) the Q-3 and (b) the F-3 sensors before and after addition of AFB1 (10 μM), FB1 (10 μM), and DMSO. Error bars represent 1 σ standard deviation (n = 3).

#### Supplement 22: Binding affinity of Q-3 and F-3 sensors

In order to determine the binding affinity of the Q-3 and F-3 sequences to SWCNT, we performed surfactant replacement experiments using SDBS (final concentration of 0.01% or 0.05%). The surfactant replacement occurs faster for the F-3 sensor compared to the Q-3 sensor, indicating that the F-3 DNA has a lower binding affinity to SWCNT surface.

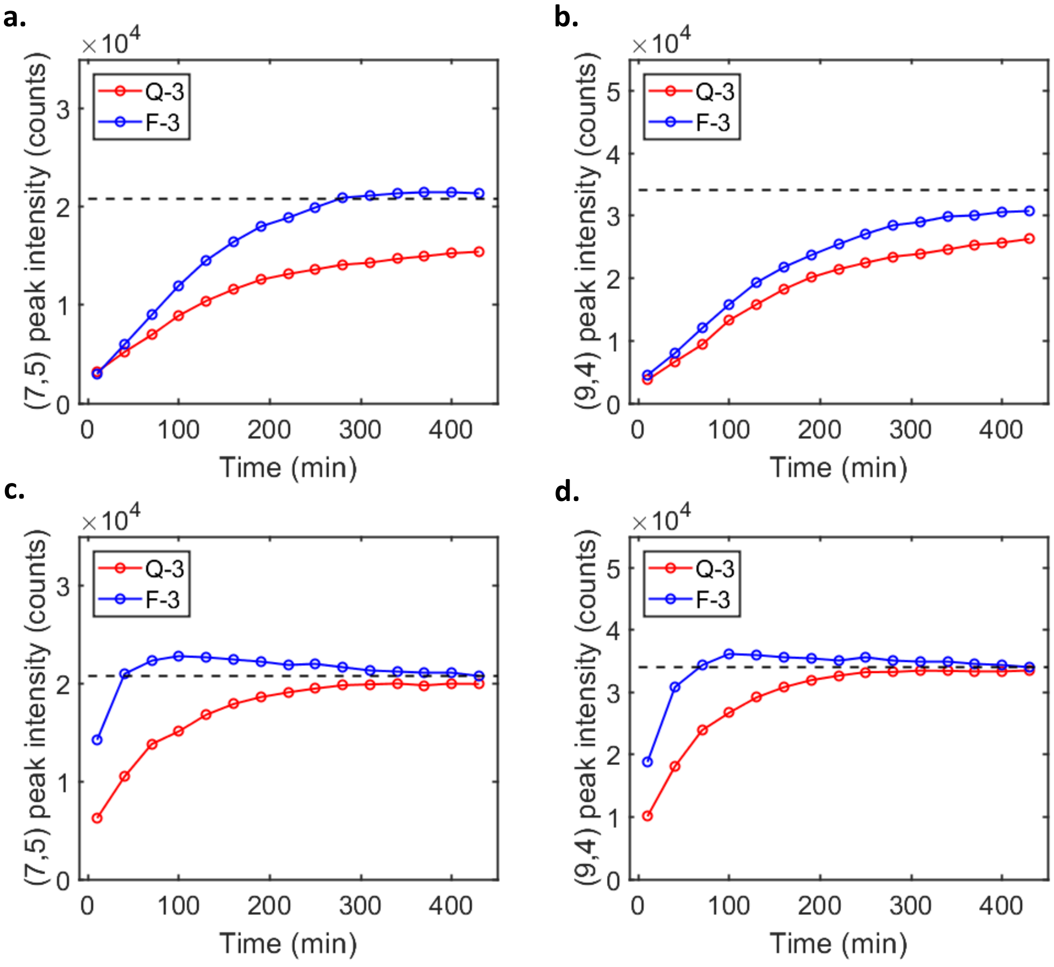


**Figure S45 -** Replacement kinetics of the DNA for the (7,5) and (9,4) chiralities following the addition of SDBS for the Q-3 (red) and F-3 (blue) sensors. SDBS was added to a final concentration of (a,b) 0.01% and (c,d) 0.05% and the fluorescence peak intensities were monitored over time. The intensity corresponds to the maximum intensity at the final position of the (7,5) and (9,4) post-SDBS addition.

#### Supplement 23: Flow chamber

Since the reversible AFB1 sensors showed minimal responsivity when immobilized (**Figure S29**), we constructed a liquid-phased flow chamber consisting of a dialysis membrane separating the sensors and the analyte and monitored the response of the F-3 sensor upon addition and removal of AFB1 (**Figure S46**).

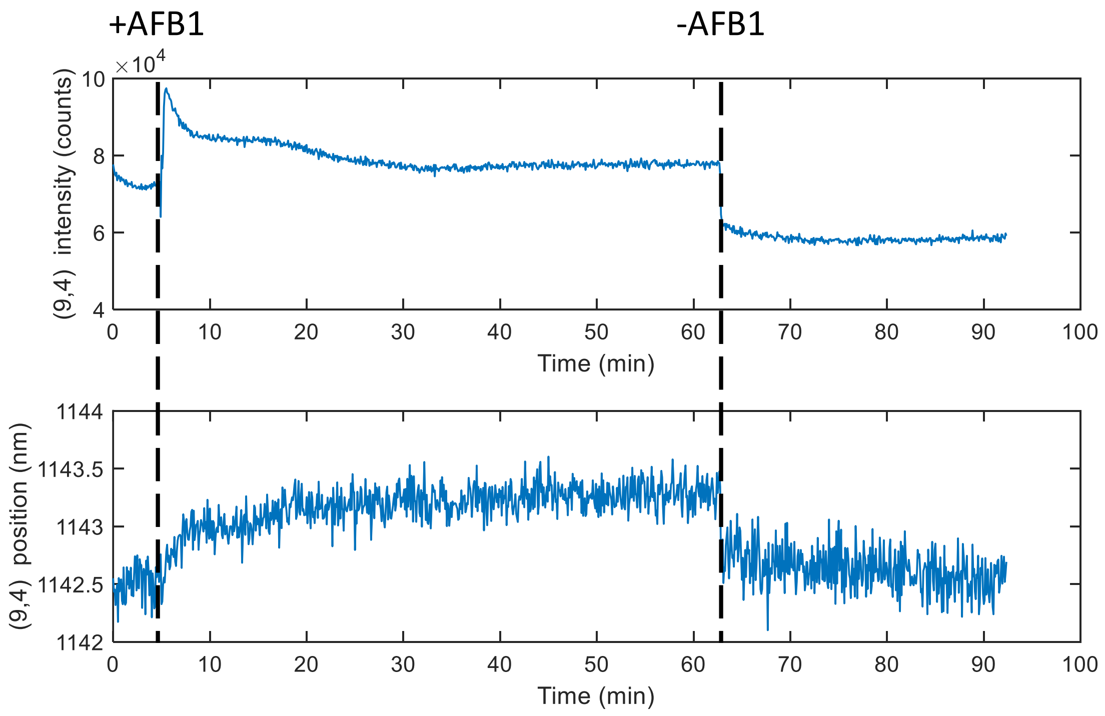


**Figure S46 -** Reversibility of the AFB1 sensing in a flow chamber. (top) Peak intensity and (bottom) peak position of the F-3 sensor in presence of 0.1M NaCl (between 0 and 5 min), 10 µM AFB1 (between 5 min and 63 min) and washed with 0.1 M NaCl again (after 63 min). The sensors experience a red-shifting of the emission upon addition of AFB1 which is reversed upon washing with buffer. This therefore proves that the sensors could be used in a flow chamber for practical use.

#### Supplement 24: High-throughput DNA removal

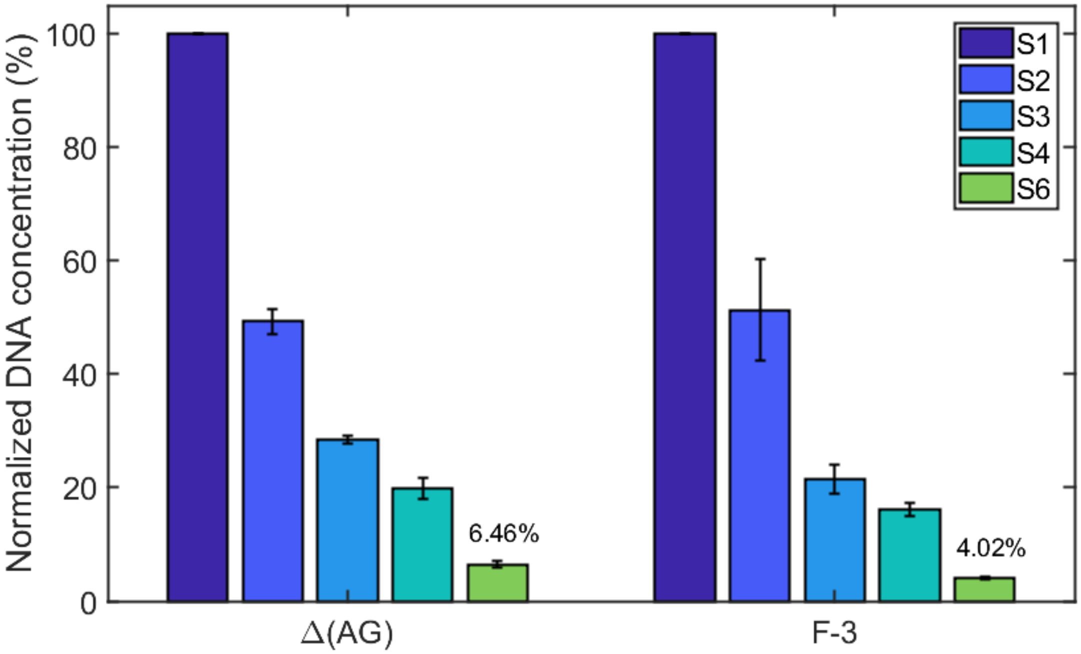


**Figure S47 -** Removal of free DNA with a high-throughput procedure. The DNA-SWCNT suspensions are precipitated with salts and ethanol similarly to the toxin removal procedure. Then, the pellet is washed several times with 70% ethanol in order to wash away the free DNA from the samples. The normalized concentrations of free DNA in the 70% ethanol supernatants of the Δ(AG) and F-3 DNA-SWCNT sensors are shown here for the first (S1) to the sixth (S6) supernatant fractions. The method effectively removes more than 94% of the free DNA in 6 washes, allowing for a more scalable approach for free DNA removal and more compatible with directed evolution. Error bars represent 1 σ (n = 3).

#### Supplement 25: Effect of ionic strength on AFB1 response

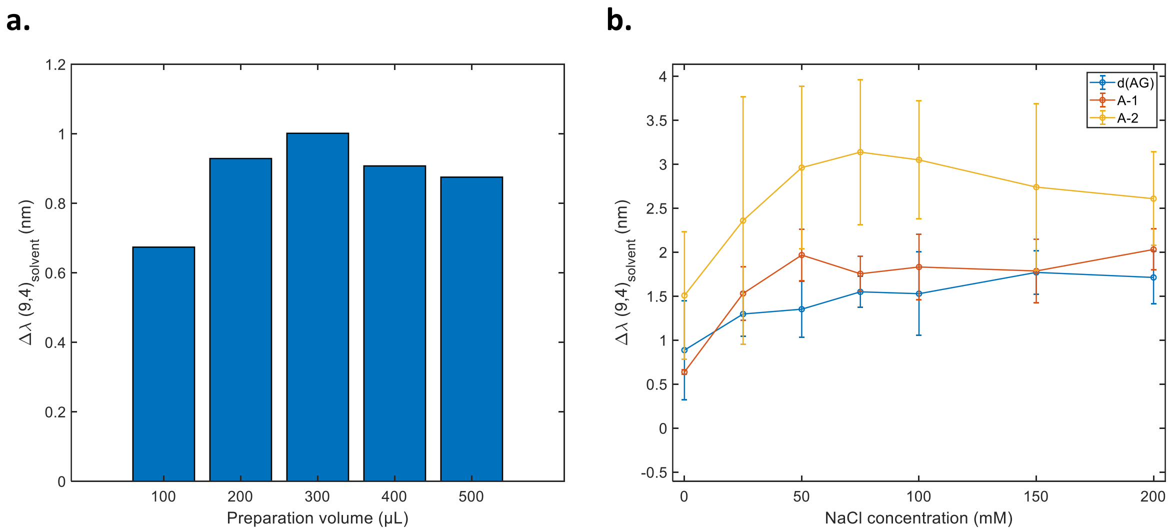


**Figure S48 -** Influence of preparation volume and ionic strength on AFB1 response. Shifting response of the (9,4) chirality in presence of AFB1 as a function of (a) the sample preparation volume for the Δ(AG) sensor and (b) the NaCl concentration for the Δ(AG), A-1 and A-2 sensors. The measurements were performed with an AFB1 concentration of 10 µM and 200 min after addition. The error bars represent 1 σ (n=3). The samples in (a) were resuspended in 0.1M NaCl and the samples in (b) were prepared in a 300 µL volume. The AFB1 batches prepared under non-inert conditions in DMSO. These controls show first that, unless prepared in low volume (100 µL), the behavior of the sensors does not depend on the preparation volume. The controls also show that the response is optimized at a salt concentration of 0.1 M.

#### Supplement 26: Reaction kinetics of the S5 sensor to AFB1 and FB1

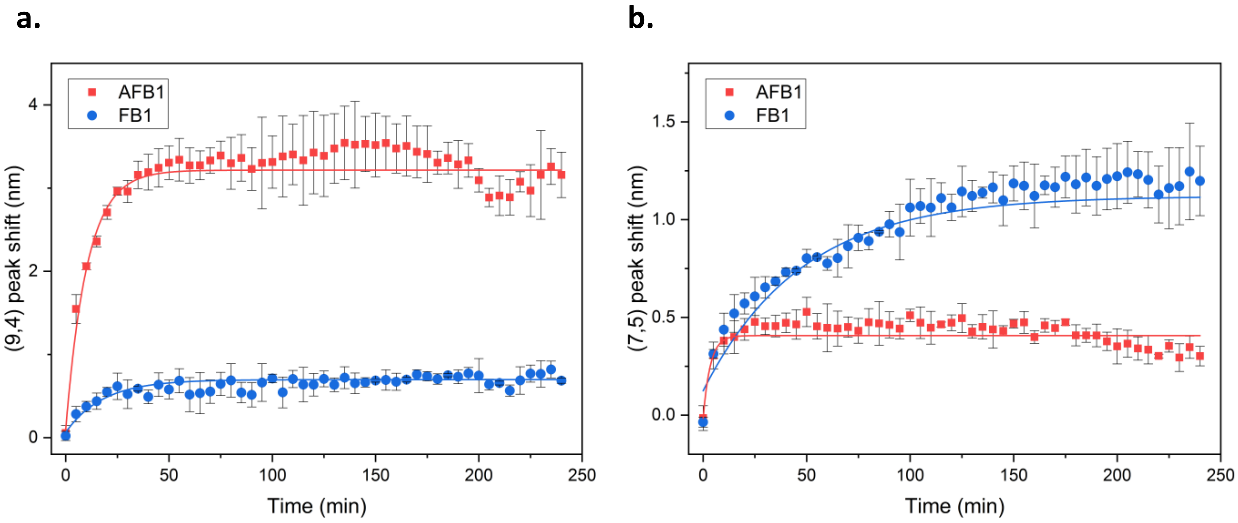


**Figure S49** - Response kinetics for the (a) (9,4) and (b) (7,5) chiralities of the S5 sensor towards AFB1 (5 μM, red) and FB1 (5 μM, blue) for the. We noted that for the S5 sensor, the AFB1 response reached its maximum after less than 50 min while the FB1 response took almost 200 min to plateau. All fits are single exponential fits. Error bars represent 1 σ standard deviation (n = 3).

#### Supplement 27: Comparison of the responses of reactive and non-reactive DNA-SWCNTs towards mycotoxins

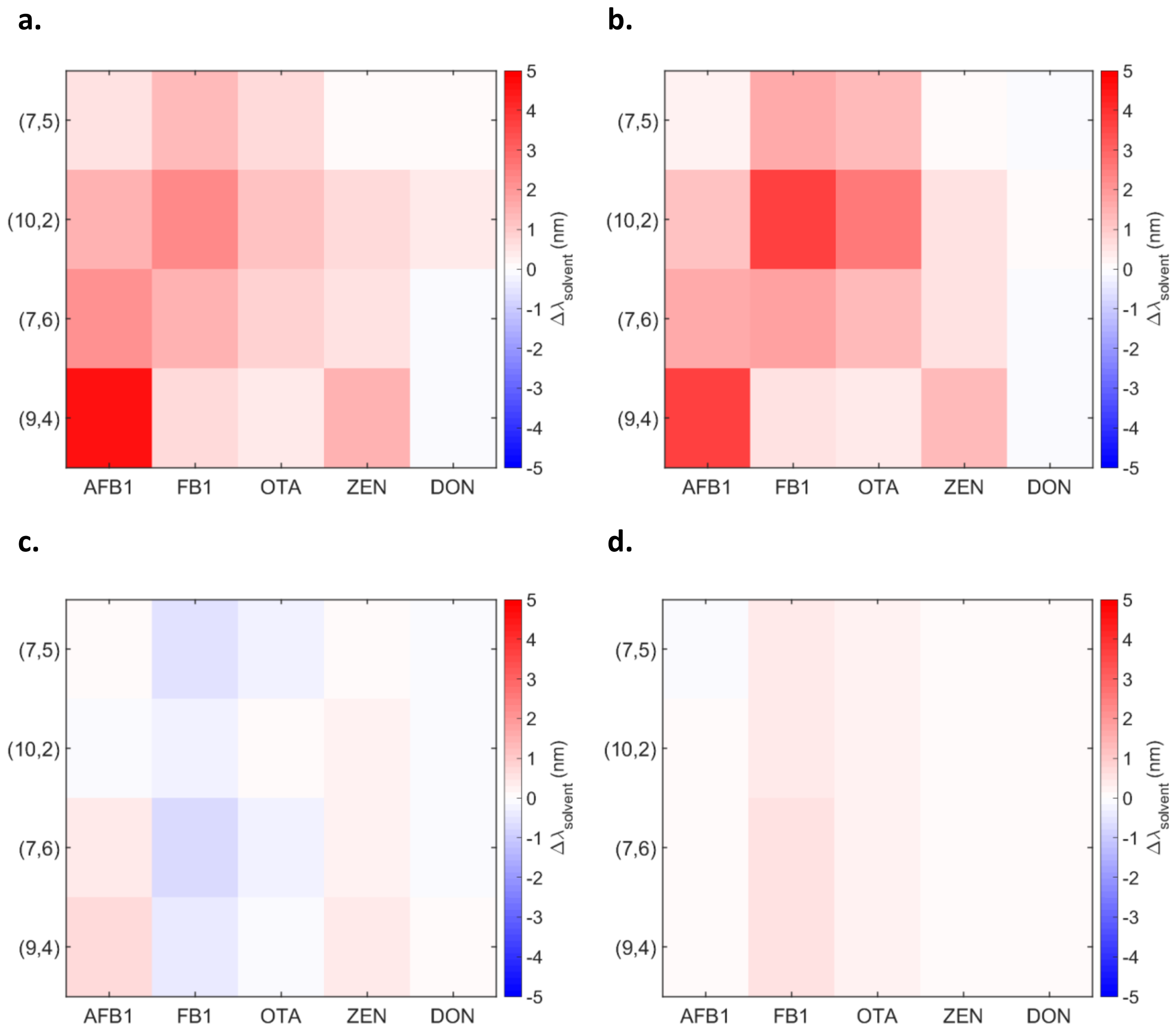


**Figure S50** - Heatmaps of the wavelength shifting response of the (a) Q-3, (b) F-3, (c) (AG)15, and (d) Δ(AT) sensors to the addition of 10 μM AFB1, FB1, OTA, ZEN or DON. All measurements were recorded 200 min post-addition.

#### References


1. Zubkovs, V., Schuergers, N., Lambert, B., Ahunbay, E. & Boghossian, A. A. Mediatorless, Reversible Optical Nanosensor Enabled through Enzymatic Pocket Doping. *Small* **13**, 1–10 (2017).
2. Lambert, B., Gillen, A. J., Schuergers, N., Wu, S.-J. & Boghossian, A. A. Directed evolution of the optoelectronic properties of synthetic nanomaterials. *ChemComm* **55**, 3239–3242 (2019).
3. Zheng, M. *et al.* DNA-assisted dispersion and separation of carbon nanotubes. *Nat. Mater.* **2**, 338–342 (2003).
4. Zhang, J. *et al.* Single molecule detection of nitric oxide enabled by d(AT)15 DNA adsorbed to near infrared fluorescent single-walled carbon nanotubes. *J. Am. Chem. Soc.* **133**, 567–581 (2011).
5. Kruss, S. *et al.* Neurotransmitter detection using corona phase molecular recognition on fluorescent single-walled carbon nanotube sensors. *J. Am. Chem. Soc.* **136**, 713–724 (2014).
6. Beyene, A. G. *et al.* Ultralarge Modulation of Single Wall Carbon Nanotube Fluorescence Mediated by Neuromodulators Adsorbed on Arrays of Oligonucleotide Rings. *Nano Lett.* **18**, 6995–7003 (2018).
7. Babu, D. & Muriana, P. M. Sensitive quantification of aflatoxin B1 in animal feeds, corn feed grain, and yellow corn meal using immunomagnetic bead-based recovery and real-time immunoquantitative-pcr. *Toxins (Basel).* **6**, 3223–3237 (2014).
8. Roxbury, D., Tu, X., Zheng, M. & Jagota, A. Recognition ability of DNA for carbon nanotubes correlates with their binding affinity. *Langmuir* **27**, 8282–8293 (2011).
9. Jena, P. V., Safaee, M. M., Heller, D. A. & Roxbury, D. DNA-Carbon Nanotube Complexation Affinity and Photoluminescence Modulation Are Independent. *ACS Appl. Mater. Interfaces* **9**, 21397−21405 (2017).
10. Yang, Y., Sharma, A., Noetinger, G., Zheng, M. & Jagota, A. Pathway-Dependent Structures of DNA-Wrapped Carbon Nanotubes: Direct Sonication vs. Surfactant/DNA Exchange. *J. Phys. Chem. C* (2020) doi:10.1021/acs.jpcc.0c00679.
11. O'Neil, M. J. *The Merck Index: An Encyclopedia of Chemicals, Drugs, and Biologicals.* (2006).
12. Bazin, I. *et al.* Impact of pH on the stability and the cross-reactivity of ochratoxin A and

citrinin. *Toxins (Basel).* **5**, 2324–2340 (2013).

13. Bol, E. K., Araujo, L., Veras, F. F. & Welke, J. E. Estimated exposure to zearalenone, ochratoxin A and aflatoxin B1 through the consume of bakery products and pasta considering effects of food processing. *Food Chem. Toxicol.* **89**, 85–91 (2016).
14. Krska, R. *et al.* Determination of molar absorptivity coefficients for major type-B trichothecenes and certification of calibrators for deoxynivalenol and nivalenol. *Anal. Bioanal. Chem.* **388**, 1215–1226 (2007).
15. Wu, X., Murphy, P., Cunnick, J. & Hendrich, S. Synthesis and characterization of deoxynivalenol glucuronide: Its comparative immunotoxicity with deoxynivalenol. *Food Chem. Toxicol.* **45**, 1846–1855 (2007).
16. McLean, M. & Dutton, M. F. Cellular interactions and metabolism of aflatoxin: An update. *Pharmacol. Ther.* **65**, 163–192 (1995).
17. Shieh, J. C. & Song, P. S. Photochemically Induced Binding of Aflatoxins to DNA and its Effects on Template Activity. *Cancer Res.* **40**, 689–695 (1980).
18. Gillen, A. J. *et al.* Templating colloidal sieves for tuning nanotube surface interactions and optical sensor responses. *J. Colloid Interface Sci.* **565**, 55–62 (2020).
